# Single cell profiling reveals strain-specific differences in myeloid inflammatory potential in the rat liver

**DOI:** 10.1101/2022.11.11.516225

**Authors:** Delaram Pouyabahar, Sai W. Chung, Olivia I. Pezzutti, Catia T. Perciani, Xinle Wang, Xue-Zhong Ma, Chao Jiang, Damra Camat, Trevor Chung, Manmeet Sekhon, Justin Manuel, Xu-Chun Chen, Ian D. McGilvray, Sonya A. MacParland, Gary D. Bader

## Abstract

Liver transplantation is currently the only treatment for end-stage liver disease and acute liver failure. Liver transplant rejection is among the most lethal complications of transplantation, and therapeutic development is limited by our lack of a comprehensive understanding of the cellular landscape of the liver. The laboratory rat (*Rattus norvegicus*), ideal in size as a model for surgical procedures, is a strong platform to study liver biology in the context of liver transplantation. Liver allograft rejection is known to be strain-specific in the rat model, although the transplantation is accepted without rejection in some strains, it leads to acute rejection in others. To shed light on the cellular landscape of the rat liver and build a foundation for strain comparison, we present a comprehensive single-cell transcriptomics map of the healthy rat liver of Lewis and Dark Agouti strains. Using a novel computational pipeline we developed to guide the detailed annotation of our rat liver atlas, we discovered that hepatic myeloid cells have strong Lewis and Dark Agouti strain-specific differences focused on inflammatory signaling pathways. We experimentally validated these strain-specific differences in myeloid inflammatory potential *in vitro* using intracellular cytokine staining. Our work provides the first examination of the multi-strain healthy rat liver by single cell transcriptomics and uncovers key insights into strain-specific differences in this valuable model animal.

**Summary:** The laboratory rat (*Rattus norvegicus*) is a standard model animal for orthotopic liver transplantation. Transplanting a liver from a Dark agouti (DA) to a Lewis (LEW) strain rat leads to transplant rejection and the reverse procedure leads to tolerance. Understanding this strain difference may help explain the cellular drivers of liver allograft rejection post-transplant. This study uses single-cell transcriptomics to better understand the complex cellular composition of the rat liver and unravels cellular and molecular sources of inter-strain hepatic variation. We generated single-cell transcriptomic maps of the livers of healthy DA and LEW rat strains and developed a novel, factor analysis-based bioinformatics pipeline to study data covariates, such as strain and batch. Using this approach, we discovered variations within hepatocyte and myeloid populations that explain how the states of these cells differ between strains in the healthy rat, which may explain why these strains respond differently to liver transplants.

## Introduction

The liver is a multitasking organ that contributes to a remarkably diverse set of processes, including metabolism and immune function. As a result of its critical roles, acute liver failure, which mainly occurs as a result of drug-induced liver injury or hepatitis A, B, and E infection, can lead to death within days ^1^. Currently, liver transplantation is the only effective treatment for patients diagnosed with acute liver failure and end-stage liver disease. Despite the inherent immunologically tolerant nature of the liver ^2,3^ and the advancements in medical strategies to treat acute liver disease ^4–6^, liver transplant patients may still be susceptible to serious complications. The development of therapeutic options to improve transplantation outcomes is limited by our incomplete understanding of the cellular landscape of the healthy and diseased liver. Further, we also need to better understand how the liver landscape differs between individual humans who have variable responses to liver treatments.

The liver is composed of multiple cell types with various complementary functions, including hepatocytes, biliary epithelial cells (cholangiocytes), stellate cells, recently recruited myeloid populations, Kupffer cells (KCs), liver sinusoidal endothelial cells (LSECs) and multiple immune cell populations. Hepatocytes make up the majority of liver volume and are involved in metabolism and drug detoxification, among other functions. Myeloid cells are found throughout the liver and can adopt proinflammatory or anti-inflammatory roles, with phenotypic characteristics of recently recruited monocytic myeloid cells and more tissue-resident Kupffer cell-like populations, respectively ^7^. Advancements in single-cell RNA sequencing technologies have equipped biologists with a powerful tool for unbiased profiling of heterogeneous tissues, such as the human ^7–12^ and the mouse ^13–16^ livers.

Mimicking human hepatic physiology in an accessible and relevant animal model for study using current single-cell-resolution genomic technologies will enable us to better understand how the liver cellular landscape functions in healthy and disease states, how this varies across individuals, and to investigate potential treatments. Laboratory rat (Rattus norvegicus) is a standard animal model used to study aspects of orthotopic liver transplantation, due to being the ideal size for microsurgery and similar postoperative immunological progression, compared to humans ^17^. Interestingly, the degree of liver allograft rejection is strain-specific in rats, and while transplantation is accepted without rejection in some strains, it leads to acute rejection in others. For instance, using the Dark Agouti (DA) strain as the donor and the Lewis (LEW) strain as the recipient, models acute liver rejection, while reversing the direction of transplantation is a model for tolerance. To understand the cellular and molecular sources that drive inter-strain variation, we need an in-depth understanding of the complex cellular landscape of the healthy rat liver.

To date, our understanding of the biology of the rat liver has been informed by technologies such as bulk RNA-seq ^18–21^, transcriptome microarrays ^22–24^, immunohistology ^25,26^, targeted qPCR ^22,25,27^ and tandem mass spectrometry ^20^. These approaches have reinforced the notion that the presence of major hepatic populations in the rat liver appears to be similar to human ^22^, however, the low resolution and targeted nature of these approaches do not allow us to have a holistic understanding of how the interaction between diverse hepatic cells shape the liver environment. Here, using single cell transcriptomics, we map the healthy rat liver at single-cell resolution and identify the molecular pathways associated with strain variations that might contribute to different transplant outcomes. A multistrain healthy map of the rat liver defines a standard reference and is essential for understanding the tolerance and rejection models in future studies.

## Results

### The cellular landscape of healthy rat liver

We generated the first multi-strain single-cell transcriptomic map of the healthy rat liver as a tool to examine the cellular complexity in this model system. Single-cell transcriptomes were generated from total liver homogenates of four 8-10 week-old healthy male rats following 2-step collagenase digestion (Figure 1A). Two livers from each of the Dark Agouti and Lewis strains were sampled and a standard scRNA-seq mapping pipeline was applied (Figure 1B). In total, 226,270 single cells were called by the 10X Genomics Cell Ranger software and 23,036 passed additional quality control filters and were included in the final map (see Methods, Figures S1-S3, Figures 1CD, Table S1). Substantial batch effects were evident while integrating the four rat samples; therefore, the Harmony ^28^ integration method was used to reduce the inter-sample technical confounding effects. After applying this batch correction, all clusters were represented by all animals (Figures 1EF). This resulted in an aggregated landscape of the healthy rat liver over all samples (Figure 1G). Cell populations were annotated using top differentially expressed (DE) marker genes ^29^ (see methods; Figure 1H, Table S2 and S3).

**Figure 1).**
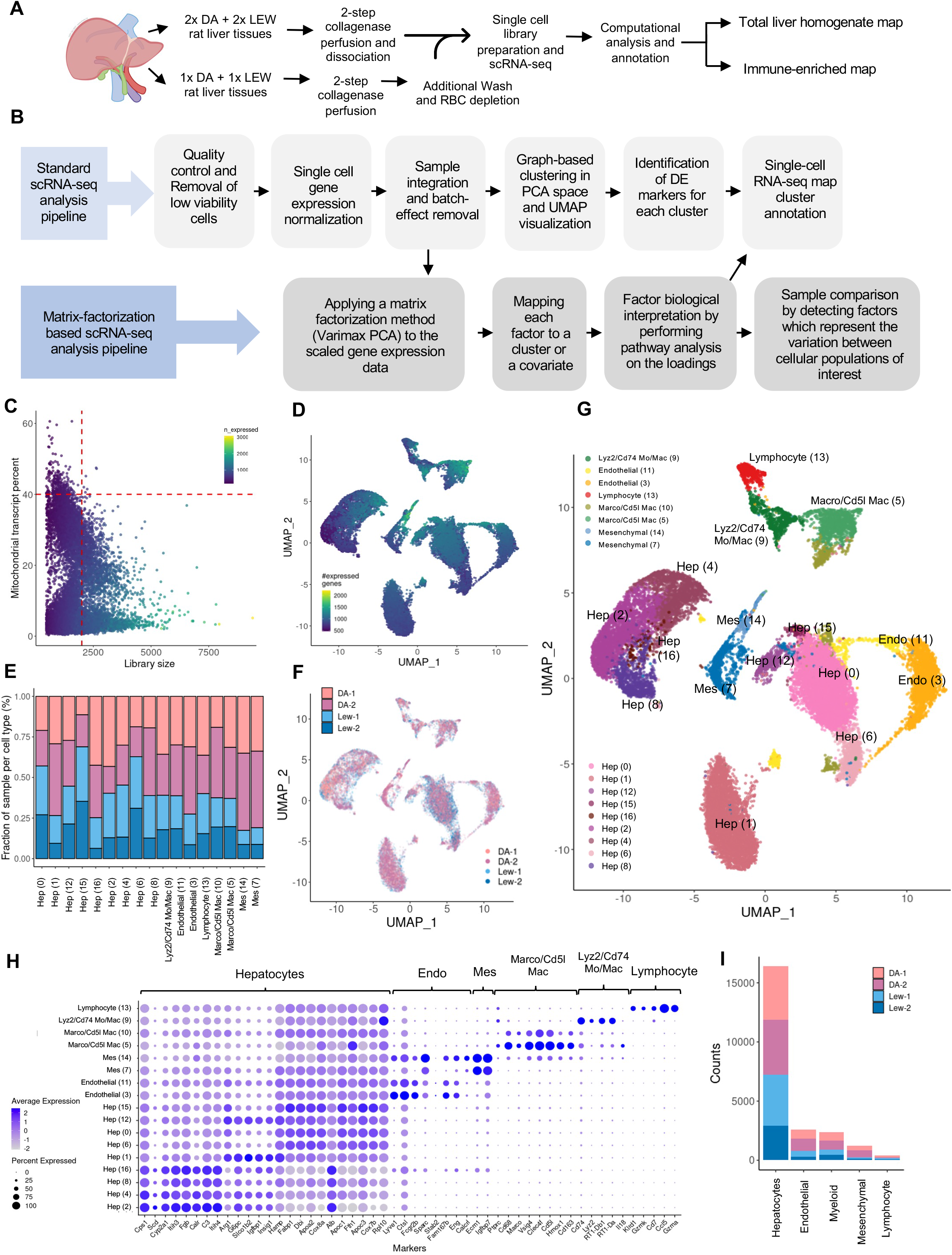
ScRNA-seq profiling of rat liver reveals 17 distinct cell populations. Overview of single-cell RNA-seq pipeline, including both the experimental and analysis workflows. Major steps of the standard and matrix factorization-based single-cell RNA-seq data analysis pipeline. (C) Viable cell selection for a Lewis rat liver sample (LEW-1) based on library size and mitochondrial transcript proportion shown as an example. High-quality cells were identified from the single-cell libraries having a minimum library size of 1500 transcripts and a maximum of 40% mitochondrial transcript proportion. (D) UMAP (uniform manifold approximation and projection for dimension reduction) plot of four rat samples including two samples from each Dark agouti (DA) and Lewis (LEW) rat strains. Cells are colored by the number of expressed genes, with lighter colors indicating higher gene counts. (E) Bar plot indicating the relative contribution of each sample in each cluster. All samples are represented in each cluster. (F) UMAP projection of cells labeled based on the input sample indicates that cells from different samples have been well-integrated and clusters represent cell-type differences rather than sample-specific variations. (G) UMAP projection of four total liver homogenate rat samples (each point represents a single cell) where cells that share similar transcriptome profiles are grouped by colors representing unsupervised clustering results. The legend indicates the unique color representing the cell-type annotation of each cluster. The cluster number is shown within the brackets. (H) Dot-plot indicating the relative expression of marker genes in each population. The size of the circle indicates the percentage of cells in each population which express the marker of interest, and the color represents the average expression value of the marker. (I) The number of cells in each major cell type population is colored by the contribution of each input sample. RBC: red blood cell, PCA: principal component analysis, DE: differentially expressed, QC: quality control, Mac: macrophage, Mo: monocyte, Endo: endothelial, Mes: mesenchymal, Hep: hepatocyte

### Hepatocytes

Hepatocytes make up the majority of the liver volume (Figure 1I) and are involved in many biological processes, such as metabolism, protein synthesis, and drug detoxification. The spatial organization of hepatocytes in hepatic lobules leads to their functional zonation from the pericentral vein to the periportal region. We identified nine hepatocyte-like clusters, comprising clusters 0, 1, 2, 4, 6, 8, 12, 15, and 16, based on their expression of hallmark hepatocyte markers without the expression of *Ptprc* (immune marker), *Sparc*, and *Lyve1* (endothelial and mesenchymal markers) (Figure 1G).

The gene expression patterns in these clusters were compared to zonated gene expression patterns characterized in laser capture microdissected periportal to pericentral regions of the healthy mouse liver lobule that were profiled by bulk RNA-seq (Figure S4) ^30^. This comparative analysis revealed expression patterns in six of the nine hepatocyte populations that significantly correlated with the mouse sinusoid zonation gene expression patterns. In this analysis, rat clusters 0, 12, and 15 correlate with central venous mouse liver layers, while rat clusters 2, 8, and 16 correlate with periportal mouse liver layers.

Periportal hepatocytes have a role in gluconeogenesis, glucagon response, and B-oxidation ^31^. Hepatocyte clusters 2, 8, and 16 which correlate with periportal mouse layers (layers 6, 7, 8, 9), show enriched expression of *Cps1*, a gene enriched in mouse periportal hepatocytes, suggesting that these hepatocyte populations may represent periportal hepatocytes (Cluster 2 top DE genes: *C3, Itih4, Alb, Itih3, Serpina3c, Fgb, Fetub, Ambp, Cps1, Hp*). Additionally, hepatocyte cluster 8 has enriched expression of *Itih4, C3*, and *Cyp2a1*, genes expressed in periportal mouse hepatocytes (Cluster 8 top DE genes: *Alb, C3, Itih4, Ambp, Itih3, Hp, Fgb, Fetub, Serpina3c, Tf, Apoe*). Hepatocyte cluster 16 also expresses periportal marker *Arg1*, a urea cycle gene specific to periportal hepatocyte function which further indicates the periportal nature of this cellular population (Cluster 16 top DE genes: *Alb, C3, Itih4, Fgp, Itih3, Ambp, Serpina3c, Fetub, Hp, F2*). Hepatocyte cluster 4 expresses *Cps1*, suggesting that this population may also be more periportal (Cluster 4 top DE genes: *Cps1, Serpina3c, Gulo, Itih3, Gjb1, Calr, Tf, Itih4, Acsl1, C3*). These results suggest that hepatocyte clusters 2, 4, 8, and 16 represent periportal hepatocytes.

Cluster 0, the most abundant hepatocyte population, is characterized by enriched expression of hepatocyte genes *Apoc1*, and *Fth1* (Cluster 0 top DE genes: *Fabp1, Rup2, Dbi, Apoc1, Fth1, Cox8a, Atp5fe, Cox7c, Cox7b, Apoc3*). Cluster 12 was identified as a hepatocyte cell population based on the enriched expression of hepatocyte genes *Dbi, Apoc3*, and *Apoc1*. This cluster also expressed *Slco1b2* ^*30*^, a gene enriched in central venous hepatocytes (Cluster 12 top DE gene: *Gadd45g, Slco1b2, Fabp1, Prp4b, Dbi, Apoc3, G6pc, Ddt, Ttr, Apoc1*). Hepatocyte cluster 15 was characterized based on the enriched expression of hepatocyte genes *Apoc1, Apoc3, Dbi, Cox7b*, and *Alb* (Cluster 15 top DE genes: *Fabp1, Apoc1, Apoc3, Dbi, Cox7b, Ddt, Cox8a, Serpina1, Apoa2, Rpl10*). These cellular populations significantly correlated with pericentral mouse zonation layers, suggesting that clusters 0, 12, and 15 may be central venous-like hepatocytes.

The remaining hepatocyte clusters did not indicate a high correlation with mouse hepatocyte layers and therefore we cannot comment on their potential zonation. Hepatocyte cluster 1 showed enriched expression of central venous hepatocyte markers *G6pc* and *Slco1b2* ^*30*^, periportal markers *Pck1, Igfbp1*, and *Insig*, and midzonal marker *Hamp* (Cluster 1 top DE genes: *Slco1b2, G6pc, Gadd45g, Akr1c1, Igfbp1, Slc38a2, Tat, Insig1, Hamp, Pck1*). Hepatocyte cluster 6 expressed periportal markers *Fabp1*, and *Apoa*, and interzonal markers *Dbi* and *Cox8a* (Cluster 6 top DE genes: *LOC689064* (Hbb-like gene), *Hba-a2, Fabp1, Dbi, Fth1, Apoa2, Cox8a, Cox7a, Cox7b, Cyb5a, Apoc3, Apoc1*). The top three markers of this cluster encoded hemoglobin subunits, suggesting that this cluster is enriched in hepatocyte-like and red blood cell-like transcripts. The expression profile of this cluster correlated with mouse erythrocytes in a mouse liver atlas (See methods, Figure S5) ^32^. Distribution of doublets was not denser in the hepatocyte clusters 1 and 6 compared to the other cell populations, rejecting the hypothesis that these clusters have a high density of erythrocyte and hepatocyte doublets (Figure S6). These findings suggest cellular populations of clusters 1 and 6 may be heterogeneous, and additional analysis will be required to confirm the identity and role of these hepatocytes.

### Mesenchymal Cells

The hepatic mesenchymal fraction includes populations such as hepatic stellate cells (HSCs), vascular smooth muscle cells, and fibroblasts ^9^. HSCs anatomically reside between sinusoidal endothelial cells and hepatocytes and are involved in vitamin A storage, extracellular matrices (ECM) synthesis, and regulation of sinusoidal circulation^33^. HSCs can exist in quiescent and activated states and are responsible for the formation of scar tissue in liver fibrosis. We identified two clusters in our map, clusters 7 and 14, which we annotated as mesenchymal-like based on DE genes including extracellular matrix proteins *Ecm1* and type III collagen alpha 1 (*Co3a1*) which are essential to the role of HSCs in extracellular matrix deposition and have been described previously as mesenchymal genes ^9,34^ (Cluster 7 top DE genes: *Ecm1, Igfbp7, Igfbp3, Col3a1, Colec11, Bgn, Angptl6, Sparc, Steap4, Pth1r*; Cluster 14 top DE genes: *Ecm1, Igfbp7, Colec11, Sparc, Igfbp4, Bgn, Col3a1, Steap4, Colec10, Prelp*) (Figure S7). Based on our correlation analyses with Dobie et al., 2019, and Andrews et al., 2022, cluster 7 is likely enriched for HSCs with active pathways in retinol storage, along with a minor fibroblast population, and cluster 14 likely contains a mixture of quiescent HSCs and fibroblasts ^34,35^ (Figure S8). These results suggest that clusters 7 and 14 are predominantly mesenchymal-like populations.

### Endothelial Cells

The hepatic endothelium consists of liver sinusoidal endothelial cells and vascular endothelium (portal and central venous endothelium). Liver sinusoidal endothelial cells (LSECs) are a specialized endothelial population that line the hepatic sinusoids and contribute to the regulation of hepatic blood pressure, nutrient uptake, waste clearance, and maintaining HSC quiescence ^36,37^. Immunohistochemical staining has characterized periportal LSECs as expressing high levels of *Cd36*, with low levels of *Lyve1* and central venous LSEC as expressing high levels of *Cd32b* and high levels of *Lyve1*, with general endothelial cells in the liver expressing high levels of *Cd31* (*Pecam*) and *Cd103* (*Eng*) ^38,39^. We identified two populations of *Ptprc*^-^ cells (clusters 3 and 11) which were annotated as endothelial-enriched based on the expression of *Calcrl* and *Ramp2*, involved in adrenomedullin signaling ^40^. Cluster 3, the most abundant endothelial cell population, was characterized by enriched expression of *Lyve1, Fcgr2b, Sparc*, and *Stab2* with little expression of *Vwf* (Cluster 3 top DE genes: *Lyve1, Ctsl, Fcgr2b, Fam167b, Kdr, Id3, Eng, Sparc, Ifi27, Igfbp7*) (Figure S9). Both clusters 3 and 11 are similar to human LSECs and also mouse sinusoidal, inflammatory, and cycling endothelial populations (Figures S10 and S5). The enriched gene expression in the cluster 11 endothelial cell population includes both central venous LSEC genes *Lyve1, Fcgr2b* (protein alias CD32), *Ctsl*, and non-LSEC portal endothelial genes *Fam167b, Id3, Eng* (Cluster 11 top DE genes: *Lyve1, Fcgr2b, Ctsl, Fam167b, Bmp2, Id3, Kdr, Gstp1, Srgn, Eng*). Furthermore, these clusters did not express high levels of known zonated endothelial genes such as *Rspo3* ^41^ *and Clec4g* and both clusters expressed high levels of *Fcgr2b* (known to be CV LSECs ^39^) and *Aqp1* (known to be periportal ^31^) (Figure S9). Therefore, the zonation of these populations could not be resolved. Additional enrichment of endothelial cells in the future will increase the resolution of this population.

### Myeloid cells

There are more resident myeloid cells in the liver than in any other organ in the body ^42^. Hepatic myeloid cells are critical mediators of the tolerogenic environment of the liver in the steady state. Tissue-resident myeloid cells exhibit immense plasticity and can perform a variety of functions and phenotypes. Depending on the local immune microenvironment and the external stimulus, bone marrow-derived monocytes can be recruited to the liver, where they participate in both liver injury and repair ^43,44^. Following transplantation, the hepatic myeloid pool is vastly reorganized and the recipient-derived myeloid cells replace donor hepatic myeloid cells in the transplanted liver(∼99%)^45^. These recruited monocyte-derived myeloid cells can then transition to attain a tissue-resident and pro-repair phenotypes ^46,47,48^). Our analysis revealed multiple clusters of *Cd68*^+^ myeloid-enriched cells. *Cd68*^+^ myeloid clusters 5 and 10 were characterized by enriched expression of *Marco, Vsig4, Cd5l, Cd163*, and *Hmox1* (Cluster 5 top DE genes: *Ccl6, C1qa, C1qb, C1qc, Tmsb4x, Cd5l, Clec4f, Aif1, Marco, Tyrobp*; Cluster 10 top DE genes: *C1qb, C1q1, Clec4f, Slfn4, Marco, Ccl6, C1qc, Cd5l, Psap, Npc2*) (Figure S11). These clusters appear to be more Kupffer cell-like due to the expression of key genes (*Marco, Cd5l, Clec4f*) which have been previously described to annotate more tissue-resident myeloid populations ^49^. Specifically, *Vsig4* is a co-inhibitory ligand that has a hepatoprotective role in maintaining the intrahepatic tolerance required to suppress triggered immune responses ^46,47^ and has been shown to be highly expressed in murine KC ^46,47^ as well as being a core KC gene in pig and macaque KCs ^16^. These findings may suggest a tolerogenic role of *Marco*^+^*Cd5l*^*+*^*Cd68*^*+*^ cells.

Our analysis of cluster 9 revealed a mixed cluster of *Ptprc+* immune cells enriched for *Cd68*^*+*^ myeloid cells. Cluster 9 was characterized by enriched expression of macrophage/monocyte markers *Cd68, Cd74, Lyz2*, and MHC class1-related genes (Cluster 9 top DE genes: *Cd74, Tmsb10, RT1-Db1, RT1-Da, Cst3, RT1-Bb, Plac8, RT1-Ba, Lyz2, Tyrobp*), without expression of *Vsig4* and *Marco* suggesting that cluster 9 is enriched for recently-recruited macrophage/monocyte populations while also containing additional populations such as T cells (*Cd3e*), cDC1s (*Clec9a, Xcr1, Batf3, Irf8*) (Figure S11) and cDC2s (*Clec10a+, Irf4, Sirpa*) and pDCs (*Siglech*).

### Varimax PCA analysis uncovers biological sources of variation between the rat strains

We mapped two rat strains, Dark Agouti and Lewis, as these are important for the rat model of orthotopic liver transplants. To better understand strain-specific differences in our map, we applied varimax principal component analysis (PCA) ^50,51,52^ a matrix factorization method, to separate Dark Agouti and Lewis signals (principal components, or factors) in the data from other signals for further interpretation (Figure 1B, Figure 2, Table S4). To identify factors that can explain strain-specific differences, we used a random forest to predict strain labels from the factors identified per cell and discovered the factors most important to the strain label classification (Figure 2A). We also identified principal components that explain cell type signals using correlation analysis (Figure 2B, Figure S12). The resulting factors were interpreted using pathway and gene set enrichment analysis (see: Methods). Using this approach, two main strain-specific factors (varimax PC5 and 15) were identified (Figure 2A, Figure S13). The strongest strain-specific signal is observed with varimax PC5, which affects all cells in the data (Figure 2CD). Genes with the strongest association with this factor are hepatocyte markers (*Apoc1, Fabp1*, Cytochrome p450 genes), suggesting that this factor mainly represents strain variations within the hepatocyte populations (Figure 2E). The global association of this factor with all cells is likely a cell-dissociation procedure artifact caused by fragile hepatocytes leaking RNA into the cell homogenate before sequencing (Figure S14) ^34^. Dark Agouti strain-associated genes in this factor are enriched in nuclear receptors, such as *Hnf4a, Pparg, Esr1* (Table S5), and pathways such as lipid, cholesterol, and xenobiotic metabolism (Figure S15). *Pparg* promotes de novo lipogenesis and fat accumulation in hepatocytes ^53,54^. The second-strongest strain-specific signal is varimax PC15, which is mainly associated with myeloid populations of both rat strains (Figure 2FG), as confirmed by the genes with the strongest association with this factor (Figure 2H), the expression pattern of *Marco, Visg4, Cd68*, and *Lyz2* marker genes (Figure 2I) and correlation with myeloid cells in our map (Figure 2J).

**Figure 2).**
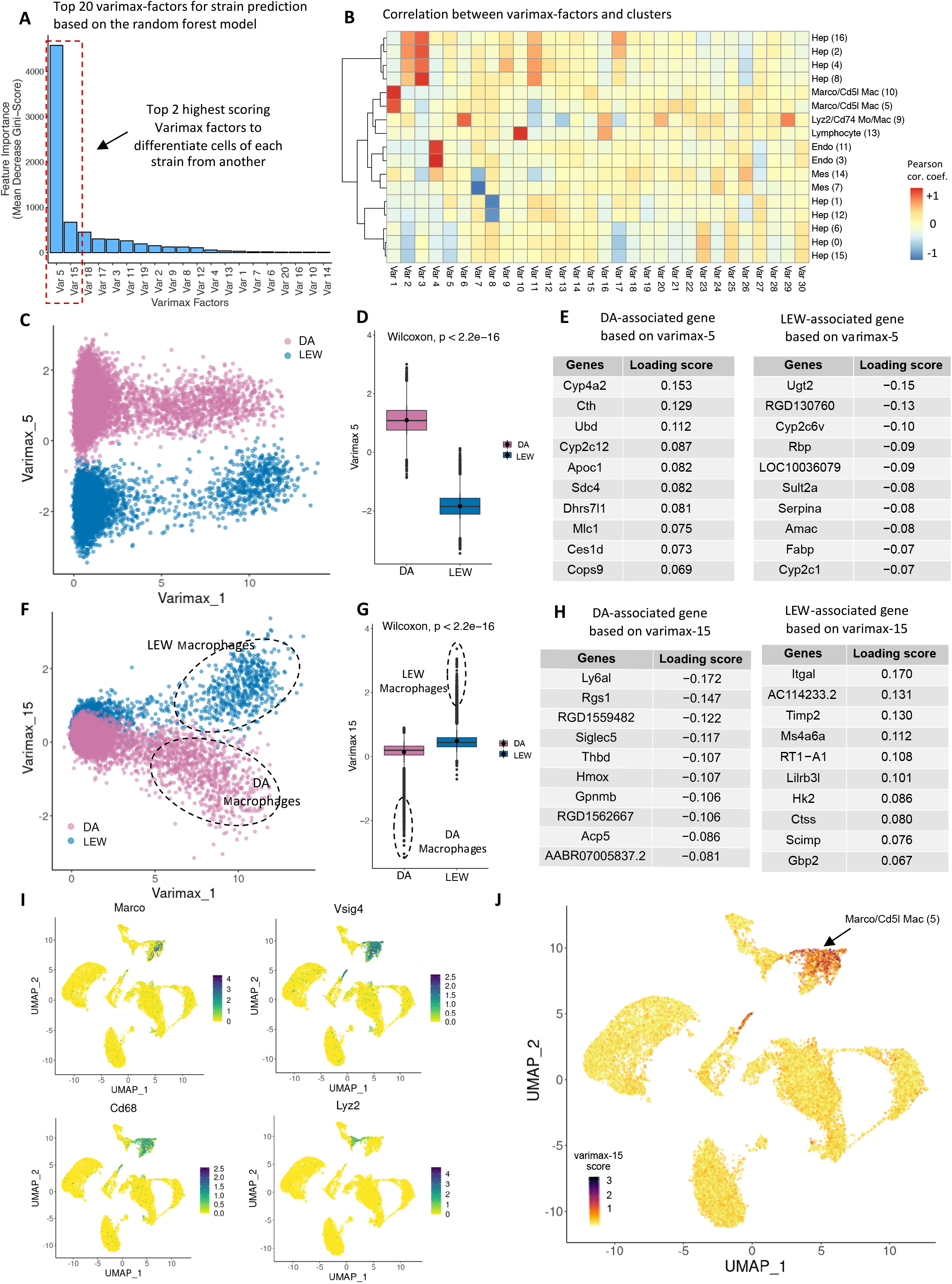
Varimax-PCs capture rat hepatic cell identity signatures and strain-specific differences. (A) Bar plot representing the feature importance scores (mean decrease in Gini impurity score) of the top 20 features (varimax factors) of the random forest model trained to predict the strain attributes of the rat hepatic cells. Varimax PC5 and 15 are the most informative features to differentiate cells of each strain from another, which indicates the two factors have captured strain-related variations within the map. (B) A correlation heatmap between the average gene expression of each cluster and the loading scores of varimax factors (capturing the contribution of all genes to a factor). Columns are varimax factors and rows are cell populations. Each cell-type cluster is defined by key marker genes and dark red or blue indicates that the expression of a marker gene set is positively or negatively correlated, respectively, with a particular varimax factor. High absolute correlation value indicates a match between a varimax factor and a cell-type cluster. (C) Projection of cells over Varimax factors 1 and 5 indicates that the cells from each strain form distinct clusters over varimax-5. (D) Boxplot indicating the distribution of Varimax-5 score over each strain. Cells from DA and LEW strains represent significantly different varimax-5 scores (Wilcoxon-test p-value < 2.2e-16), indicating that varimax-5 has captured strain differences. (E) The top 10 genes on the top (left table) and bottom (right table) of the Varimax-5 loading list mainly contain known hepatocyte markers, indicating that Varimax-5 has captured hepatocyte-specific strain differences. Genes with high positive scores (left table) are associated with the DA strain and genes indicating negative loading scores (right table) are LEW-related. The absolute loading scores indicate the contribution of each gene to the corresponding factor. (F) Projection of cells over Varimax-1 and 15 indicates that a population of cells from each strain (dotted lines) form distinct clusters over varimax-15. Annotation of the selected cells indicates that they are mainly from the *Marco*+ myeloid cluster-5. (G) Boxplot indicating the distribution of hepatic cells based on strain over Varimax-15. (Wilcoxon-test p-value < 2.2e-16). The outlier data points (dotted lines) are mainly myeloid cells. (H) The top 10 genes with positive (right table) and negative (left table) Varimax-15 loading scores are immune-response related. Genes with positive scores (right table) are associated with the LEW strain and genes indicating negative loading values (left table) are DA-related. The absolute loading scores indicate the contribution of each gene to the corresponding factor. (I) Expression pattern of known myeloid marker genes *Marco, Vsig4, Cd68*, and *Lyz2* on the UMAP of all total liver homogenate cells. Dark green represents high expression values. The distribution of general myeloid markers (*Cd68, Vsig4*) and non-Inflammatory myeloid marker (*Marco*) is consistent with the varimax-15 distribution (Figure 2J). (J) The UMAP projection of cells colored based on varimax-15 score shows the enrichment of varimax-15 over *Marco*+ myeloid population (cluster 5). Darker colors represent higher values of varimax-15 scores. Cor. coef.: correlation coefficient, Var: Varimax principal component (PC). Varimax PCs are referred to as PCs within the main text.

Comparing the expression level of the top varimax PC15 genes in the myeloid cells of the two strains confirms the strain-specificity of this factor (Figure 3ABC). Pathway analysis identified higher activation of lymphocyte-mediated immune responses, lymphocyte migration and chemotaxis, response to interferon and allograft rejection pathways in LEW compared to DA *Marco*-enriched myeloid cells (Cluster 5) (Figure 3D, Table S6). This factor is enriched in myeloid and T-cell differentiation transcription factors (TFs). The Lewis-enriched TFs are *Irf8, Spi1, Pou5f1, Stat4*, and *Stat5a* which are mostly inflammatory process-associated genes present in chronic diseases like rheumatoid arthritis. For example, *Irf8/Spi1* (PU.1) are known to work cooperatively to shape the chromatin landscape to polarize the macrophage for inflammatory responses whilst *Stat4* deficiency leads to repolarization towards alternatively activated macrophages ^55–57^. The DA-specific TFs *Nucks1, Runx1, Mitf*, and *Gata1* have more diverse functions ^56,58–60^ (Figure 3E, Figure S16, Table S5). For example, *Nucks1* and *Runx1* are implicated in immunomodulation, while *Gata1* and *Mitf* are associated with cell fate and differentiation ^61–64^.

**Figure 3).**
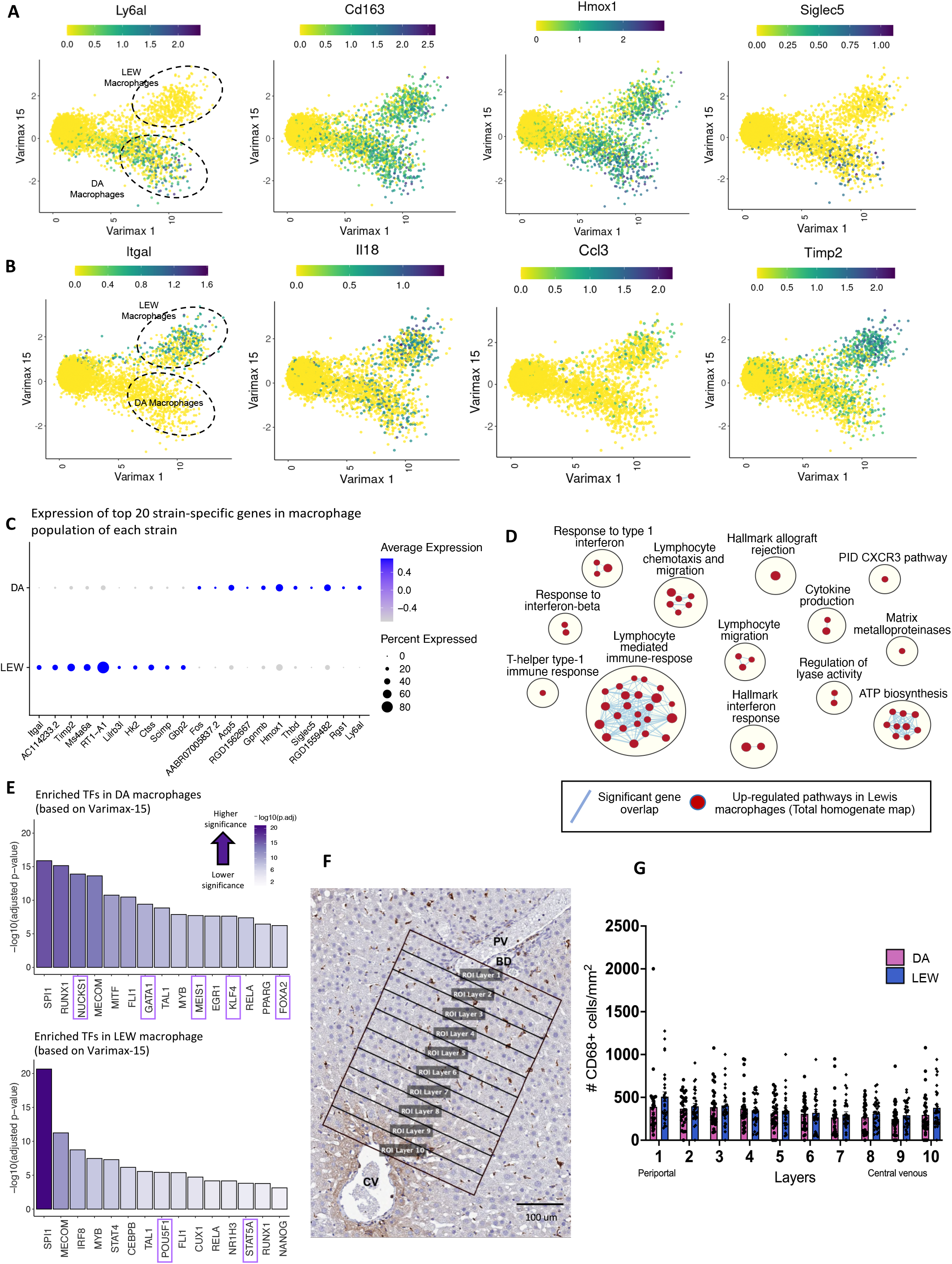
Strain-specific differences are found in intrahepatic myeloid cells. (A) Expression pattern of the top DA-enriched genes (*Ly6al, Cd163, Hmox1, Siglec5*) over PC15 and 1. LEW and DA myeloid cells have been marked with dotted circles. Dark green indicates higher expression values. Comparison with Figure 2F confirms that the selected genes have higher expression in the DA strain compared to LEW. (B) Expression pattern of the top LEW-enriched genes (*Itgal, Il18, Ccl3, Timp2*) over Varimax-15 and 1. Comparison with Figure 2F confirms that the selected genes have higher expression in the LEW strain compared to DA. (C) Dot-plot indicating the relative expression of strain-related genes within the myeloid fraction (clusters 5, 10, 9) of each strain. The top 10 genes with positive (LEW-associated) and negative (DA-associated) Varimax-15 loading scores have been selected. The size of the circle indicates the percentage of cells in each population expressing the marker, and its color shows the average expression value. (D) Pathway enrichment analysis using GSEA (Gene Set Enrichment Analysis) to examine active cellular pathways in LEW vs. DA myeloid cells based on varimax-15 loadings visualized as an enrichment map. Each circle represents a gene ontology (GO) biological process term. The size of the circles represents the number of genes in that pathway and blue lines indicate significant gene overlap. Since PC15 is positively correlated with the LEW strain and negatively correlated with DA, red circles represent activated pathways in LEW and blue indicates up-regulated pathways in DA. No pathway was significantly upregulated in DA. (E) Transcription factor (TF) binding site-based gene-set enrichment analysis using gProfiler on the ChEA ChIP-Seq database identifies TFs which may be activated in DA and LEW myeloid cells. TFs are sorted based on their enrichment significance calculated as –log10(adjusted p-value). Dark purple indicates higher significance. Purple boxes highlight TFs which are uniquely enriched in that strain. (F) Representative spatial distributions of CD68+ cells in the rat liver lobule. Rectangular layers 350um wide were drawn from the portal tract (layer 1) to the central vein (layer 10) region. Digital images were scanned at 20X magnification. Scale bar represents 100um. Each rectangular layer is referred to as a region of interest (ROI). (G) Quantification of CD68+ cell densities (#CD68+ cells/layer mm^2^) in the liver lobule for DA and LEW rats. 30 ROIs were assessed per strain across three animals. Statistical significance was evaluated using the non-parametric Mann-Whitney analysis. No significant strain-specific differences in the spatial distribution of CD68+ cells were noted. p<0.05. ROI: region of interest, BD: bile duct, CV: central vein, PV: portal vein.

These results suggest that the baseline hepatic microenvironment in the Lewis rat is more pro-inflammatory compared to the Dark Agouti strain, and highlights myeloid cells as the potential key drivers of enriched inflammatory pathway activation in myeloid cells of Lewis rats. We then asked if myeloid cell frequency in the DA versus LEW livers is distinct, and also if the more inflammatory status of the LEW myeloid cells might be accompanied by a higher intrahepatic frequency. Alterations in cell-type frequencies in scRNA-seq data are highly affected by the amount of exposed vasculature that can be cannulated in the rat which can impact sample-specific perfusion efficiency; therefore, we relied on immunohistochemistry to compare the frequency of CD68^+^ cells between the two strains. Quantification of histology staining using a publicly available QuPath-based image analysis protocol ^65^ indicated no significant difference between the frequency of LEW and DA CD68^+^ cells either in the liver in total or within individual hepatic layers (Figure 3FG). These results suggest that the variation in inflammatory potential is not caused by differences in the frequency of intrahepatic CD68^+^ cells.

### Immune enrichment depicts rat lymphocyte and myeloid populations with higher resolution

Our total liver homogenate (TLH) map contained hepatocyte-derived ambient RNA, as expected ^34^ (Figure S14), that interfered with immune cell marker identification and resulting immune cell annotation. To provide a more detailed resource of rat hepatic immune cells, two additional immune-enriched samples were mapped (Figure 4A). These samples underwent additional washing steps and red blood cell depletion to reduce the hepatocyte-released ambient RNA (Figure S17). The percentage of cells annotated as hepatocytes decreased from 71.14% in the total liver homogenate map to 49.11% in the immune-enriched map. The general immune cell marker *Ptprc* was expressed in 24% of the total cells in the immune-enriched map compared to 4% within the initial map (Figure 4BC). Unfortunately, the total liver homogenate and immune-enriched maps could not be integrated computationally, presumably due to the technical differences in their generation (Figure S18). The varimax-based pipeline was also ineffective to deconvolute the sources of variation in the merged dataset of both sets of samples (Figure S19). Consequently, the immune-enriched samples were analyzed separately. In total, 3830 (1161+2669) single cells from the DA and LEW samples were integrated into the immune-enriched map after quality control (see: Methods, Figures S20, and S21). Similar to the total liver homogenate map, the immune-enriched samples were batch-corrected and the final clusters represent cells from both DA and LEW rats (Figure 4DE). The clusters were annotated based on the same approaches used for the initial samples (Extended results, Table S7).

**Figure 4).**
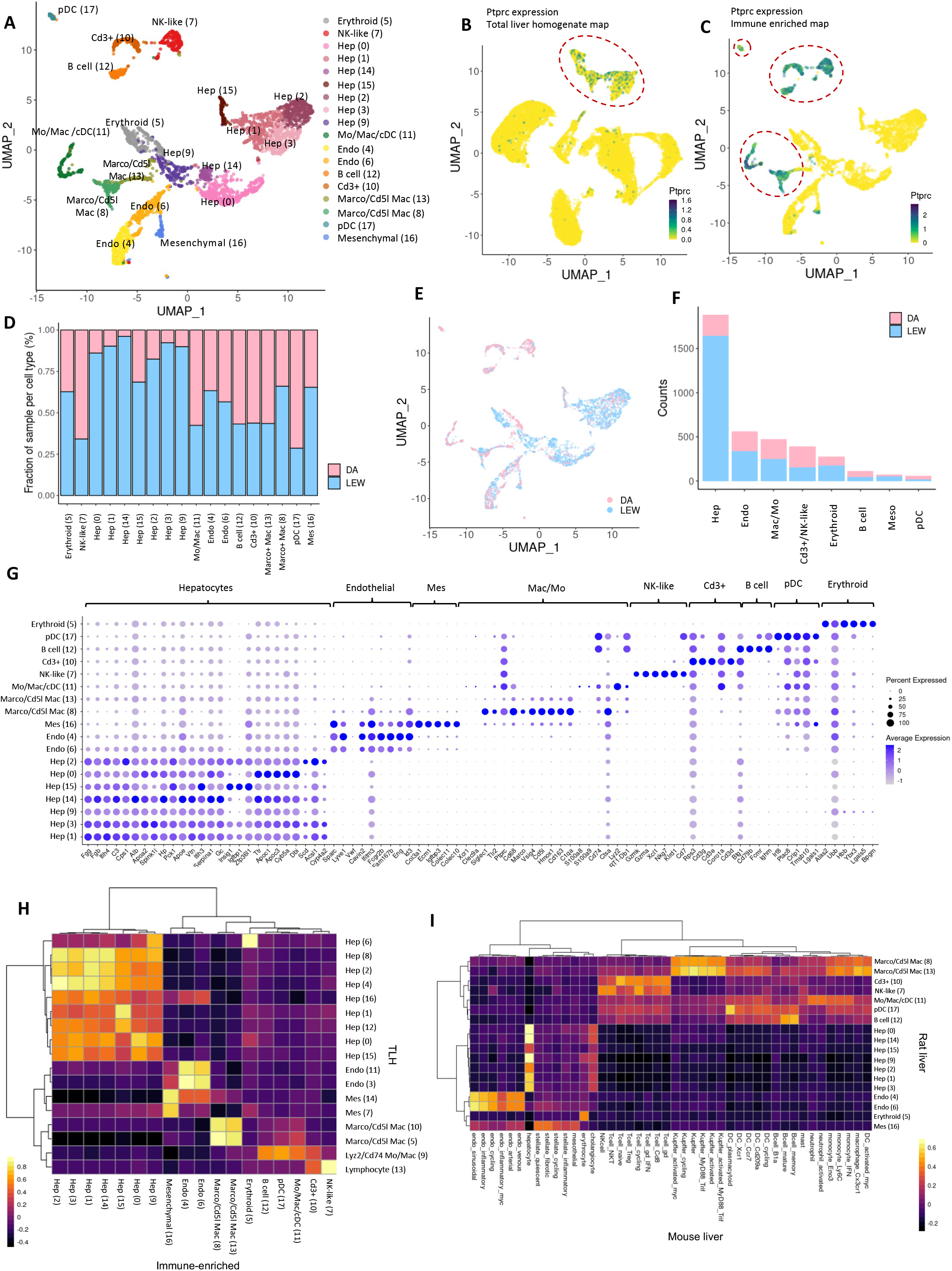
An immune-enriched scRNA-seq rat liver map provides a higher resolution view of lymphocytes and myeloid populations. (A) UMAP projection of immune-enriched samples where cells that share similar transcriptome profiles are grouped by colors representing unsupervised cell clustering results. As opposed to the total liver homogenate map, B cells and plasmacytoid dendritic cells (pDCs) have been well-captured in the immune-enriched map, and *Cd3*+ and NK-like cells form distinct populations. The legend indicates the unique color representing the cell-type annotation of each cluster. The cluster number is shown within the brackets. (B) Expression distribution of *Ptprc*, a general immune cells marker, over the UMAP projection of total liver homogenate cells. (C) *Ptprc* expression over the UMAP projection of immune-enriched map’s cells. Comparison with Figure 4B indicates that the immune-enriched map provides a better representation of the immune population compared to the total liver homogenate map. (D) Bar plot indicating the relative contribution of input samples to each cell population. Both samples have been represented in each of the clusters (cell types). (E) UMAP projection of immune-enriched cells labeled based on the input sample indicates that cells from different samples have been well-integrated and clusters represent cell-type differences rather than sample-specific variations. (F) The number of cells in each major population colored by the contribution of each input sample. (G) Dot-plot indicating the relative expression of marker genes in each population. The size of the circle indicates the percentage of cells in each population which express the marker of interest. (H) Comparison of total liver homogenate and immune-enriched rat liver maps. Rows and columns of the correlation heatmap represent the clusters within total liver homogenate and immune-enriched maps, respectively. The color of the heatmap cells indicates Pearson correlation values between the cluster average gene expressions. The top 500 highly variable genes in each map were used for the correlation calculation. (I) Comparison of rat healthy liver immune-enriched map) and mouse healthy liver map ^32^. Rows and columns of the correlation heatmap represent the rat and mouse clusters, respectively. The color of the heatmap cells indicates Pearson correlation values between the cluster average gene expressions. The one-to-one orthologs in the top 2000 highly variable genes of the two maps were used for correlation calculation (see methods). The comparison indicates a high consistency between the gene expression pattern of hepatic cell types between rats and mice.

The immune-enriched map captures a more diverse set of liver resident immune cells (Figure 4AF), enabling a more detailed description of these cell populations (Figure 4G) compared to the total liver homogenate map. A comparison of the total liver homogenate and immune-enriched maps using correlation analysis confirmed that the immune-enriched map provides a higher resolution of lymphocytes and myeloid cells (Figure 4H). As a refinement to the immune annotations in the total liver homogenate map, individual populations of *Cd3*^+^ T cells (clusters 10), Natural Killer(NK)-like cells (cluster 7), B cells (cluster 12), plasmacytoid dendritic cells (cluster 17) (Figure S22), were identified (described below) in the immune-enriched map (Figure 4H). Cluster 10 was characterized by enriched expression of *Cd3*^+^ T cell markers (*Cd3g, Cd3e, Cd3d, Coro1a*) (Figure S23). Solidifying the annotation of this cluster, comparison with the human liver map (Figure S24) indicates a high similarity between cluster 10 gene expression and human T cells. Cluster 12 identified a subset of cells enriched for B cell genes *Cd19, Ms4a1 (Cd20), Ighm, Cd74, Cd79b*, and *Fcmr*, with no expression of *Ighd* or *Ighg*, suggesting that this cluster might be *Cd19*+*Cd20*+*IgM+IgD-* immature B cells ^66^(Figure S25). Correlation heatmaps (Figure 4I, Figure S24) indicated high gene expression similarity with the mouse ^32^ and human B cell populations, suggesting that this is a B cell population. Enriched gene expression in cluster 17 correlates with both monocyte-like macrophages (*Cd74* and *Tyrobp*), and plasmacytoid dendritic cells (*Siglech* ^*16*^, *Ptprcap* ^*67*^, and *Ptcra* ^67^)(Figure S26). When comparing the expression of this cluster to the mouse liver cell atlas, we see a high correlation with plasmacytoid dendritic cells (Figure 4I), suggesting that the predominant cellular population of this cluster might be pDC-enriched ^32^ (Top DE genes: *Irf8, Siglech* ^*16*^, *Tyrobp, Cd74, Plac8, Ifi30, Crip1, Tmsb10, Lgals1*). The DE genes in cluster 7 included *Tbx21* [aka T-bet], *Prf1, Nkg*7, and *Ccl5, Cd8a, Gzmk, Klrd1*, and *Cd7*, with no expression o*f Cd3d*, which suggests an activated NK-like population (Figure S27). The expression of top genes in this cluster correlated with NK cell population in the mouse dataset (Figure 4I) reinforcing that this cluster is an NK enriched cluster. The *Ptprc*^+^ clusters of the immune-enriched map were subclustered for further evaluation (Figures S28-S33, Table S8). Upon subclustering of the *Ptprc*^+^ clusters, cDCs (cDC1: *Clec9a, Xcr1, Batf3*; cDC2: *Clec10a, Tmem176b* ^*68–70*^) which were mixed with other immune populations in the total liver homogenate map, formed a separate subcluster (See: extended results, Figures S22 and S33).

Comparison of previously published mouse and human liver data with the rat single-cell atlas indicates high consistency of the majority of the cell types between these species (Figure 4I, Figure S24). In the immune-enriched map analysis, we also attempted to determine if we could capture the strain-specific factors identified based on the total liver homogenate map (Figures S34 and S35). The top 10 genes which represented each factor were selected and their enrichment pattern within the immune-enriched map was evaluated. Consistent with our predictions based on the total liver homogenate map, both varimax PC5 and 15 signatures indicated strain differences within the immune-enriched samples and were specific to hepatocyte and myeloid populations respectively.

In summary, the immune-enriched map represents a more detailed evaluation of the immune landscape of the healthy rat liver and provides additional information on B cells, DCs, *Cd3*^+^ T cells, and NK-like populations in comparison with the total liver homogenate map.

### Validation of computationally inferred strain-specific inflammatory differences with orthogonal approaches

To functionally validate the computationally inferred strain-specific differences in the inflammatory potential of hepatic myeloid cells, we performed *ex vivo* LPS stimulations followed by intracellular cytokine staining. In these assays, we stimulated fresh non-parenchymal cells from flushed, enzymatically dissociated livers from both LEW and DA rats *in vitro* and examined cytokine secretion from tissue-resident myeloid cells *via* intracellular cytokine staining for TNFα (See methods, Figure S36). We found a higher frequency of LEW intrahepatic myeloid cells (CD45^+^CD68^+^CD11b^+^) secreting TNFα in response to LPS stimulation compared to DA liver resident-myeloid cells (Figure 5ABC, 4B, and 4D), which suggests, in agreement with our transcriptomic analysis (Figure 3C-3E), that the inflammatory potential of hepatic myeloid cells in LEW positive (% TNFα positive = 35.25 ± 3.18 (SEM)) is higher than that of DA (%TNFα positive =22.25 ±1.45 (SEM)). In addition, this finding is consistent with previous studies that show DA myeloid cells exhibit less inflammatory characteristics and a lower ability to stimulate T cell proliferation than LEW myeloid cells in mixed lymphocyte reaction assays in the context of alloreactivity ^71^. However, despite the overall higher TNFα response in LEW myeloid cells, the overall difference in the TNFα^+^ mean fluorescence intensity (MFI) did not reach significance (Figure 5D). In the computational analysis, the higher inflammatory potential in LEW liver myeloid cells was accompanied by a relatively enriched expression of *Itgal* transcripts (Figure 2H). ITGAL (CD11a) encodes part of LFA-1, Lymphocyte function-associated antigen 1, the expression of which is associated with inflammation and several autoimmune conditions ^72^. Our computational findings that ITGAL transcript expression is higher in LEW myeloid cells, is in keeping with our assertion of higher myeloid inflammatory potential in the hepatic myeloid cells of these animals. Further examination of intracellular cytokine data post-stimulation revealed that the strain-specific proinflammatory differences rested primarily within ITGAL^+^ myeloid cells, reflecting transcriptomic analysis that LEW possesses a more inflammatory CD68^+^ CD11b^+^ myeloid population, compared to the DA rat (Figures 5EF, 4CEF). We also observe a lack of strain-specific differences in the frequency of either CD68^+^/ITGAL^+^ or CD68^+^ myeloid cells in the flow cytometry analysis (Figure S37).

**Figure 5).**
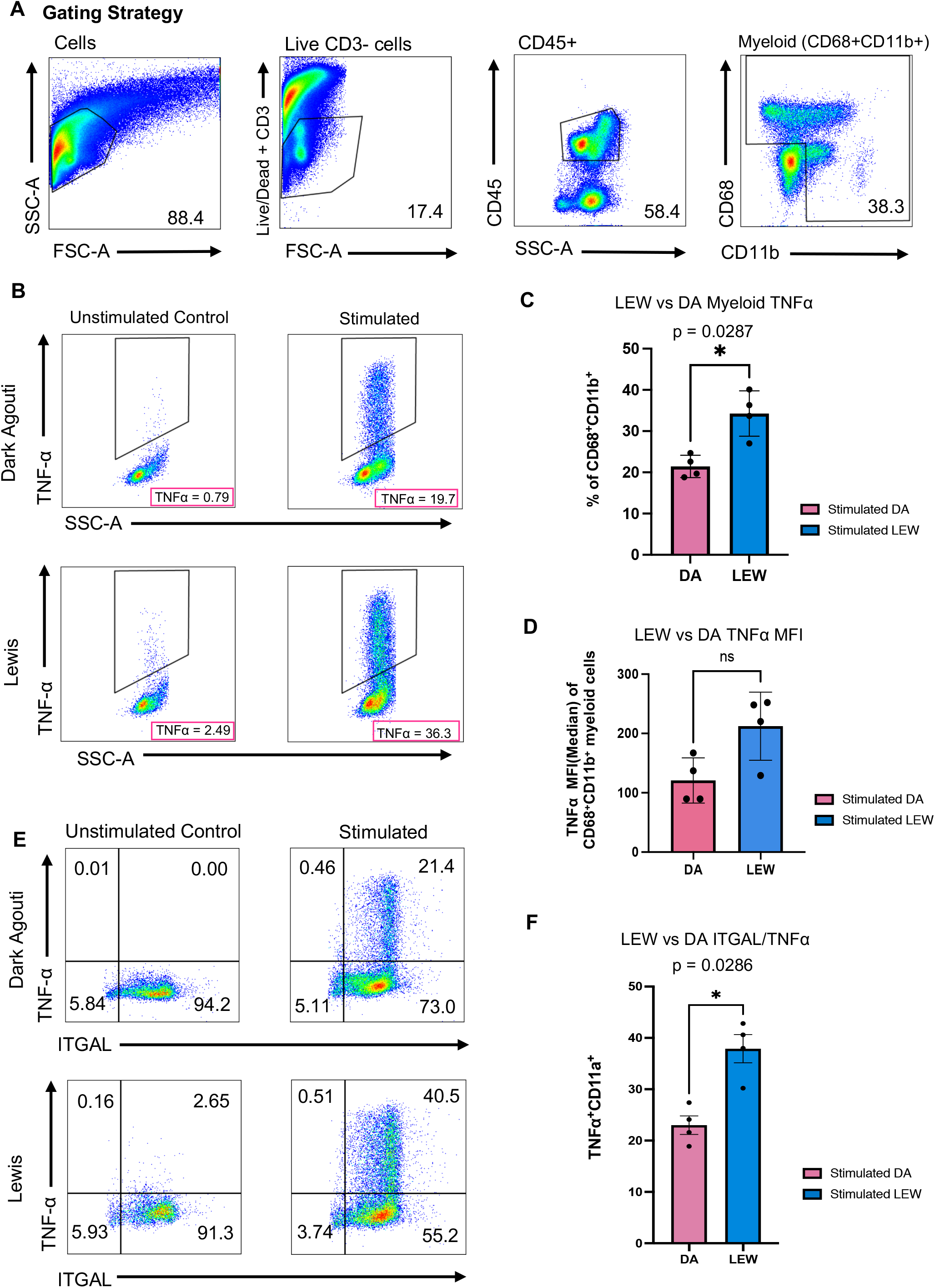
The inflammatory potential of myeloid cells found in LEW rats is greater than that found in DA rats. Myeloid cell inflammatory potential was evaluated after lipopolysaccharide (LPS) stimulation of freshly isolated liver-resident non-parenchymal cells. LPS-induced TNFα secretion was measured via intracellular cytokine staining. The non-parenchymal liver cell (NPC) dissociate was obtained via a gentle enzymatic perfusion process and differential centrifugation. The resulting cells were plated in 12 well plates for 3.5 hours before being stimulated for 6 hours under a concentration of 1ng/mL of LPS in the presence of 1:1000 concentration of Monensin and Brefeldin. (A) Flow cytometry plots showing the gating strategy for CD68^+^CD11b^+^ myeloid cells. (B) Representative flow cytometry plots depicting percentage of TNFα^+^ secreting CD68+CD11b+ myeloid cells in the unstimulated control and stimulated conditions of Dark Agouti and Lewis myeloid cells. (C) Summary graphs of Lewis versus Dark Agouti total TNFα as a percentage of CD68^+^CD11b^+^ myeloid, (D) and of the TNFα’s mean fluorescence intensity (MFI) of Lewis vs Dark Agouti myeloid cells. (E) Representative flow cytometry plot of TNFα secretion patterns based on ITGAL subpopulations. (F) Summary graph of TNFα and ITGAL expressing CD68^+^CD11b^+^ myeloid subpopulations. Plotted are the values from all four experimental replicates (n=4). Statistical significance was determined using a two-tailed Mann-Whitney test.

## Discussion

Here, we used single-cell transcriptomics to provide the first multi-strain atlas of the healthy rat liver. This study also identifies cellular and molecular sources which contribute to strain differences and highlights the role of myeloid cells in establishing higher baseline inflammation-response levels in the LEW compared to the DA strain. Examination of these strain-specific differences may help better understand and develop treatments for liver transplant rejection. Moreover, our findings indicate the importance of in-depth characterization of myeloid subpopulations in future studies on liver transplant rejection and tolerance. This study was limited by the well-known challenge of hepatic tissue dissociation in which there is a trade-off between the loss of dissociation-sensitive hepatocytes and levels of ambient RNA contamination coming from damaged cells. We addressed these challenges using computational approaches, including varimax PCA, which enabled us to better extract the cellular signal within the data. Cell enrichment protocol optimization and additional single cell genomic technologies could help improve cell type identification. For example, simultaneous capture of the transcriptome and protein expression using Cellular Indexing of Transcriptomes and Epitopes by Sequencing (CITE-seq) could help identify certain cell populations like immune cells, single-nucleus RNA-seq which is less affected by cell dissociation effects could be performed to define hepatocyte subtypes and zonation patterns, for example ^34,73^ inclusion of more samples could help identify less frequent cell populations such as cholangiocytes. The proportion of various cell-types is highly influenced by the dissociation steps and capturing efficiency differs between cell populations. Thus, the relative fraction of cell-types in our map is not necessarily equal to their actual frequency within the original rat liver tissue. Single-nucleus RNA-seq or spatial transcriptomics methods are better suited to study cell frequency.

We find that myeloid cells from LEW livers have higher inflammatory potential than myeloid cells from DA livers. We speculate that there is a baseline higher inflammatory status in LEW rats that is driving the strain-specific differences in these animals. Future studies are needed to study why this occurs, including studying the effects of genetic changes that may explain this difference, and how these affect the rat liver cellular environment during liver transplantation. Our data strengthens the notion that reprogramming hepatic myeloid cells may be an attractive way to target to modulate rat liver inflammation ^74^.

Taken together, our single-cell transcriptional map of the rat liver microenvironment adds to our understanding of the cellular basis of the rat liver function and uncovers hepatic strain differences within this animal system. It also suggests ways to investigate new therapeutic options in this model animal which can be ultimately transferred to humans to treat and prevent hepatic inflammation.

## Methods

### Biological Specimens

Whole livers were acquired from male Dark Agouti (DA) and Lewis (LEW) rats between 8-10 weeks of age. All experimental procedures followed principles and guidelines for the care and use of animals established by the Animal Resources Centre (ARC) at the University Health Network and are in accordance with the guidelines of the Canadian Council of Animal Care. Rat experiments were performed at the Toronto General Research Institute, Toronto, ON, Canada under the approval of the Institutional Committee on Animal Bioethics and Care (AUP 5840). All surgery was performed under isofluorane anesthesia, and all efforts were made to minimize animal suffering.

### Rat liver tissue dissociation

The rats were anesthetized with 5% isoflurane with an anesthetic apparatus, and the abdominal cavity was opened. Heparin (LEO) is directly injected into the Inferior Vena Cava (IVC). The IVC is then cannulated with a 20G cannulae (Braun) and flushed with a 4°C 1X Krebs solution with 0.01mM EGTA at a rate of 10mL/min for 5 minutes and then a 37°C warm Krebs solution with 1.35 mM of CaCl_2_ and 0.04% Collagenase (Milipore Sigma) from *H. Clos* at a rate of 5mL/min for 12 minutes. The digested liver is excised into HBSS, and the Glisson capsule is shaken and opened to release the single-cell suspension. This total liver homogenate is then filtered by a 40um mesh filter (Falcon).

### 10x sample processing and cDNA library preparation

Samples were prepared as outlined by the 10x Genomics Single Cell 3’ v2 (TLH samples) and v3 (immune-enriched samples) Reagent Kit user guides and as described previously ^7,34^. Briefly, following cell counting (using Trypan blue exclusion), we targeted the capture of 9000 cells and loaded them onto the 10x Genomics Single cell A and B Chips for the total liver homogenate and immune-enriched samples, respectively. cDNA libraries were prepared as per the Single Cell 3′ Reagent Kits v3 user guide. TLH samples were sequenced on a HiSeq 2500 and the immune-enriched samples were sequenced on a NovaSeq 6000. Sequencing QC summaries for each liver profile are found in Table S1.

### Quality control, normalization, and map integration

All the FASTQ files were run on 10 Genomics cell ranger 3.1.0 pipeline with reference genome Rattus_norvegicus.custom_6.0.98. The CellRanger (10X Genomics) analysis pipeline was then used to construct the gene expression matrix from all rat samples. The resulting raw gene expression matrix was filtered based on established quality control criteria (library size, mitochondrial transcript ratio, and the number of expressed genes per cell) using R (version 4.0.3) [https://www.R-project.org/]. Parameters for all quality control criteria were optimized for each sample using a parameter scan and parameter effectiveness was evaluated by manual inspection of the quality of the resulting clustering, visualization, and cell-type annotation, as established ^29^. Parameters were optimized separately for each sample of the total liver homogenate and immune-enriched maps, as each had different quality levels (Figure S1). Various parameters were tested for each sample to maximize low-quality cell (indicated based on library size and mitochondrial gene transcript ratio) removal while minimizing the loss of viable cells. Cell filtering was performed as follows: cells with low (total liver homogenate: [DA-1, LEW-1, LEW-2 <1500; DA-2 <2000], immune-enriched: <1000) library size and high (total liver homogenate: [DA-1>30; DA-2>20; LEW-1>40; LEW-2>40], immune-enriched: >50) mitochondrial gene transcript ratio were removed. The distribution of quality control covariates over the total liver homogenate map indicates that no cluster is highly enriched in these covariates (Figure S2). As expected, hepatocyte clusters have slightly higher mitochondrial gene expression. We also evaluated three different mitochondrial fraction cut-offs to ensure that our map was robust at all mitochondrial cut-offs (Figure S3). Because of additional washing steps and removal of ambient RNA applied to immune-enriched samples, the immune-enriched map had a higher baseline quality, therefore, less stringent QC parameters needed to be applied. The final version of the total liver homogenate and immune-enriched maps includes 23036 (cells per sample: 6623; 7112; 5457; 3844) and 3830 (cells per sample: 1161; 2669) cells respectively. The median expressed genes per cell ranged from 768 to 974 for the total liver homogenate, and the immune-enriched map’s values were 1138 and 1228.

Normalization and clustering of the data were performed using the Seurat (version 4.0.2)^75^ software. Each input sample was normalized using Seurat’s default ‘scTransform’ ^76^ normalization method, which implements a regularized negative binomial regression model for each gene. Samples were then concatenated (merged) to construct the total liver homogenate (n=4 samples) and immune-enriched (n=2) maps. After scaling the merged gene expression matrices, principal component analysis (PCA) ^52^ was used to reduce the number of dimensions representing each cell. A scree plot was used to determine the number of principal components to use for our data set, based on selecting an elbow, as established. 15 principal components were used. Harmony (version 1.0) ^28^ integration was then applied to the principal components of each map to remove the technical batch variations. Non-linear dimension reduction methods, and Uniform Approximation and Projection method (UMAP) ^77^ were applied to Harmony-adjusted top components for visualization.

Doublet detection was performed using the scDblFinder (1.10.0) package ^78^. The “supposed doublets” had a uniform distribution within the maps and were not removed (Figure S6). Plots are generated using the ggplot2 ^79^ package in the R environment.

### Cell clustering, differential expression, cluster annotation

Seurat’s shared nearest neighbor Louvain clustering algorithm was used to cluster the cells, based on the Harmony-corrected principal components. Differentially expressed (DE) genes associated with each cluster were identified using Seurat’s FindMarkers (logfc.threshold = 0, min.pct=0, min.pct=0, min.cells.group = 1) implementation of the non-parametric Wilcoxon rank-sum test. scClustViz ^80^ was incorporated into the clustering pipeline to help find the optimal clustering resolution manually, based on known cell annotations ^29^. Resolution 0.6 was chosen for both the total liver homogenate and immune-enriched maps. The *Ptprc*^+^ clusters of the immune-enriched map were subclustered to examine cell subtypes, and in this case resolution, 1.0 was used. Manual cell annotation involved evaluating the top DE genes based on known markers according to the literature.

### Matrix Factorization using varimax PCA

We used matrix factorization to separate out and study the hidden patterns (factors) within our scRNA-seq data^81^, which may represent factors such as cell type gene expression program or a technical factor. Matrix factorization decomposes the gene expression matrix into the product of two lower-dimension matrices: 1) the loading matrix, which defines the relationship between the genes and the factors and can be used for pathway analysis and gene expression marker discovery; and 2) the score matrix, which represents the relationship between the factors and the cells and can be used for cluster analysis and data set visualization. Here, we used a matrix factorization method called varimax PCA ^50^ to identify the hidden factors within our healthy single-cell RNA-seq rat liver maps. Standard PCA identifies orthogonal dimensions that capture the maximum amounts of variation in the data. Varimax PCA applies an orthogonality-maintaining rotation to the PCA loading matrix with the goal of improving the interpretability of the PCs. This higher interpretability is mathematically achieved by maximizing the variance of the squared loadings in each factor ^50^.

Varimax PCA was applied to the normalized total liver homogenate and immune-enriched gene expression matrices separately. Interpretation of the varimax factors starts with matching factors with cell clusters and known covariates of interest (e.g. strain, sex). Varimax factors were serially plotted against PC1, to create a two-dimensional plot to help visually identify whether a separation on the basis of a specific cluster or strain was evident. For instance, different distributions of DA and LEW-derived cells over the factor of interest indicate that it has captured strain-specific variations (Figure 2C-2F). Other factors visually correlated with known cell types (Figure 2A, Figure S13).

We used correlation analysis and random forest (RF) binary classifiers to automate the factor interpretation process. The correlation between the average gene expression of each cell cluster and the loading scores of each varimax factor was calculated. The top 10 differentially expressed genes of each cell population (cluster) were used to calculate the Pearson correlation scores. The results were plotted as a heat map (Figure 2B), which was used to match each cluster with one or more varimax factors with a high absolute correlation value. The resulting matched factor and cluster pairs were robust to the number of selected top DE genes (10, 20, 30, 50). A Random Forest model was used to identify the varimax factors that capture strain-specific variations. This classifier was trained to predict the strain attributes of each cell by using varimax factors as input features. Evaluating the feature importance of the trained model uncovers the most informative varimax factor to predict the strain of interest. The model was implemented by the randomForest ^82^ (version 4.6.14) package and evaluated using the caTools (version 1.18.2) [https://CRAN.R-project.org/package=caTools] library (Accuracy : 0.9995, Sensitivity : 0.9994, Specificity : 0.9996). The feature matrix (varimax factors) and the corresponding labels were split in a 3:1 ratio using the ‘sample.split’ function of the caTools package into the train and test sets, and the feature importance of the trained models was assessed. The factor with the highest feature importance score (as measured by the mean decrease in Gini score ^83^) was chosen as the best-matched factor for the predicted covariate.

To deconvolve strain-associated biological variations from sample-related confounding factors, at least two samples per strain are required. Consequently, the immune-enriched map’s strain-specific varimax factors were disregarded. Two strain-specific factors were identified from the total liver homogenate map. To assess whether the varimax PCs represent technical or biological signals, the correlation between each factor and three major technical covariates, including library size, number of expressed genes, and percentage of mitochondrial gene expression was calculated. All the strain-specific components indicated a near-zero correlation with these technical covariates (Figure S13C). To further confirm the biological relevance of strain-specific factors found in the total liver homogenate map, we created a gene signature for each strain-related factor (PC5 and PC15) by selecting the top 10 positively and negatively loaded genes for each and used these to score each cell within each strain sample of the immune-enriched map, using the UCell ^84^ package (version 1.0.0) (Figure S29, Figure S30).

### Pathway and gene set enrichment analysis

Gene-set enrichment and pathway analysis methods were used to study the biological signatures represented by each factor. The gene scores corresponding to the factors of interest were selected from the loading matrix to order the list of genes from most to least contribution to the given factor. Pathway enrichment analysis was performed on the ordered list of genes using Gene Set Enrichment Analysis (GSEA))^85^ software from the Broad Institute (software. broadinstitute.org/GSEA) using default parameters (parameters: collapse=false, nperm=1000, scoring_scheme=weighted, plot_top_x=20, rnd_seed=12345, set_max=200, set_min=15) the Gene Ontology Biological Process gene set database (Rat_GOBP_Allpathways_no_GO_iea_May_01_2021_symbol.gmt from http://baderlab.org/GeneSets). To identify activated transcription factors, the gProfiler ^86^ [https://biit.cs.ut.ee/gprofiler/gost] enrichment tool was used with the CHEA-2016 gene set database [https://maayanlab.cloud/Enrichr/#stats]. GSEA results were visualized using the EnrichmentMap ^87,88^ (version 3.3.2) and AutoAnnotate apps ^87,88^ (version 1.3.4) in Cytoscape ^89^ (version 3.8.2).

### Rat/Mouse hepatic zonation correlation analysis

Rat-mouse orthologous genes were identified from the Ensembl database using the biomaRt ^90^ packages (version 2.46.2). Using the significantly (q value < 0.01) differentially expressed genes identified by the Halpern et al. ^30^ study for nine layers of mouse liver cells, we selected 1102 and 1088 genes detected in both rat and mouse in total liver homogenate and immune-enriched maps, respectively. Expression values of each gene among the hepatocytes clusters of rat datasets and nine layers of mouse liver cells were scaled and centered (separately in rat and mouse) by z-scores.

Finally, Pearson correlation was calculated using z-scores across all the selected genes to compare our rat hepatocytes clusters with the nine layers of mouse liver cells in Halpern et al. (Figure S4)

### Rat/Human and Rat/Mouse liver map comparison

Previously published human ^7^ and mouse ^32^ healthy liver maps were downloaded to compare with both total liver homogenate and immune-enriched maps (Figure 5I, Figure S5, Figure S10, Figure S23). The specific pathogen-free (SPF1-3) samples (considered ‘normal’ samples) from the mouse liver data were selected and pre-processed using Seurat’s standard pipeline. Rat and human/mouse orthologs were identified using Ensembl biomaRt as described above. In each pairwise cross-species cell type comparison, the one-to-one orthologs in the top 2000 highly variable genes of the two maps were used for Pearson correlation calculation (final number of one-to-one orthologs genes in each comparison: rat total liver homogenate-human: 517, rat immune-enriched-human: 591, rat total liver homogenate-mouse: 623, rat immune-enriched-mouse: 670). The final heatmap was clustered using Ward’s hierarchical clustering.

### Total liver homogenate map’s mesenchymal population correlation analysis

The mesenchymal cluster of the total liver homogenate map (clusters 7 and 14) were subclustered to perform correlation analysis with mesenchymal subpopulations of Dobie et al., 2019, and Andrews et al., 2022 datasets (Figure S8). The average gene expression of the mesenchymal populations of the three single cell transcriptomics maps were calculated. Pearson correlation was performed based on the one-to-one orthologs in the top 2000 highly variable genes of the maps (final number of one-to-one orthologs genes in each comparison: rat total liver homogenate-Andrews et al.: 1644, rat total liver homogenate map-Dobie et al., 2019: 1161).

### Intracellular Cytokine Stimulation Assay

Rats matching the same strain and weight as the ones used for the scRNA-seq were sacrificed in as above to generate a total liver homogenate. Centrifugation at 50 x *g* for 5 minutes was used to remove hepatocytes and produce the non-parenchymal cell fraction (NPC) from the TLH. To examine the inflammatory potential of myeloid cells in Lewis vs. Dark Agouti rats, NPC fractions were cultured for 4 hours for adherence and subsequently stimulated for 6 hours in 12-well tissue culture plates with 1ng/mL of LPS in the presence of BFA/monensin. The cells were harvested and intracellular secretion of TNFα was examined using flow cytometry.

### Flow Cytometry

The collected cells were first stained with live/dead aqua to exclude non-viable cells from the analysis and stained with fluorophore-conjugated antibodies against surface markers anti-CD45-BV786 (BD Bioscience Clone: OX-1), anti-CD11b-V450(BD Bioscience Clone: WT.5), anti-CD11a-PE (BD Bioscience Clone: WT.1), anti-CD3-BV510(BD Bioscience Clone: 1F4). The cells were then fixed with 1X fixation buffer (eBioscience) and permeabilization buffer 1X (eBioscience), and then stained with the following intracellular antibodies; monoclonal CD68-AF700 (Novus Biologicals Clone: ED1), TNFα (AbCam Clone: EPR21753-109). Each surface staining and intracellular staining step was accompanied by a rat FcBlocking step via an anti-CD32(BD Bioscience Clone: D34-485). Events were acquired on a 5-laser custom BD Fortessa X20 analyzer. The gating strategy for both cell surface markers and intracellular markers was based on Fluorescence Minus One (FMO) controls for each marker. Intracellular cytokine TNFα gating strategy was based on the fluorescence seen in both FMO and the unstimulated control (Figure S36). Event analysis was performed using FlowJo.

### Immunohistochemical Staining

Paraffin-embedded sections from rat liver were stained by the Pathology Research Program (PRP) at the Toronto General Hospital according to standard histological procedures. Paraffin-embedded rat tissues were stained with antibodies for CD3 (Roche, 2GV6), CD8 (Bio-Rad, OX-8), and CD68 (Abcam, ab125212). The stained slides were scanned by the University Health Network Advanced Optical Microscopy Facility using a Leica Aperio AT2 whole slide scanner (Leica Microsystems, Carlsbad CA), and converted into digital images. The QuPath software version 0.2.3 software was used to perform image analysis and quantification according to the protocol available at dx.doi.org/10.17504/protocols.io.bs6gnhbw.

## Supporting information

Extended Results

## Acknowledgments

This project has been made possible in part by grant number CZF2019-002429 from the Chan Zuckerberg Initiative DAF, an advised fund of Silicon Valley Community Foundation. This research was supported in part by the University of Toronto’s Medicine by Design initiative, which receives funding from the Canada First Research Excellence Fund (CFREF) to SAM, GDB, and IDM; by the NRNB (U.S. National Institutes of Health, grant P41 GM103504) to GDB and by the Toronto General and Western Hospital Foundation. This study was supported by the Canadian Institutes for Health Research, (grant no: PJT 162098 to SAM and PJT 162298 to IDM, and PJT 469829 to GDB). We would like to acknowledge Jawairia Atif and Tallulah S. Andrews for their helpful discussions.

## Author contributions

Conceptualization, D.P., S.W.C., I.D.M, S.A.M., and G.D.B.; Methodology and software, D.P; Investigation, D.P., S.W.C.; Validation, S.W.C., D.C., X.W.; Resources, C.T.P, X.Z.M., C.J., M.S., J.M. and X.C.C., D.C; Formal Analysis, D.P.; Data Curation, O.I.P., S.W.C., T.C.; Writing – Original Draft, D.P., S.W.C., S.A.M., G.D.B; Writing – Review & Editing, S.A.M., G.D.B, I.D.M; Supervision, G.D.B., S.A.M., I.D.M; Funding Acquisition, G.D.B., S.A.M., I.D.M.

## Ethics

All experimental procedures followed principles and guidelines for the care and use of animals established by the Animal Resources Centre (ARC) at the University Health Network and are in accordance with the guidelines of the Canadian Council of Animal Care. Rat experiments were performed at the Toronto General Research Institute, Toronto, ON, Canada under the approval of the Institutional Committee on Animal Bioethics and Care (AUP 5840). All surgery was performed under isofluorane anesthesia, and all efforts were made to minimize suffering.

## Code availability

R Scripts for the data process are available through https://github.com/BaderLab/HealthyRatLiverMap

## Data availability

Interactive atlases of the total liver homogenate, immune-enriched, and the immune-subcluster maps are available through the Cell Browser Interface: https://rat-liver-atlas.cells.ucsc.edu

