## Extended Results for "Single cell profiling reveals strain-specific differences in myeloid inflammatory potential in the rat liver"

### Total liver homogenate map lymphocyte populations

*Ptprc*+ cells within the first four samples formed four major clusters, including two *Marco*$^{+}$*Cd5l*$^{+}$*Cd68*$^{+}$ myeloid clusters (Clusters 5 and 10), one cluster with an enriched expression of *Cd68* and *Cd3e* which appeared transcriptionally to represent a mixture of recently recruited monocyte/macrophage populations and *Cd3*+ T cells (Cluster 9), and a *Ptprc+Cd68-Cd3e-Nkg7+* lymphocyte cluster (Cluster 13) (Figure 4A) which showed enriched expression of NK cell genes *Klrd1*, *Gzmk*, and *Cd7* (Cluster 13 top DE: *Ccl5, Gzma, Nkg7, Gzmm, Tyrobp, Xcl1, Prf1, Tmsb10, Crip1, Cd7*), suggesting that cluster 13 was an NK-like cluster.

### Immune-enriched map myeloid and *Ptprc*+ populations subclustering

To examine immune populations with higher resolution, after enriching immune cells with RBC lysis and additional washing steps, we generated a map that enabled better capture of myeloid and lymphocyte fractions. The immune-enriched map included seven *Ptprc*+ clusters, including two *Marco*+ myeloid clusters (Clusters 8,13), *Cd3*+ T cells (Cluster 10), NK-like cells (Cluster 7), B cells (Cluster 12), plasmacytoid dendritic cells (Cluster 17), and a mixed myeloid and cross-presenting dendritic cell population (Cluster 11), as described further below (Figure 4G).

Our analysis revealed three myeloid clusters identified by *Cd68* expression. *Cd68*+ cells in clusters 8 and 13 were characterized by enriched expression of *Marco, Vsig, Cd5l, Cd163,* and *Hmox1,* which are typical tissue-resident Kupffer cell-like markers[1]. Cluster 8 top DE genes are *Marco*, *C1qa*, *Ccl6*, *C1qb*, *C1qc*, *Clec4f*, *Cd5l*, *Vsig4*, *Aif1*, *Npc2*. Cluster 13 top DE genes are *RT1-Db1*, *C1qa*, *C1qb*, *Cd74*, *RT1-Da*, *Clec4f*, *Aif1*, *C1qc*, *Npc2*, *Ccl6*). These DE genes highlight tolerogenic (*Vsig4*) and resting Kupffer cell (*Clec4f*) [2,3] functions. Co-inhibitory ligand Vsig4 maintains the intrahepatic tolerance required to mediate the inflammatory cascade [4]. The enriched genes and pathways in this cluster suggest the presence of tissue-resident KC-like macrophages. Cluster 11 is characterized by enriched expression of recently recruited monocyte/macrophage markers *S100a8*, *S100a9*, *Cd74*, *Cts3*, and *Lyz2* (the mouse marker for inflammatory macrophage gene *LYZ* in the human liver). Cluster 11 top DE genes are *Lyz2*, *S100a9*, *Slpi*, *S100a8*, *S100a4*, *RTI-Db1*, *S100a6*, *RTI-Da*, *Cst3*, *RT1-Bb*. Furthermore, Cluster 11 includes populations of *Xcr1+Clec9a+Batf3*+ and *Clec10a*+ cells annotated as cDC1s and cDC2s respectively. Based on the expression of these key genes and following a rat-human liver map correlation analysis (Figure 4I), we propose that *Cd68*+ clusters 8 and 13 were more tissue-resident KC-like in nature, and *Cd68*+ cluster 11 is more of a recently blood-recruited macrophage population and contains populations of cDC1 and cDC2 cells.

### Immune-enriched map *Ptprc*+ subclustering

The *Ptprc*+ populations from the immune-enriched map were subclustered (see Methods) to examine the hepatic immune population in more detail (Figure S28). This subclustering revealed 14 immune subpopulations (Figure S28A, Table S8). Amongst these subclusters, we identified seven subclusters containing Cd68+ myeloid cells (2, 4, 6, 9, 10, 11, 13), two T cell subclusters (0 and 12), two NK-like cell subclusters (1 and 8), one B cell subcluster (3), one pDC subcluster (7) and finally, a cDC subpopulation (5). We identified some of these subclusters as heterogeneous cellular populations containing transcripts associated with LSECs, and hepatocytes, which may indicate that some non-immune cells could have been grouped with the immune cells through the initial clustering due to less-well-defined boundaries for some clusters (e.g. myeloid cluster 13, Figure 4A).

**T Cells.** In this dataset, we identified two *Cd3*+ T cell populations (Figure S29). The most abundant T cell subcluster (Subcluster 0) was identified based on the enriched expression of *Cd3* genes (*Cd3g, Cd3e, Cd3d, Cd2*) suggesting that this subcluster is a *Cd3*+ T cell population. Additionally, this T cell subcluster included ribosomal protein genes such as *Rpl12, Lef1*, and *Rps16* (Top DE Genes: *Cd3g, Cd3e, Cd3d, Lef1, Fam189b, Rps18, Rpl12, Lgals1, Rps16, Rps15a, Cd2, Cd8a, Cd247*). Subcluster 12 is the second population characterized as *Cd3*+ T cell. This subcluster showed enriched expression of γδ T cell gene *Tbx21* (aka T-bet), and phosphoantigen reactive γδ T cell genes *Top2a, Nusap1,* and *Cdca8* [5]. The T cells in this subcluster appeared to be highly proliferative as the most highly expressed DE genes *Stmn1* and *MKi67* which are expressed in proliferating and dividing T cells [6]. (Top DE Genes: *Stmn1, Hmgb2, Top2a, Mki67, Tuba1b, Ube2c, Cenpf, H2afx, Nusap1, Gzma*).

**NK-Like Cells.** Two NK-like cell subclusters (Subcluster 1 and Subcluster 8) were found in the *Ptprc*+ sub clustering map (Figure S30), characterized by expression of *Klrd1, Ncr1*, and *Gzmk* without upregulation of *Fcgr3a* or *Itga1*. Expression of *Gzmk* and *Prf1* genes is essential to NK cell identification, as literature has indicated that hepatic NK cells respond to antigens by releasing lytic granules expressing high levels of granzyme and perforin genes [7]. (Subcluster 1 Top DE Genes: *Ccl5, Gzma, Xcl1, Prf1, Gzmm, Nkg7, Ccl4, Klrb1a, Klrd1, Il2rb*) (Subcluster 8 Top DE Genes: *Gzmk, Gzmm, Nkg7, Ccl5, Klrb1a, Prf1, Il2rb, Klrd1, Gzma, Ncr1*).

**B Cells.** Amongst the immune subcluster map, we identified a *Cd79b+Ighm*+ B cell-like subcluster (Subcluster 3), characterized by enriched expression of *Ighm*, *Cd74, Fcmr, Cd19, Ms4a1(Cd20)*, and *Cd79b*, with no expression of *Ighd* or *Ighg,* suggesting that this subcluster might be Cd19+Cd20+*IgM+IgD-* immature B cells [8] (Figure S31).

**Myeloid cells**. The immune-enriched subclustering map revealed seven subclusters (2, 6, 9, 11, and 13) containing myeloid cells (Figure S32). *Clec4f* is a known KC C-type lectin receptor that recognizes desialylated glycans and is involved in platelet destruction within the liver [9]. *Clec4f* was a highly differentially expressed gene in subclusters 2, 6, 9, and 13. The key differentially expressed genes in these clusters are described below.

**Subclusters 2** and **13** are identified as *Clec4f*/*Vsig4*+ myeloid clusters based on their top DE genes (subcluster 2 top DE Genes: *C1qa, Cd5l, Clec4f, Vsig4, C1qb, C1qc, Marco, Hmox1, Cd163, Cd68*; subcluster 13 top DE Genes: *Cd5l, Clec4f, Cd163, Ccl6, C1qc, Mrc1, C1qb, Slc40a1.1, Vsig4, Hmox1, Ax*l). Subcluster 6 is a predominantly *Clec4f*+ population that appears to be contaminated with hepatocytes, as the top expressed genes include both KC-like myeloid genes *C1qa, Aif1*, and *Clec4f*, and hepatocyte genes *Alb, Serpina1, Hp,* and *Apoa1* (Top DE Genes: *C1qa, C1qb, Aif1, Clec4f, Gc, Serpina1, Ttr, Apoc3, Ifitm3, Apoa1, Fabp1*). Subcluster 9 is a predominantly *Clec4f*/*Vsig4*+ population that appears to be contaminated with LSECs, showing enriched expression of *Sparc* and *Calcrl,* KC-like genes *C1qb, Mrc1, Vsig4,* and *Aif1* (Top DE Genes: *Kdr, Sparc, Fcgr2b, Clec4f, Mrc1, C1qc, C1qb, Lyve1, Fam167b, Tspan7).* Subcluster 11 appears to be an intermediate population with an expression of *Fcgr3a* (CD16) and no expression of *Vsig4* (Top DE: *RT1-Db1, RT1-Bb, RT1-Da, C1qb, C1qc, Cd74, Fcgr3a, Aif1, Axl, Cd68, Fcer1g*). Subclusters 10 were characterized as *Lyz2+S100a8*/*9*+ recently recruited macrophages based on the enriched expression of *Lyz2, S100a8*, and *Lsp1* (subcluster 10 Top DE Genes: *S100a9, S100a8, Lyz2, Tspo, Dusp1, Il1b, Anxa1, Fos, S100a11, Clec7a, Ctsa, Ev12a, Tyrobp*). Subcluster 4 also showed enriched expression of recently recruited monocyte/macrophage markers *Cd74* and *Tyrobp* (Top DE Genes: *Lyz2, S100a6, Cebpb, Klf4, S100a11, Lsp1, Fcer1g*).

**pDC-enriched:** As in the immune-enriched map, we again found a cluster with an enriched expression of DC genes *Siglech, Ptcra, Ifi30, Tcf4, Runx2,* and *Tlr7* in subcluster 7, suggesting that this subcluster was composed of a DC population (Figure S33). Specifically, enriched expression of differentially expressed *Ptcra* suggests that these dendritic cells may be plasmacytoid DCs (pDCs) because *Ptcra* is a key binding target for pDC master regulatory transcription factor TCF4 [10] (Subcluster 7 Top DE Genes: *Irf8, Gapt, Siglech, Fcrla, RT1-Da, Cd74, Tcf4, Ptcra, Ifi30, Ighm, Runx2, Tlr7*). This cluster contained some contaminating B cell transcripts, including *Ighm*, which suggest that this is a heterogeneous population.

**cDC-enriched:** Importantly, our analysis revealed subcluster 5 which contained a mixture of cross-presenting DCs and myeloid cells (Figure S33). Subcluster 5 revealed enriched expression of recently recruited monocyte/macrophage markers *Cst3* and *Cd74*, as well as cross-presenting DC markers *Xcr1*, *Clec9a*, and *Tlr3* [11]. When looking at the physical distribution of these DC markers in our UMAPs, we see that a subpopulation on the right side of this subcluster had enriched expression of cDC1 genes (*Xcr1*, *Clec9a* [12]) and the subpopulation on the left is enriched in cDC2 markers (*Clec10a* [12], *Tmem176b* [13,14], suggesting that this subcluster contained a mixture of cDC1 and cDC2 cells (Top DE: *RT1-Db1, Cst3, RT1-Ba, RT1-Bb, RT1-Da, Cd74, Lyz2, Lsp1, Clec9a, Xcr1, Tlr3, Glipr1, Fgr*).

Lab. Invest. *72*, 100–113.

7. Vermijlen, D., Luo, D., Froelich, C.J., Medema, J.P., Kummer, J.A., Willems, E., Braet, F., and Wisse, E. (2002). Hepatic natural killer cells exclusively kill splenic/blood natural killer-resistant tumor cells by the perforin/granzyme pathway. J. Leukoc. Biol. *72*, 668–676.
8. Forsthuber, T.G., Cimbora, D.M., Ratchford, J.N., Katz, E., and Stüve, O. (2018). B cell-based therapies in CNS autoimmunity: differentiating CD19 and CD20 as therapeutic targets. Ther. Adv. Neurol. Disord. *11*, 1756286418761697. 10.1177/1756286418761697.
9. Hoover, C., Kondo, Y., Shao, B., McDaniel, M.J., Lee, R., McGee, S., Whiteheart, S., Bergmeier, W., McEver, R.P., and Xia, L. (2021). Heightened activation of embryonic megakaryocytes causes aneurysms in the developing brain of mice lacking podoplanin. Blood *137*, 2756–2769. 10.1182/blood.2020010310.
10. Villani, A.-C., Satija, R., Reynolds, G., Sarkizova, S., Shekhar, K., Fletcher, J., Griesbeck, M., Butler, A., Zheng, S., Lazo, S., et al. (2017). Single-cell RNA-seq reveals new types of human blood dendritic cells, monocytes, and progenitors. Science *356*. 10.1126/science.aah4573.
11. Collin, M., McGovern, N., and Haniffa, M. (2013). Human dendritic cell subsets. Immunology *140*, 22–30. 10.1111/imm.12117.
12. Guilliams, M., Bonnardel, J., Haest, B., Vanderborght, B., Wagner, C., Remmerie, A., Bujko, A., Martens, L., Thoné, T., Browaeys, R., et al. (2022). Spatial proteogenomics reveals distinct and evolutionarily conserved hepatic macrophage niches. Cell *185*, 379-396.e38. 10.1016/j.cell.2021.12.018.
13. Visan, I. (2019). cDC2 subsets. Nat. Immunol. *20*, 1558. 10.1038/s41590-019-0552-5.
14. Brown, C.C., Gudjonson, H., Pritykin, Y., Deep, D., Lavallée, V.-P., Mendoza, A., Fromme, R., Mazutis, L., Ariyan, C., Leslie, C., et al. (2019). Transcriptional basis of mouse and human dendritic cell heterogeneity. Cell *179*, 846-863.e24. 10.1016/j.cell.2019.09.035.
