## Supplemental Figures for "Single cell profiling reveals strain-specific differences in myeloid inflammatory potential in the rat liver"

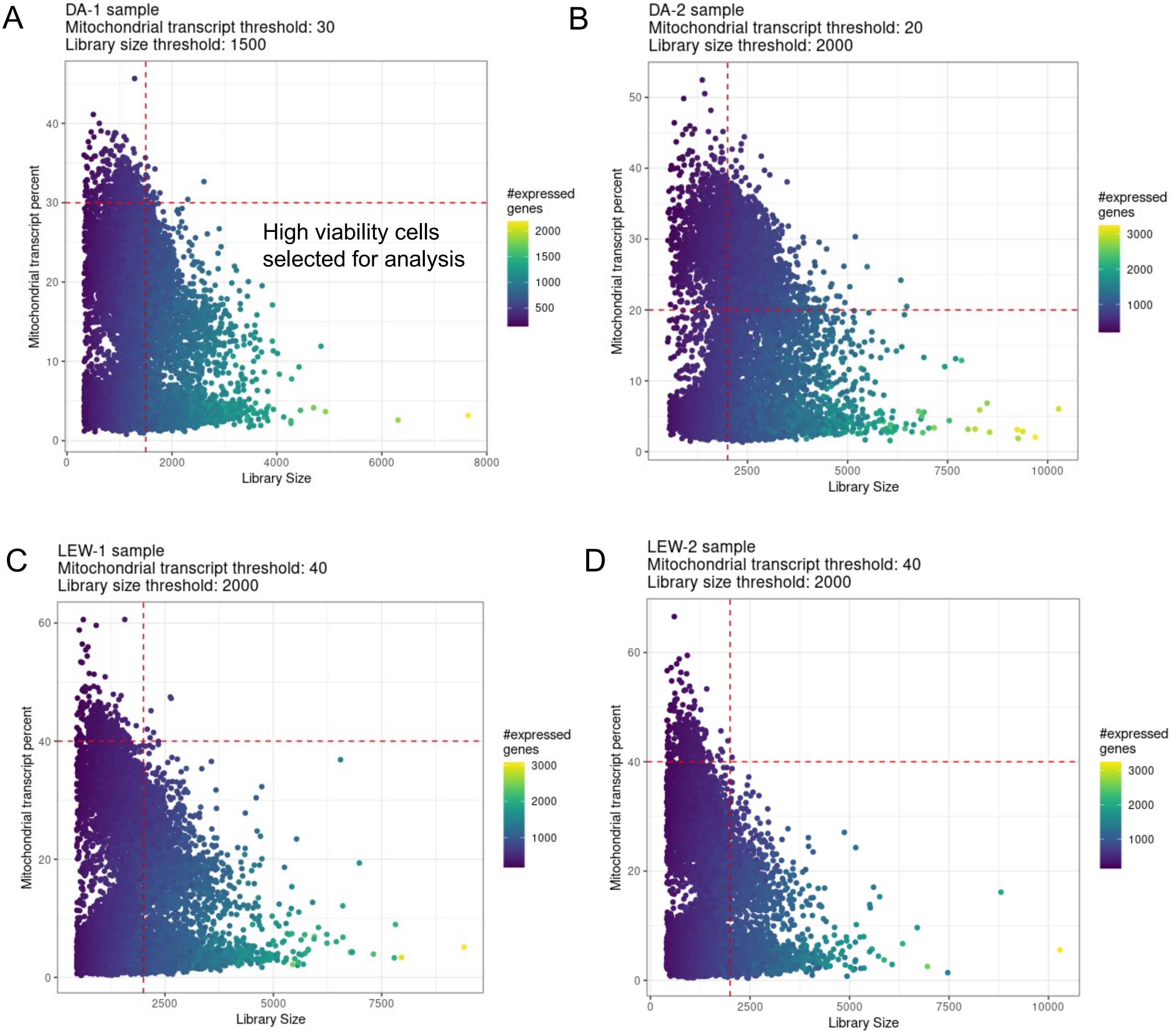


**Supplementary Figure 1: Four healthy rat total liver homogenate samples quality control and selection of high viability cells.** Viable cells were identified from the single-cell gene-expression data based on having a minimum library size of [DA-1, LEW-1, LEW-2] 1500 and [DA-2] 2000 transcripts and a maximum of [DA-1: 30; DA-2: 20; LEW-1: 40; LEW-2: 40] percent mitochondrial transcript proportion. A) DA-1 B) DA-2 C) LEW-1 D) LEW-2 rat healthy liver sample.

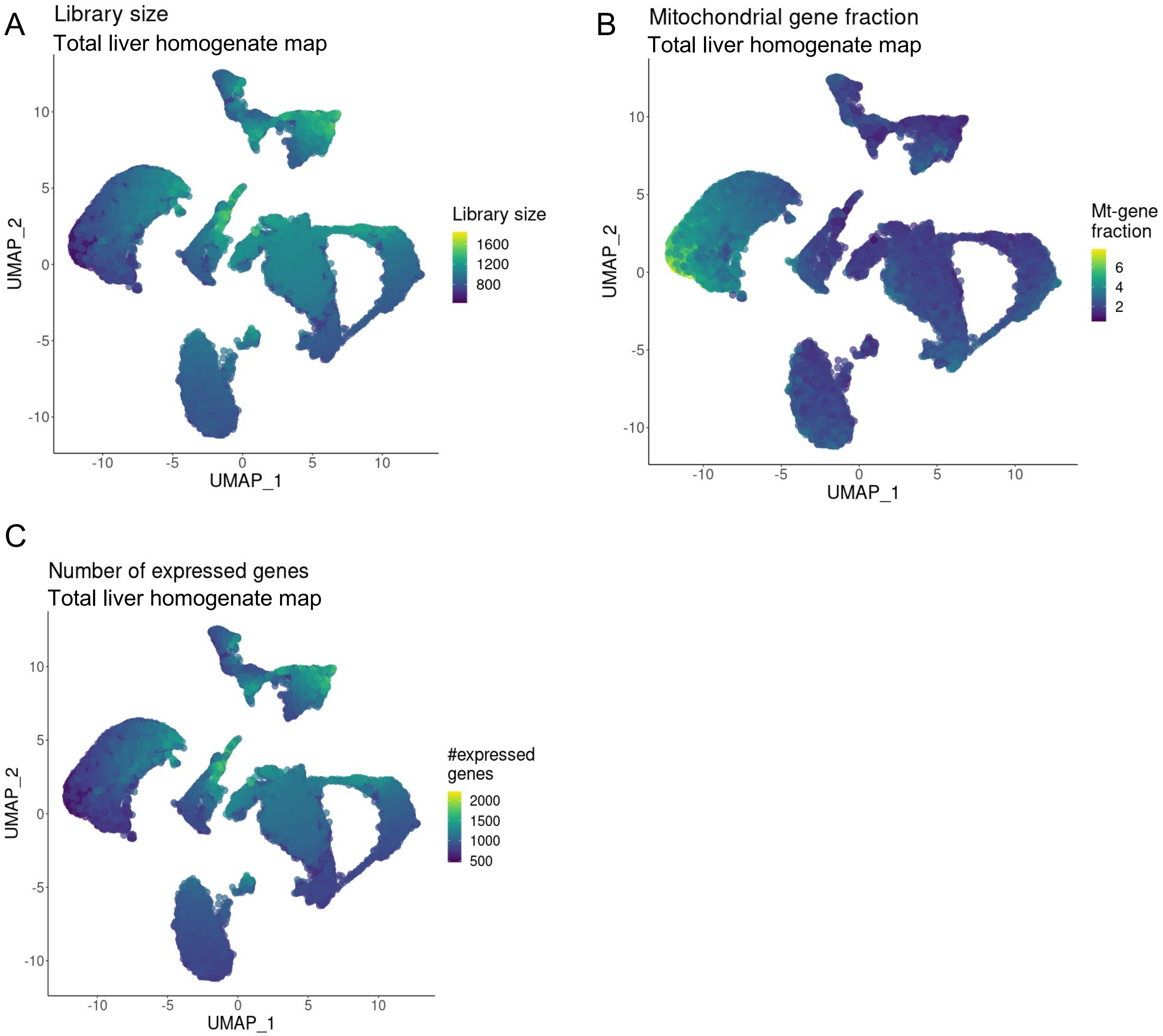


**Supplementary Figure 2: Distribution of quality control covariates over the total liver homogenate map.** UMAP projection of total liver homogenate map where cells are colored based on A) library size, B) mitochondrial transcript proportion and C) the total number of expressed genes in each cell. Yellow indicate higher values and dark blue indicates a lower value of the QC-covariates.

| | A) Mt-gene fraction threshold : 20% Total # cells: 21262 | B) Mt-gene fraction threshold : 30% Total # cells: 22897 | C) Mt-gene fraction threshold : 40% Total # cells: 23036 |
|---|---|---|---|
| **cluster** | **num.cell.mt.20** | **num.cell.mt.30** | **num.cell.mt.40** |
| Hep (0) | 4989 | 4989 | 4989 |
| Hep (1) | 4212 | 4212 | 4212 |
| Marco/Cd5l Mac (10) | 518 | 518 | 518 |
| Endothelial (11) | 517 | 517 | 517 |
| Hep (12) | 446 | 446 | 446 |
| Lymphocyte (13) | 409 | 410 | 410 |
| Mesenchymal (14) | 265 | 265 | 265 |
| Hep (15) | 167 | 167 | 167 |
| Hep (16) | 76 | 126 | 127 |
| Hep (2) | 1182 | 2497 | 2606 |
| Endothelial (3) | 2090 | 2091 | 2091 |
| Hep (4) | 1670 | 1739 | 1740 |
| Marco/Cd5l Mac (5) | 1149 | 1150 | 1150 |
| Hep (6) | 1123 | 1124 | 1124 |
| Mesenchymal (7) | 983 | 984 | 984 |
| Hep (8) | 753 | 949 | 977 |
| Lyz2/Cd74 Mo/Mac (9) | 713 | 713 | 713 |

**Supplementary Figure 3: Cell count of final clusters at various mitochondrial cut-offs.** The number of cells in each cluster (resolution: 0.6) was evaluated for three different mitochondrial fraction cut-offs to ensure that our map was robust at all mitochondrial cut-offs. For this analysis, all samples were filtered using the noted harmonized threshold. In all mitochondrial cut-offs (40%, 30%, 20%), we have cells from all 17 clusters identified.

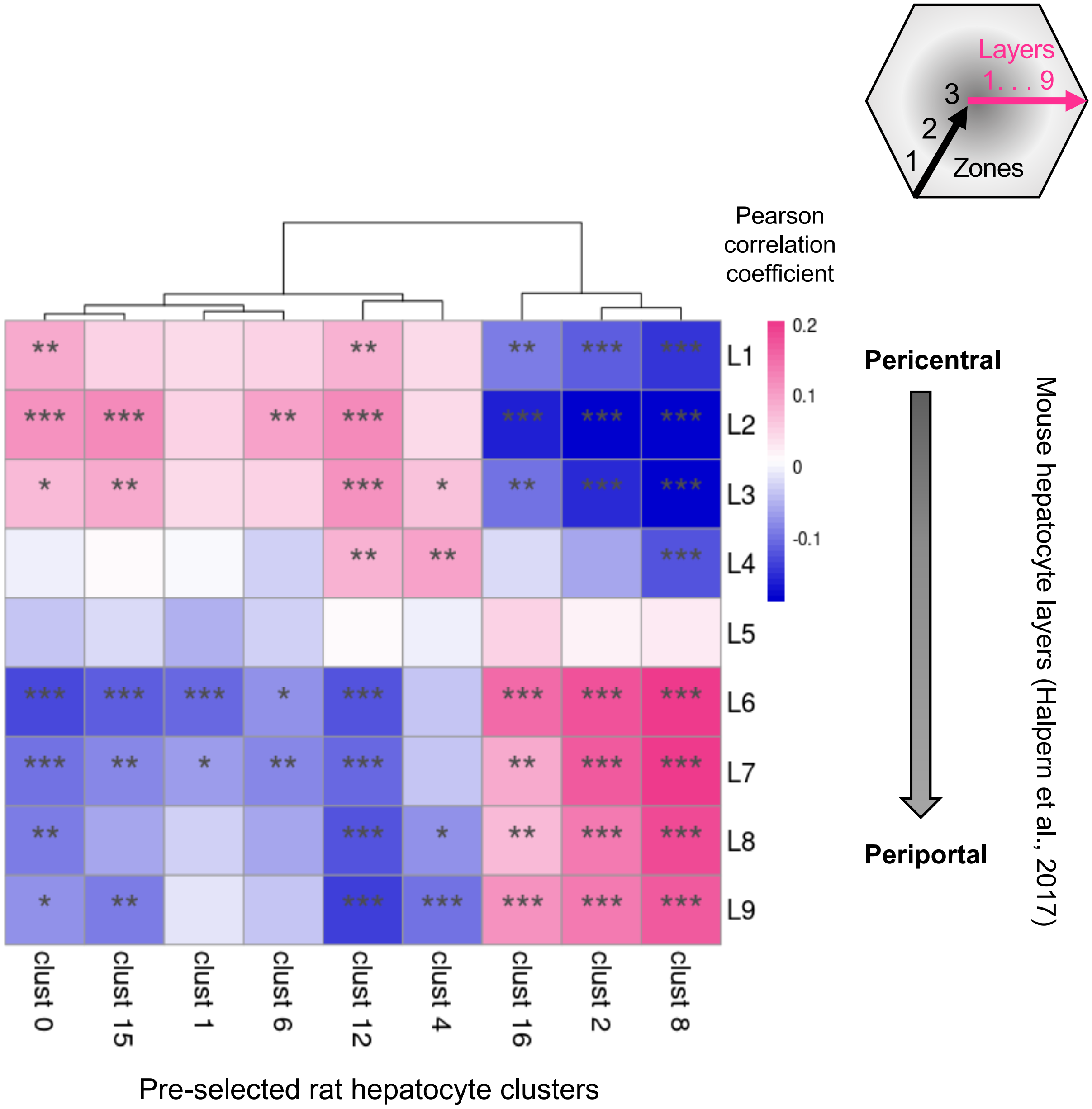


**Supplementary Figure 4: Rat liver map hepatocyte correlation with mouse zonation layers.** Mouse liver layer-9 is more periportal and layer-1 is pericentral. Pearson correlation between the average gene expression of the genes across hepatocyte clusters and the nine layers of mouse liver cells was calculated (see: methods). Red represents positive correlation and blue represents negative correlation. (* : p-value < 0.05, ** : p-value < 0.01, *** : p-value < 0.001)

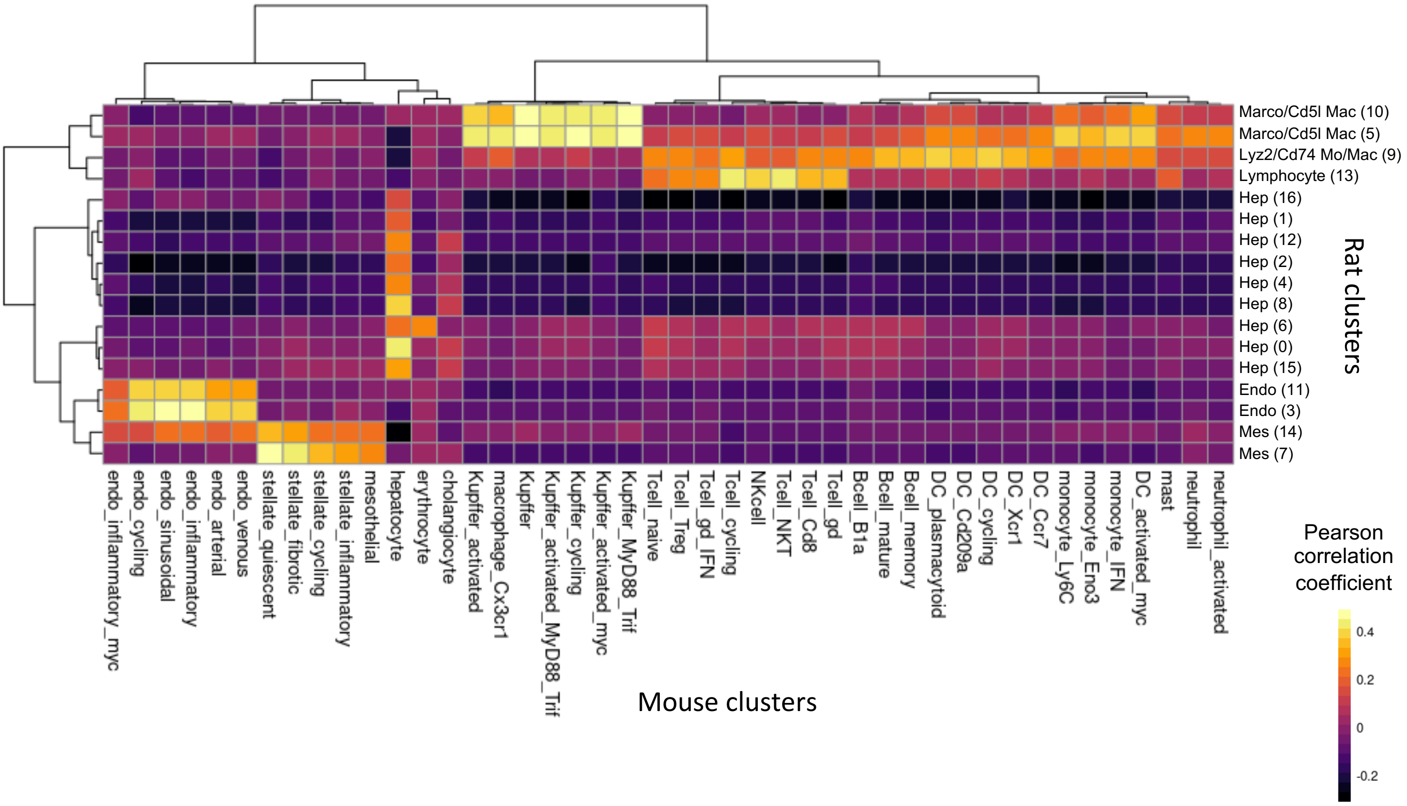


**Supplementary Figure 5: Comparison of rat and mouse liver atlases.** Comparison of the total liver homogenate map and mouse healthy liver map [Kolodziejczyk et al. 2020]. Rows and columns of the correlation heatmap represent the rat and mouse clusters, respectively. The color of the heatmap cells indicates Pearson correlation values between the cluster average expressions. The one-to-one orthologs in the top 2000 highly variable genes of the two maps were used for correlation calculation (see: methods). The comparison indicates a high consistency between the gene expression pattern of hepatic cell types between rats and mice.

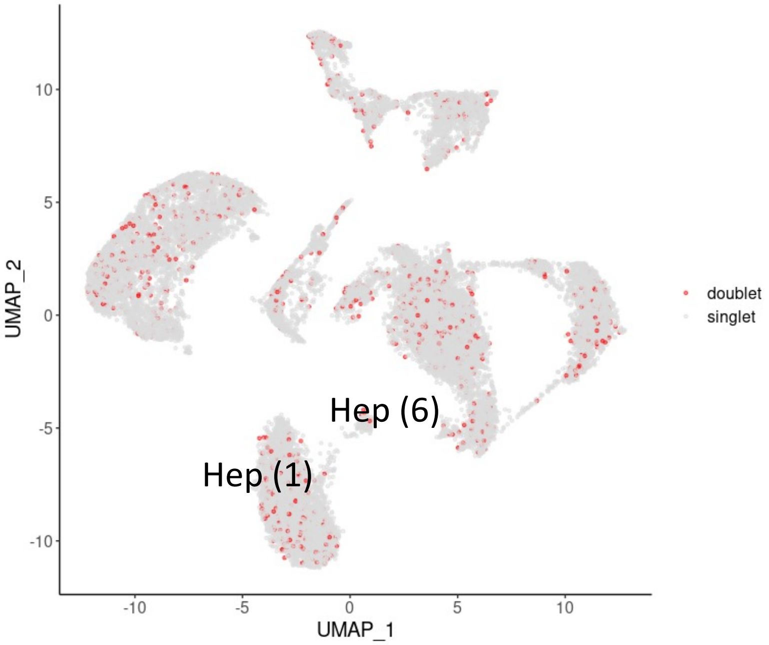


**Supplementary Figure 6: Doublet detection.** A doublet detection algorithm was applied to the rat liver map to identify potential doublets. We decided not to remove these “supposed doublets” since the detected doublet cells had a uniform distribution within the map. Moreover, there are naturally occurring binucleotide hepatocytes in the liver and it’s very challenging to distinguish true binucleotide cells from doublets. The distribution of doublets was not denser in the hepatocyte clusters 1 and 6 compared to the other cell populations, rejecting the hypothesis that these clusters have a high density of erythrocyte and hepatocyte doublets.

Mesenchymal markers (Clusters 7, 14) - TLH map

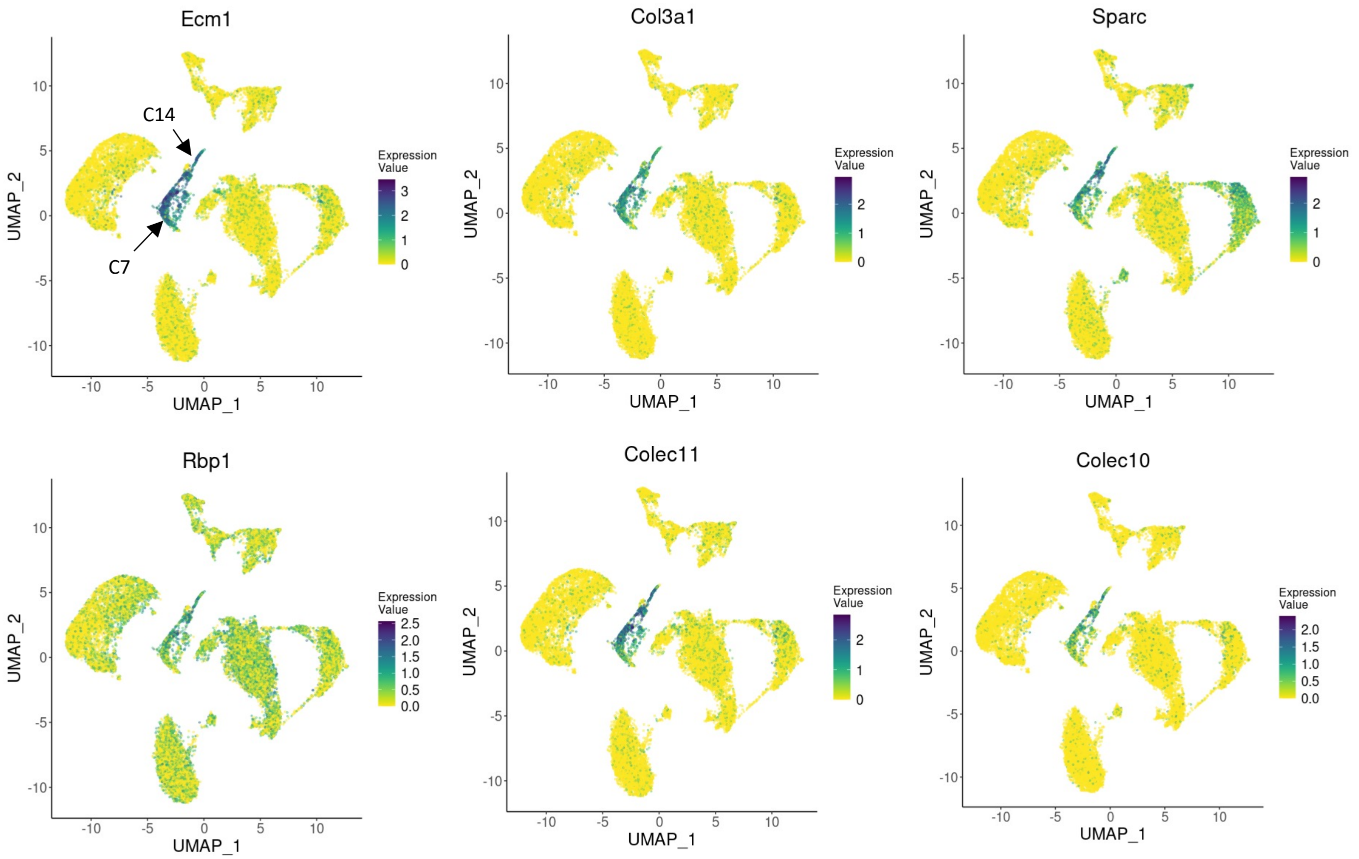


**Supplementary Figure 7: UMAP plots showing the relative distribution of commonly expressed mesenchymal genes (clusters 7, 14) in the healthy rat total liver homogenate map.** Legend for the relative expression of each marker from lowest expression (yellow dots) to highest expression (dark blue dots) is placed on the right. TLH: total liver homogenate, C: cluster.

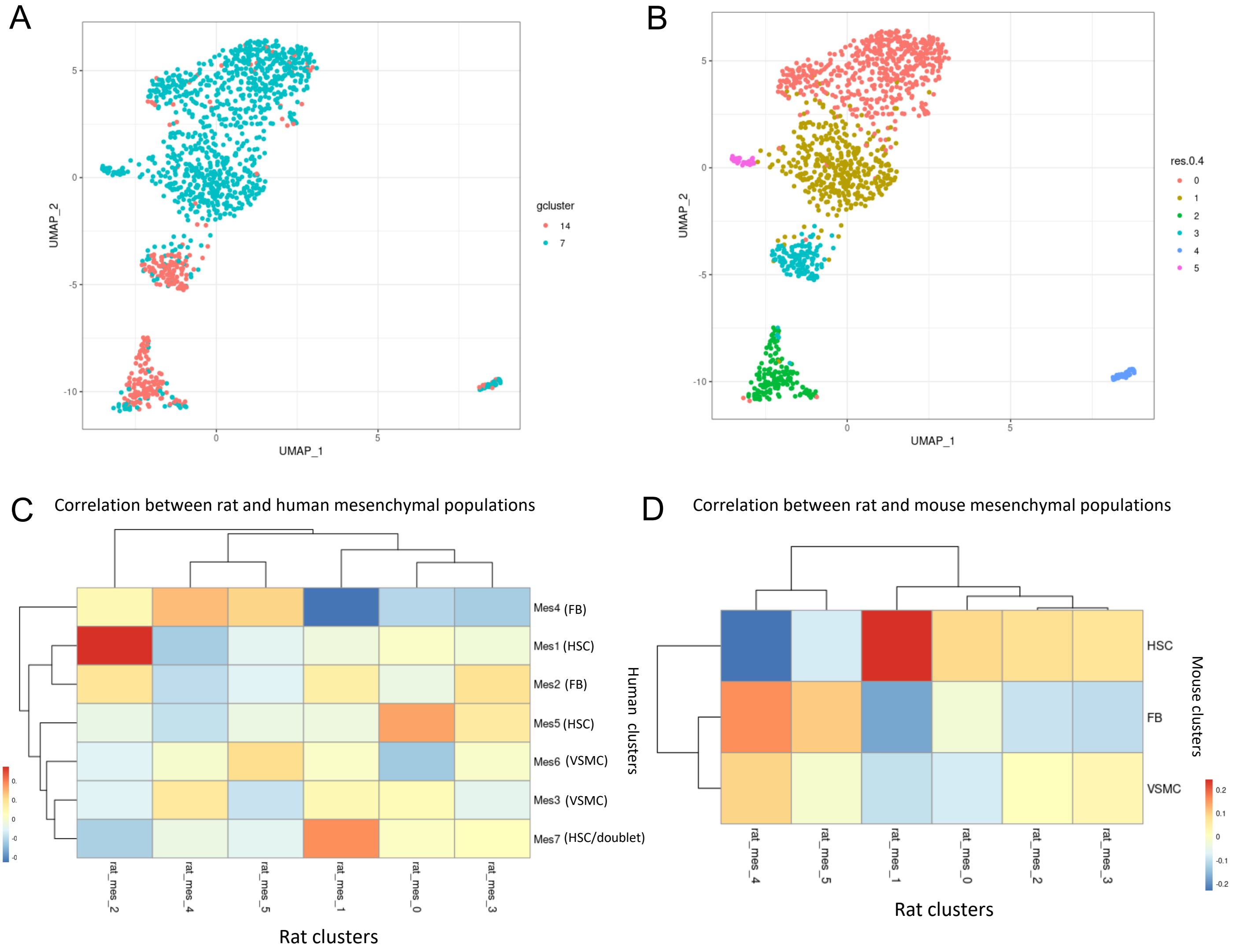


**Supplementary Figure 8: Comparison of rat total liver homogenate map mesenchymal cells with mesenchymal subpopulations of the mouse (Dobie et al., 2019), and human (Andrews et al., 2022) liver transcriptomic maps.** The mesenchymal (mes) population (clusters 7 and 14) of the total liver homogenate map was subclustered (clustering resolution = 0.4) to increase the resolution. A) UMAP plot of mesenchymal subclustering colored by the cluster of origin. B) UMAP plot of mesenchymal subclustering colored by the subcluster number. C) Comparison of rat and human (Andrews et al., 2022) mesenchymal subpopulation. Rows and columns of the correlation heatmap represent the human and rat subclusters, respectively. The color of the heatmap cells indicates Pearson correlation values between the cluster average gene expressions. Rat mesenchymal subcluster 0 (mes-0) is correlated with Mes5 in humans which have been annotated as activated HSCs (hepatic stellate cells) due to expression of HSC-associated retinol storage genes and enrichment of lipid metabolism and fibrin clot pathways and inflammation-associated genes. Rat mes-1 is highly correlated with the mouse and human HSC population. Rat mes-2 is highly correlated with a human quiescent HSC population (Mes1). Rat mes-4 indicates a correlation with a mouse fibroblast (FB) cluster. Annotation of rat mesenchymal clusters 3 and 5 could not be resolved using correlation analysis. VSMC: vascular smooth muscle cell

Endothelial markers (Clusters 3, 11) - TLH map

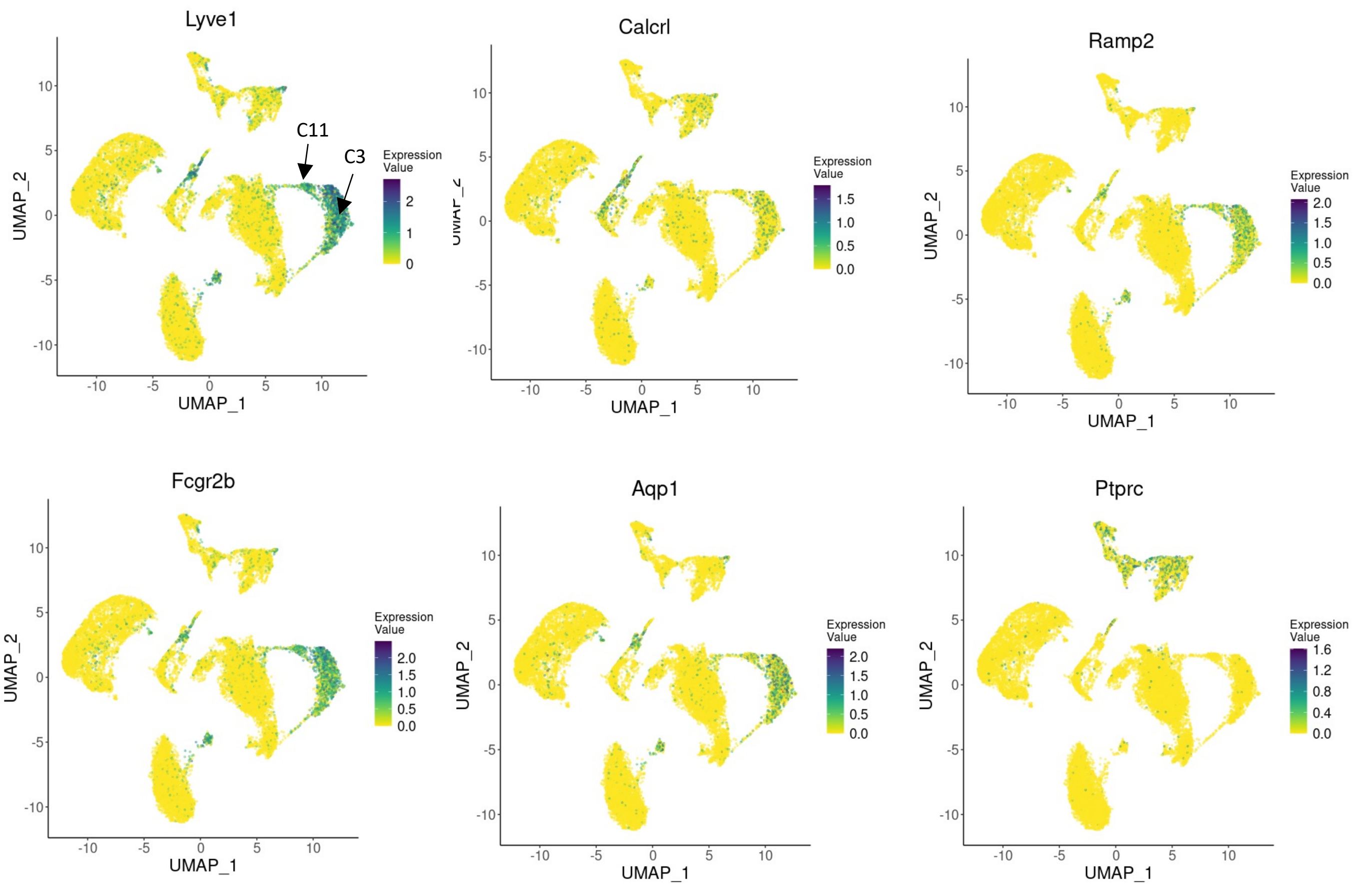


**Supplementary Figure 9: UMAP plots showing the relative distribution of commonly expressed endothelial genes (clusters 3, 11) in the healthy rat total liver homogenate map.** Immune marker *Ptprc* is included for comparison. Legend for relative expression of each marker from lowest expression (yellow dots) to highest expression (Purple dots) is placed on the right. TLH: total liver homogenate, C: cluster

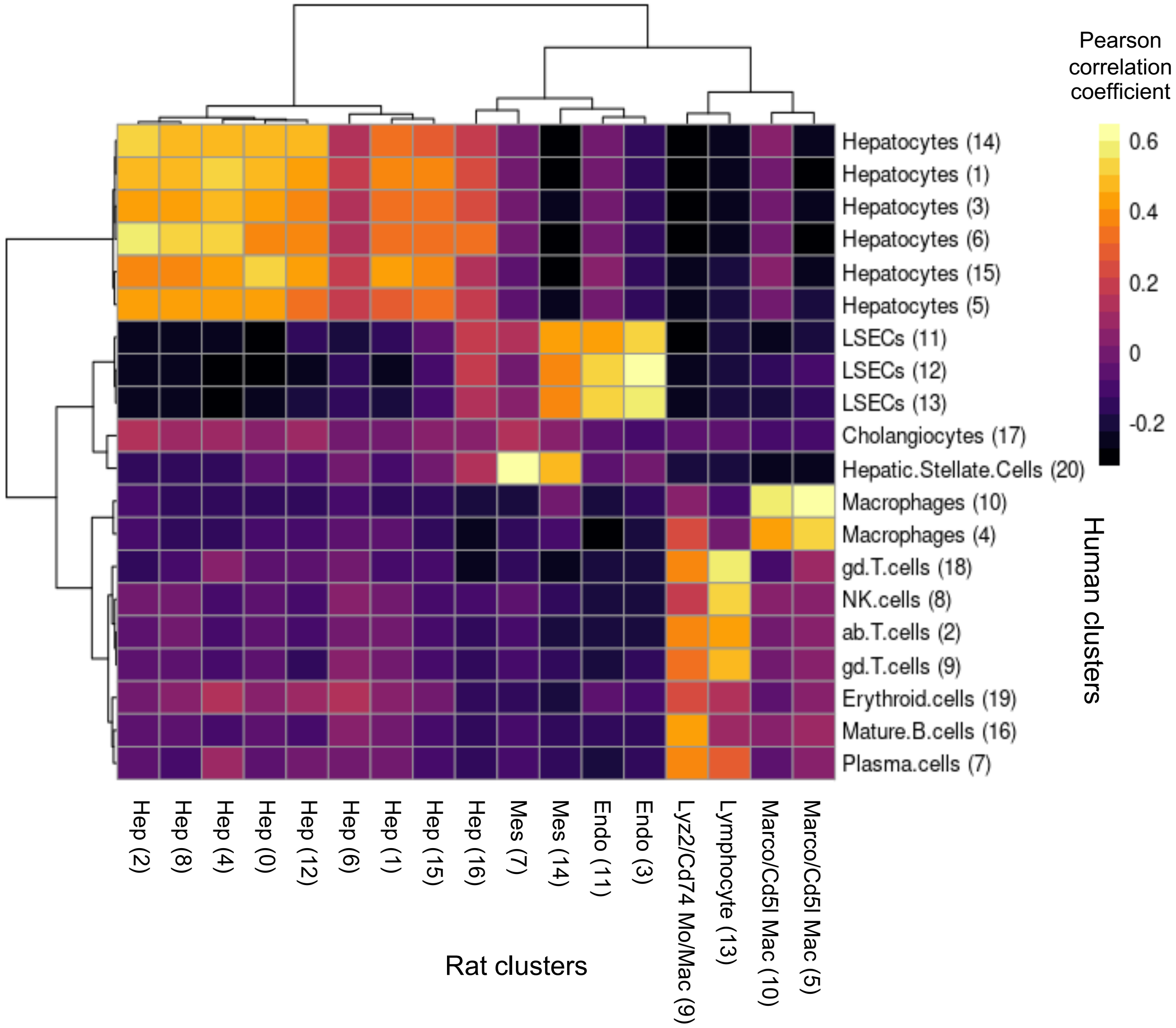


**Supplementary Figure 10: Comparison of rat and human liver atlases.** Comparison of total liver homogenate map and human liver map [Sonya A. MacParland et al.]. Rows and columns of the correlation heatmap represent the human and rat clusters, respectively. The color of the heatmap cells indicates Pearson correlation values between the cluster average expressions. The one-to-one orthologs in the top 2000 highly variable genes of the two maps were used for correlation calculation (see: methods). The comparison indicates a high consistency between the gene expression pattern of hepatic cell types between rats and humans.

Myeloid markers (Clusters 5, 9) – TLH map

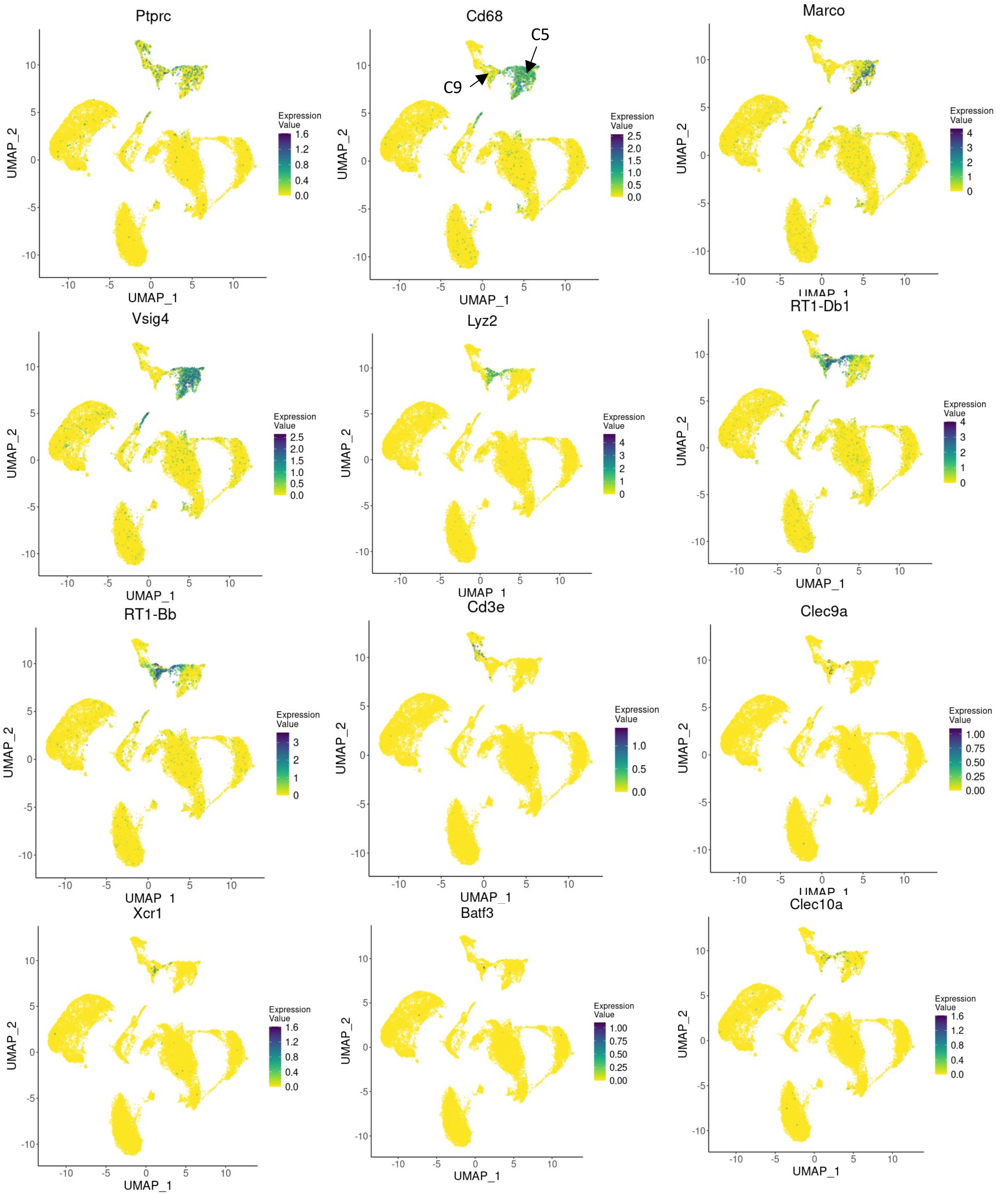


**Supplementary Figure 11: UMAP plots showing the relative distribution of commonly expressed myeloid genes (clusters 5, 9) in the healthy rat total liver homogenate map.** Legend for the relative expression of each marker from lowest expression (yellow dots) to highest expression (dark blue dots) is placed on the right. TLH: total liver homogenate, C: cluster

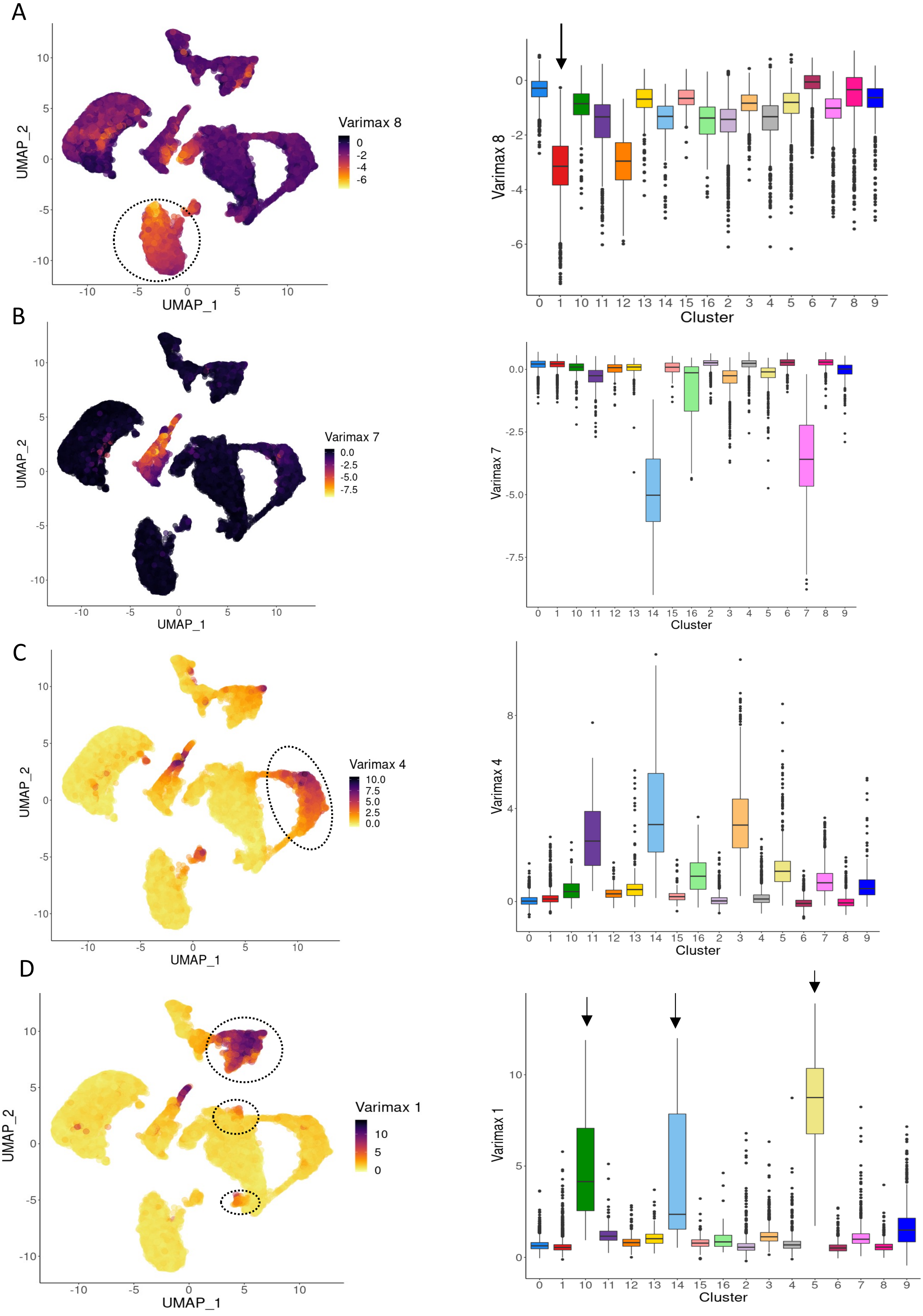
A
UMAP_1
UMAP_2
Varimax 8
Cluster
B
Varimax 7
C
Varimax 4
D
Varimax 1

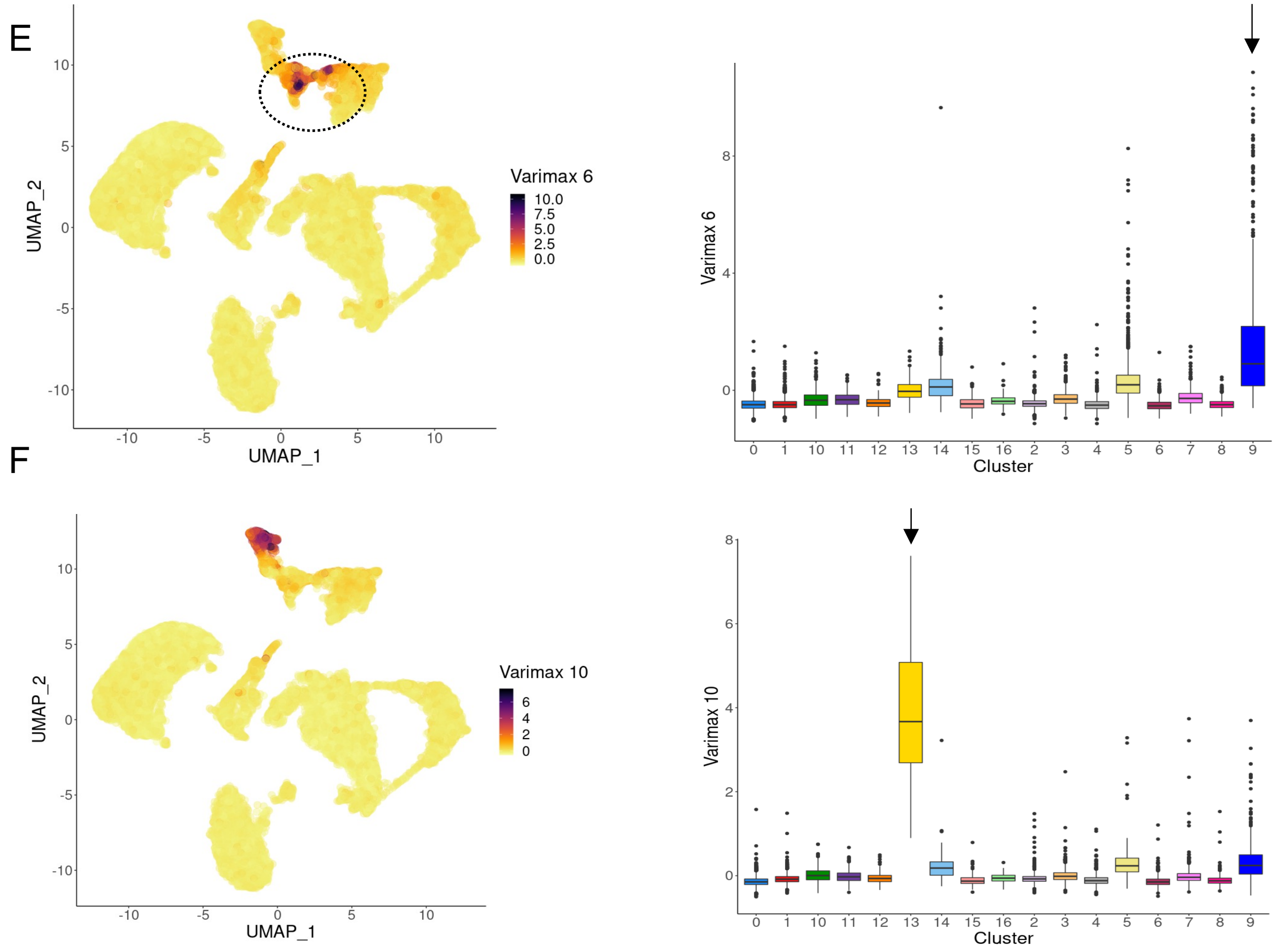


**Supplementary Figure 12: Varimax factors capture the total liver homogenate map cell type signatures.** Overlying A) varimax-8 B) varimax-7 C) varimax-4 D) varimax-1, E) varimax-6, F) varimax-10 scores upon total homogenate map UMAP plot. Legend for the relative score of each cell from low values (yellow dots) to high (dark purple dots). The Boxplots on the right represent the distribution of varimax factors over each cluster.

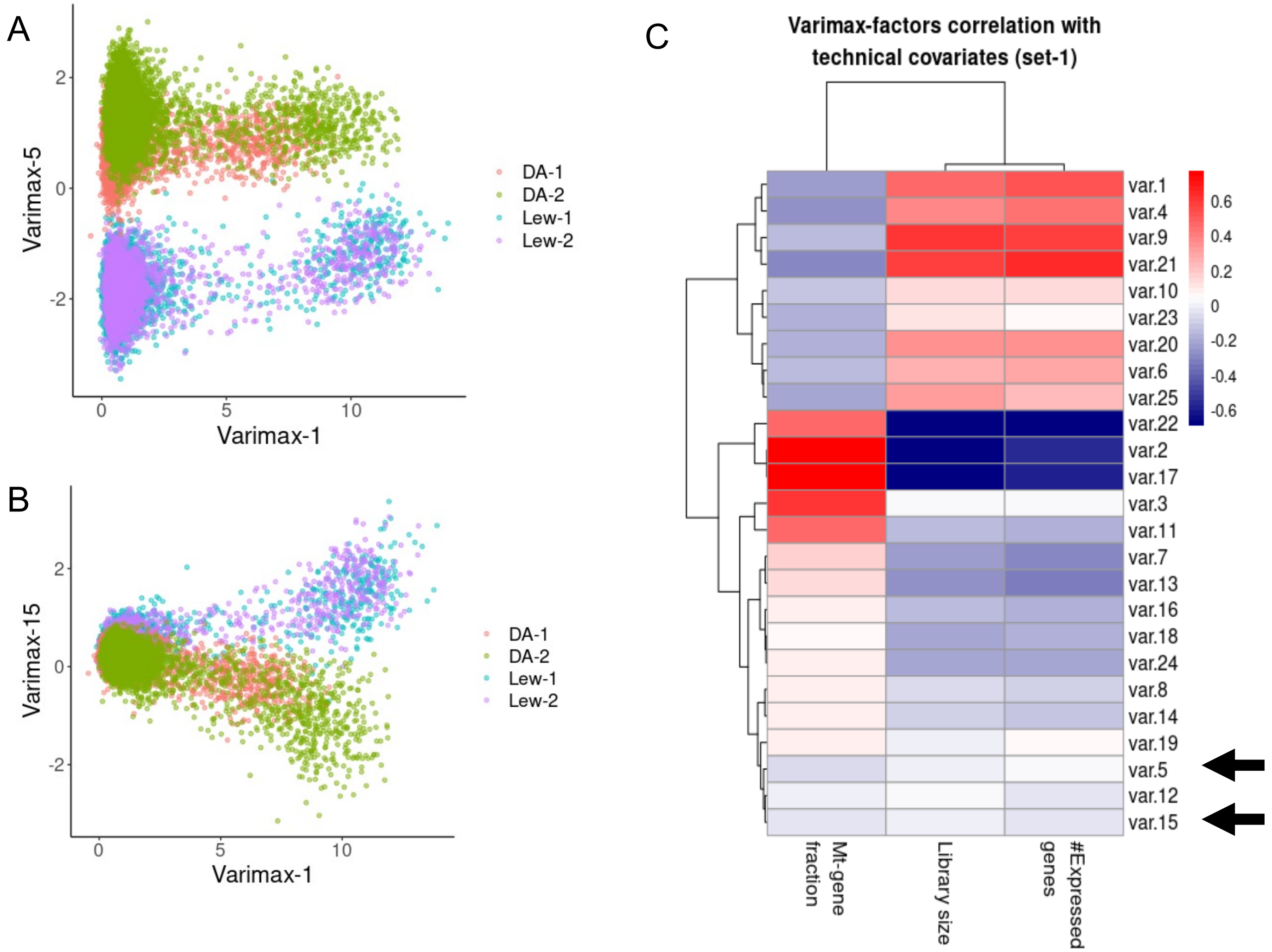


**Supplementary Figure 13: Varimax factors 5 and 15 capture biological, rather than technical variations in the healthy rat liver map.** Figure A and B represent the distribution of rat hepatic cells (total liver homogenate map) over varimax 5/15 (y-axis) and varimax-1 (x-axis). Although all the four samples went through similar sample preparation, cells have been divided based on strain (DA/LEW) and the two samples from the same rat strain are overlapping. Figure C indicates the correlation heatmap of rat liver map (total liver homogenate map) top 25 varimax factors with common technical covariates (Library size, number of expressed genes, and mitochondrial-transcript proportion). The near-zero correlation of varimax 5 and 15 with technical factors suggests that both have been able to capture biological signatures and do not represent technical batch effects. Red and blue indicate positive and negative correlations respectively.

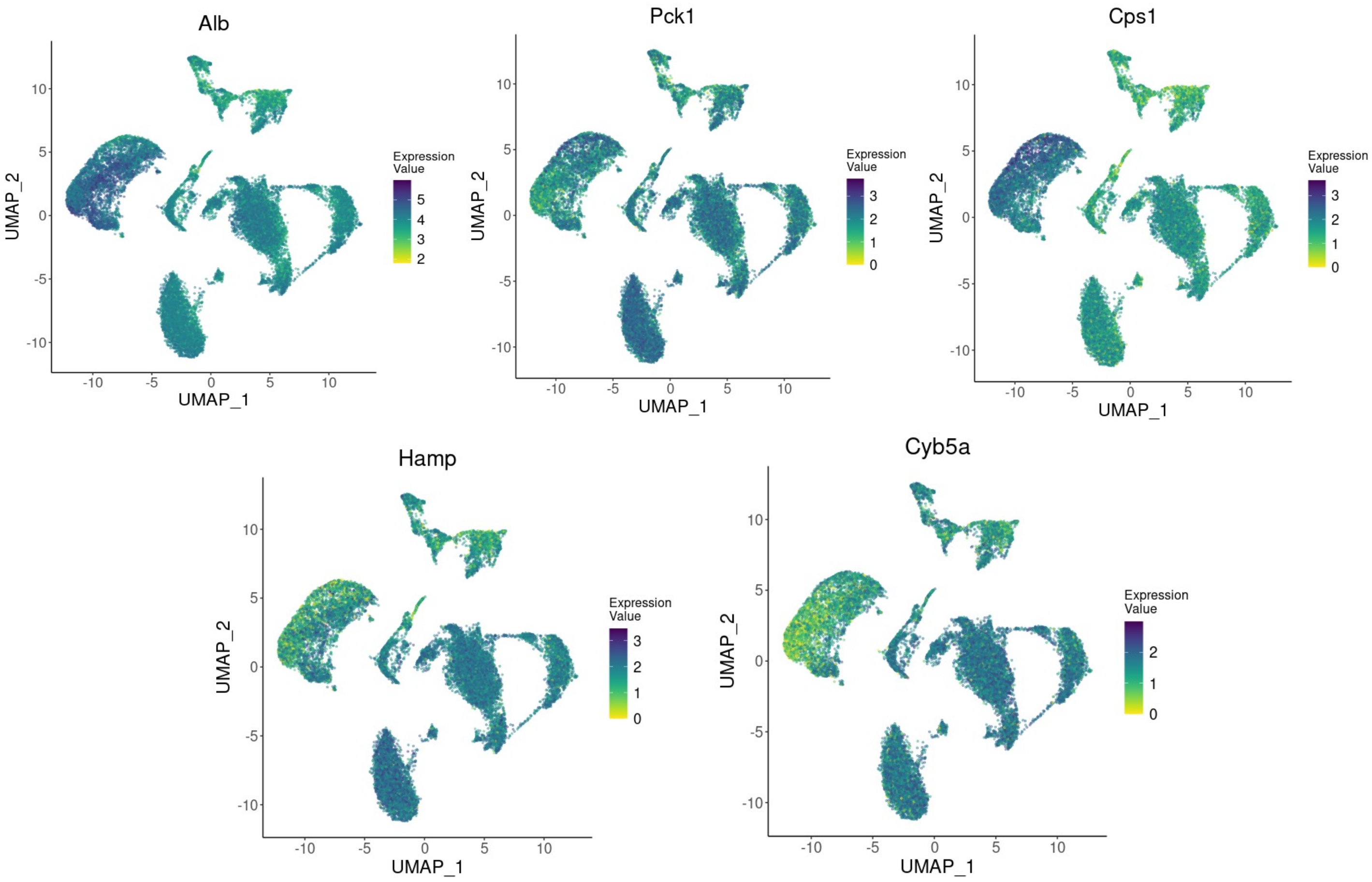


**Supplementary Figure 14: Ambient RNA is released by hepatocyte populations.** High expression values of hepatocyte-specific markers (Alb, Pck1, Cps1, Hamp, Cyb5a) all over the map confirms that the RNA released by fragile hepatocytes is the main source of ambient RNA contamination within the map.

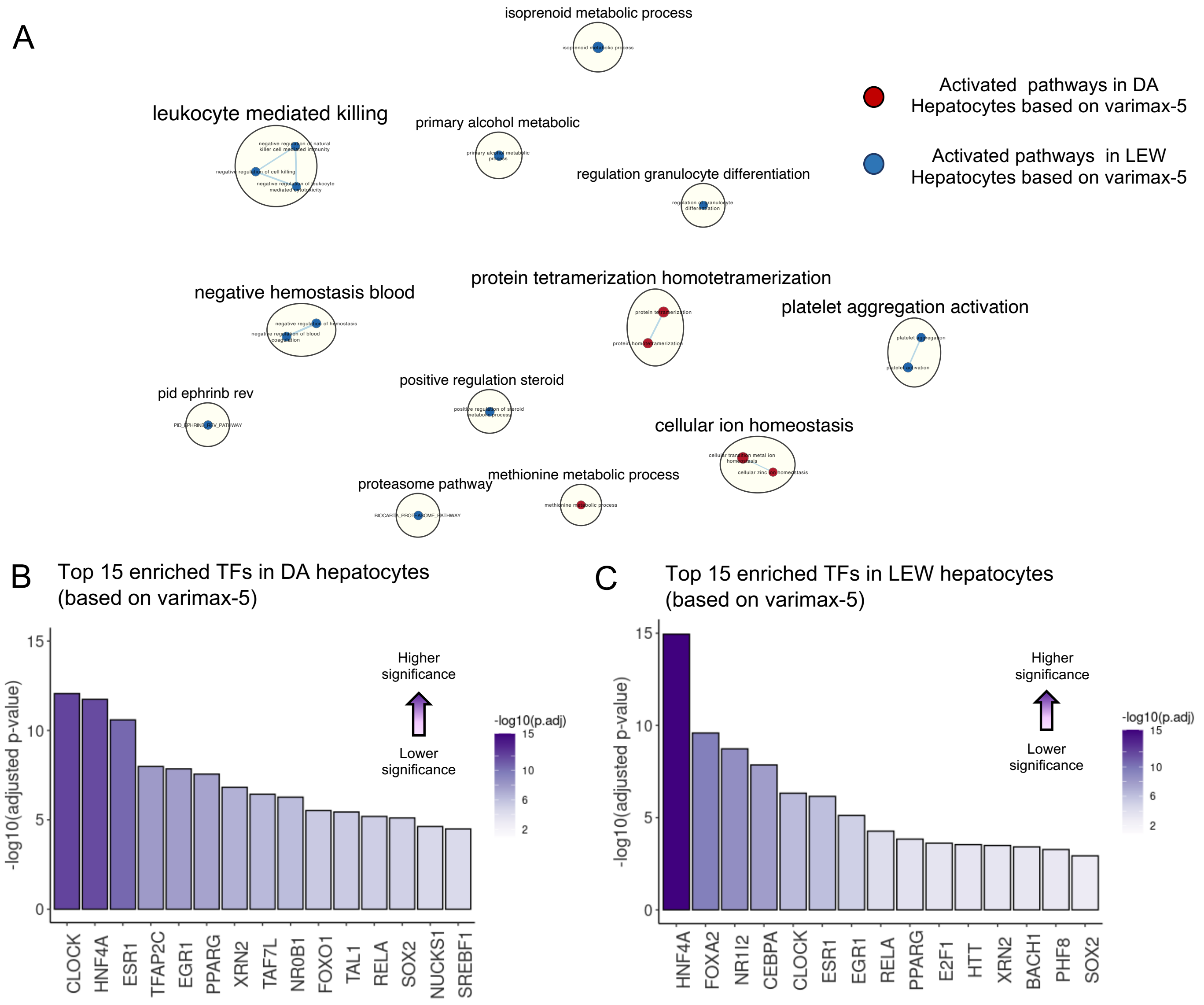


**Supplementary Figure 15: Enrichment analysis of Varimax-5 hepatocyte-related strain-specific differences based on Varimax-5.** A) Pathway enrichment analysis using GSEA to examine active cellular pathways in DA rat vs LEW based on varimax-5 loadings. Each circle represents a GO biological process term. The size of the nodes represents the number of genes in a particular pathway. Since varimax-5 is positively correlated with the DA strain and negatively correlated with LEW, red circles represent up-regulated pathways in the DA rat and blue indicates activated pathways in the Lewis rat. Blue lines depict intra- and inter-pathway relationships according to the number of genes shared between each pathway. Black circles group related pathways into labeled themes. B) Gene-set enrichment analysis using gProfiler on ChIP-Seq-based ChEA dataset to unravel the activated transcription factors (TF) in DA healthy rat liver. Enrichment results have been calculated based on the top 300 genes on the positive side of varimax-5. Activated TFs are sorted based on –log10(adjusted p-value). Dark purple indicates higher significance. C) Gene-set enrichment analysis using gProfiler on ChIP-Seq-based ChEA dataset to unravel the activated transcription factors (TF) in LEW healthy rat liver. Enrichment results have been calculated based on the top 300 genes on the negative side of varimax-5. Activated TFs are sorted based on –log10(adjusted p-value). Dark purple indicates higher significance.

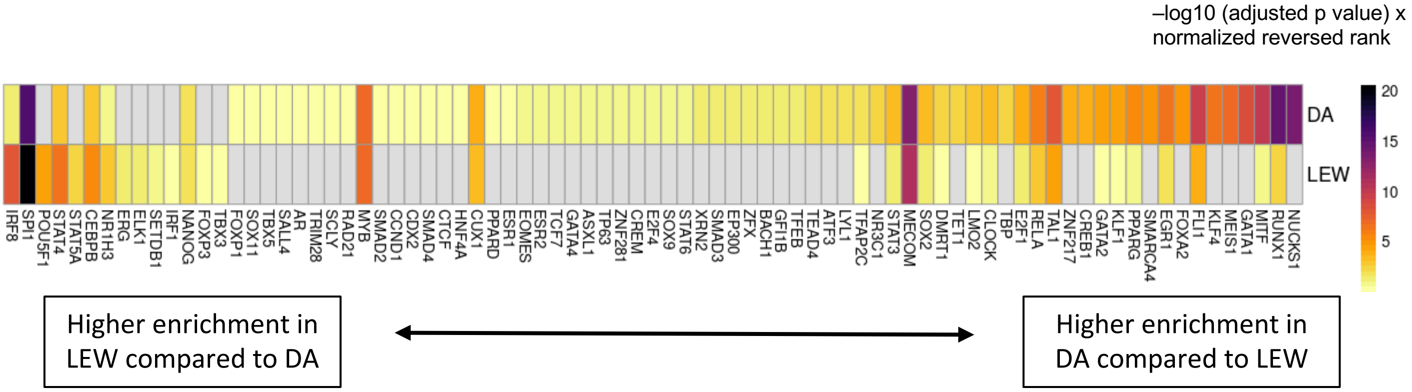


**Supplementary Figure 16: Differential enrichment of strain-specific TFs based on Varimax-15.** Heatmap representing the ranked enrichment of strain-related transcription factors in DA and LEW rat. The TFs on the right, indicate higher enrichment in the DA strain compared to LEW and the TFs on the left represent the reversed enrichment. Significantly enriched TFs in both strains have been scored based on -log10 (adjusted p value) normalized reversed rank. For each TF, enrichment scores in DA and LEW were subtracted from each other and the resulting values were used to sort the TFs. Purple and yellow represent higher and lower enrichment scores, respectively. The grey color indicates that the TF is not enriched in the given strain.

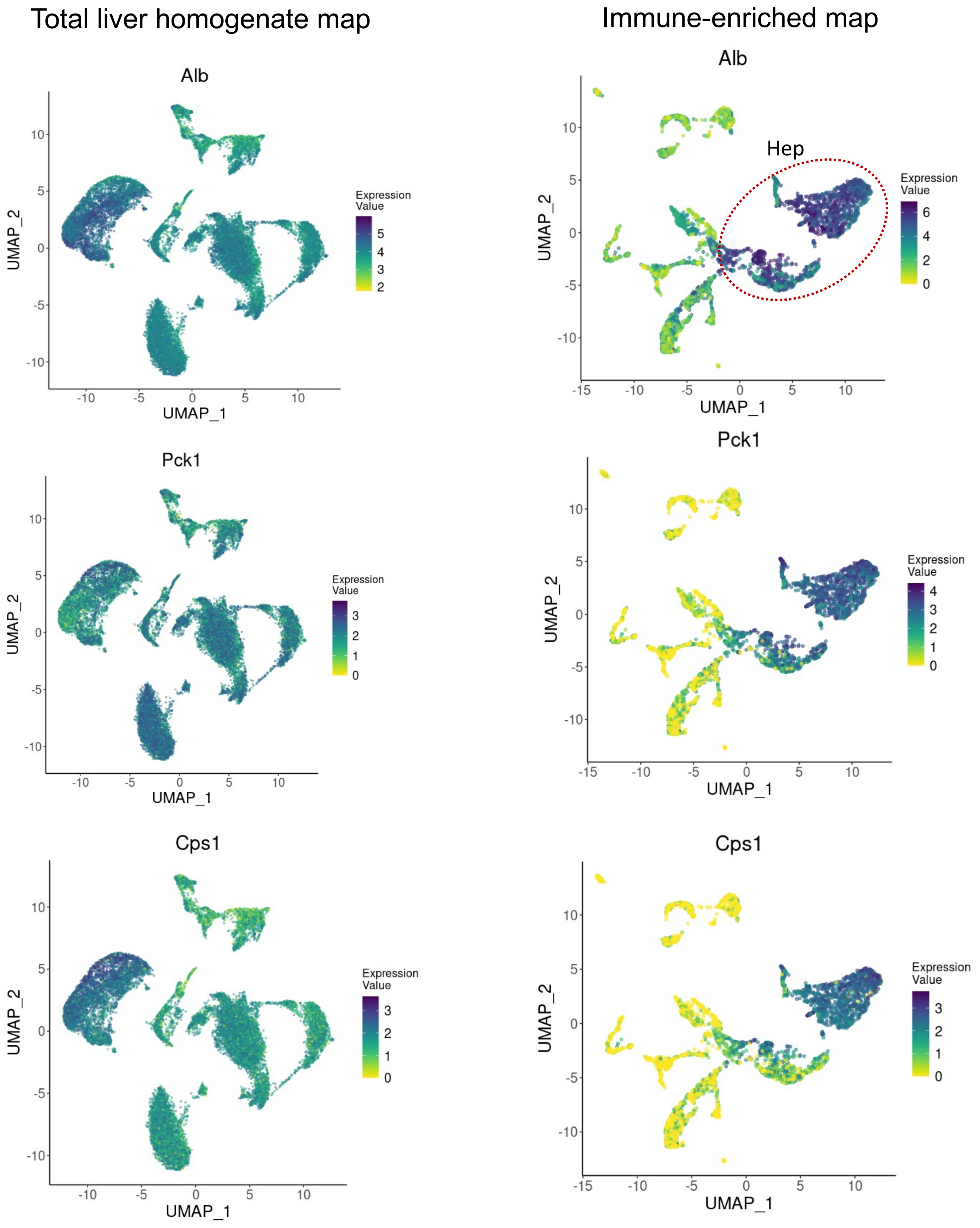


**Supplementary Figure 17: Hepatocyte-released ambient RNA decreased in the immune-enriched map.** Expression distribution of hepatocyte markers Alb, Pck1, and Cps1 in the immune-enriched map and the total liver homogenate map indicates that hepatocyte markers are mainly specific to hepatocyte clusters in the immune-enriched map compared to the total homogenate rat liver atlas**.** This comparison confirms that the level of ambient RNA has been decreased in the immune-enriched map likely due to the additional washing steps.

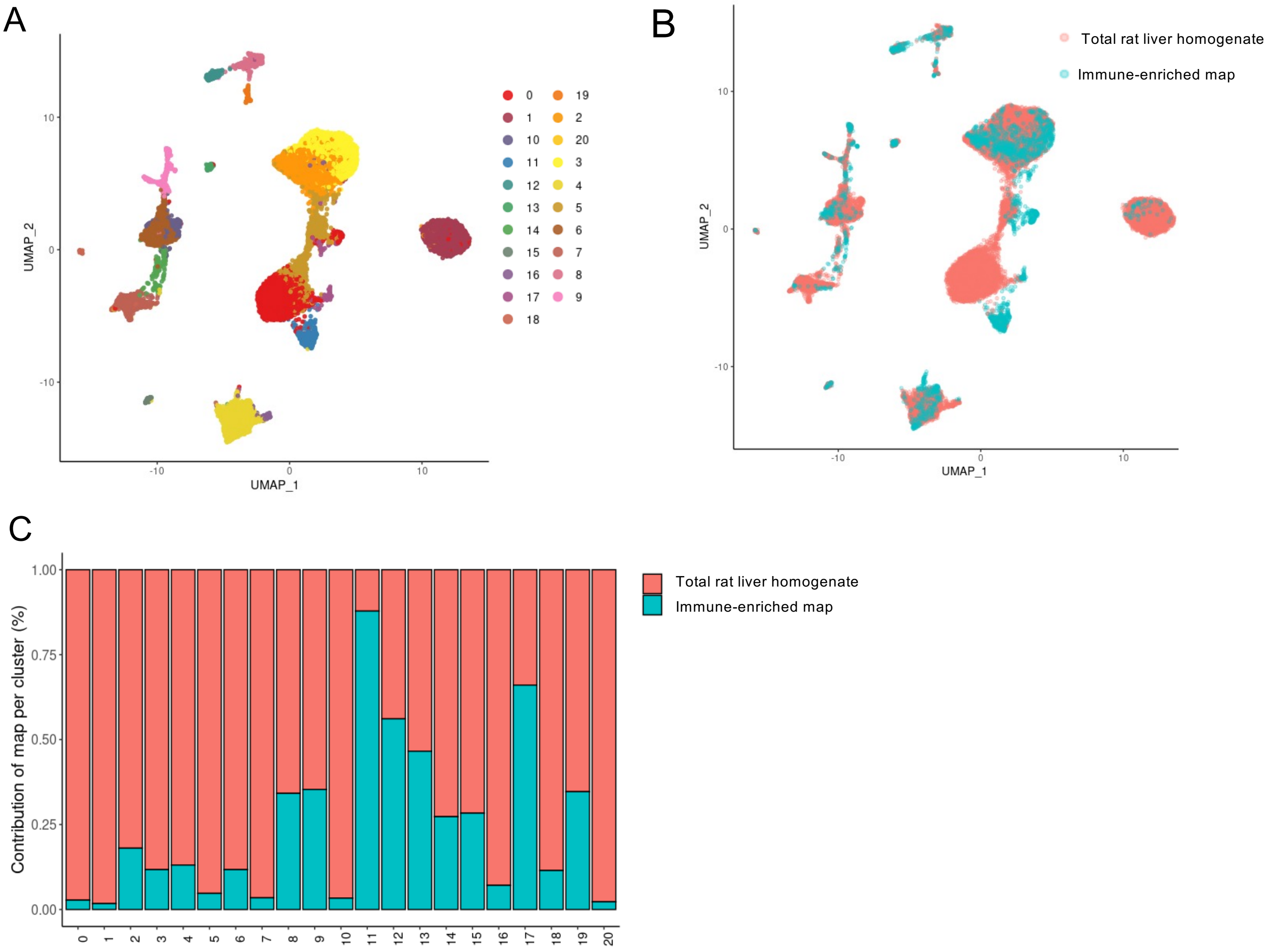


**Supplementary Figure 18: The total liver homogenate and immune-enriched samples could not be well-integrated with standard methods.** The initial four total liver homogenate samples and the two immune-enriched rat livers were merged, batch corrected, and clustered. A) UMAP projection of the merged (total liver homogenate and immune-enriched) rat samples where cells that share similar transcriptome profiles are grouped by colors representing unsupervised clustering results. B) Labeling UMAP projection of cells based on the input sample set indicates that cells from the total liver homogenate and immune-enriched maps do not form well-integrated clusters. C) Bar plot indicating the relative contribution of input samples to each cluster. Although low clustering resolution has been chosen, Clusters 0, 1, 5, 7, 10, and 20 are restricted to cells from the total liver homogenate map and cluster 11 is mainly represented by immune-enriched, indicating that the two maps have not been well-integrated.

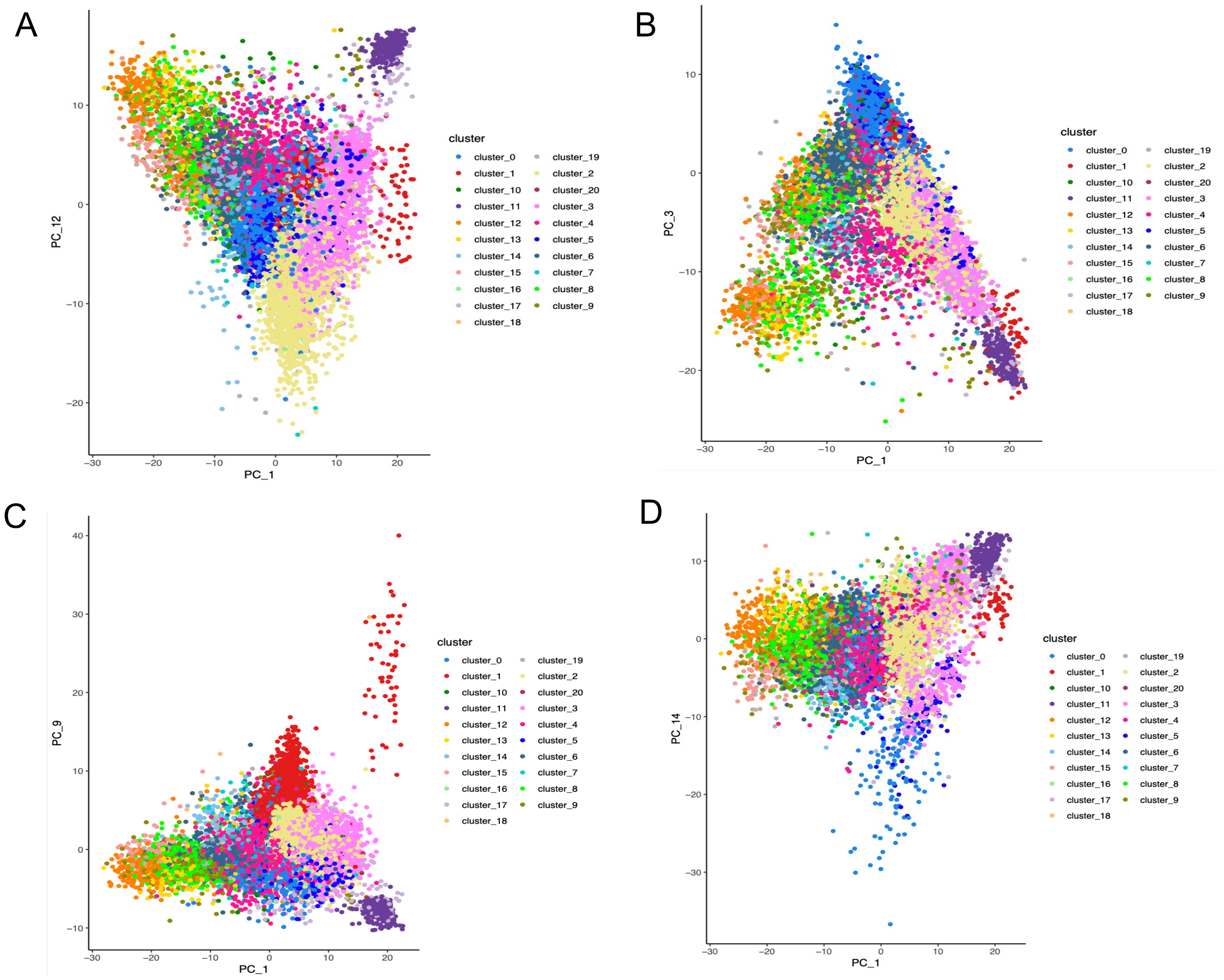


**Supplementary Figure 19: Varimax pipeline is ineffective on the merged total liver homogenate and immune-enriched maps.** The varimax pipeline was applied to the merged total liver homogenate and immune-enriched maps. Distribution of cells over varimax-PC1 (x-axis) and A) varimax-PC12, B) varimax-PC3, C) varimax-PC9 and D) varimax-PC14 on the y-axis are represented as examples. As indicated, it’s not possible to associate the resulting factors with any cluster or covariate and the varimax pipeline has been unable to separate the sources of variation within the merged map into interpretable factors.

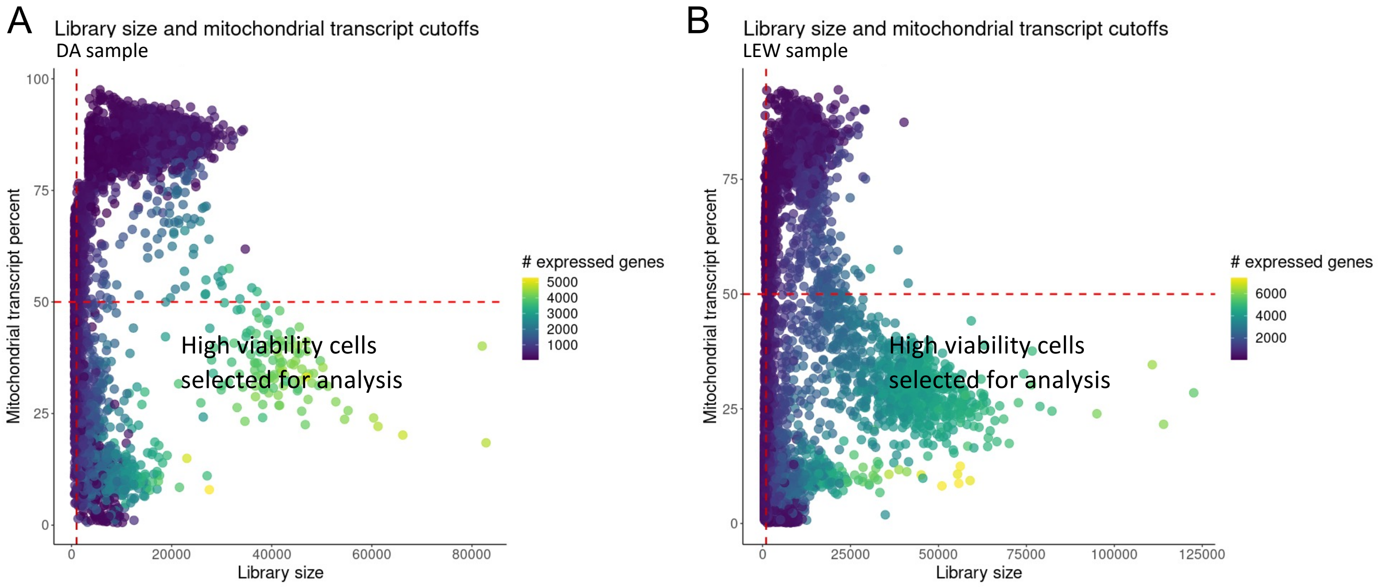


**Supplementary Figure 20: Immune-enriched liver samples quality control and selection of high viability cells.** Viable cells for the immune-enriched map were identified from the single-cell gene-expression data based on having a minimum library size of 1000 transcripts and a maximum of 50% mitochondrial transcript proportion. A) DA B) LEW rat healthy liver sample.

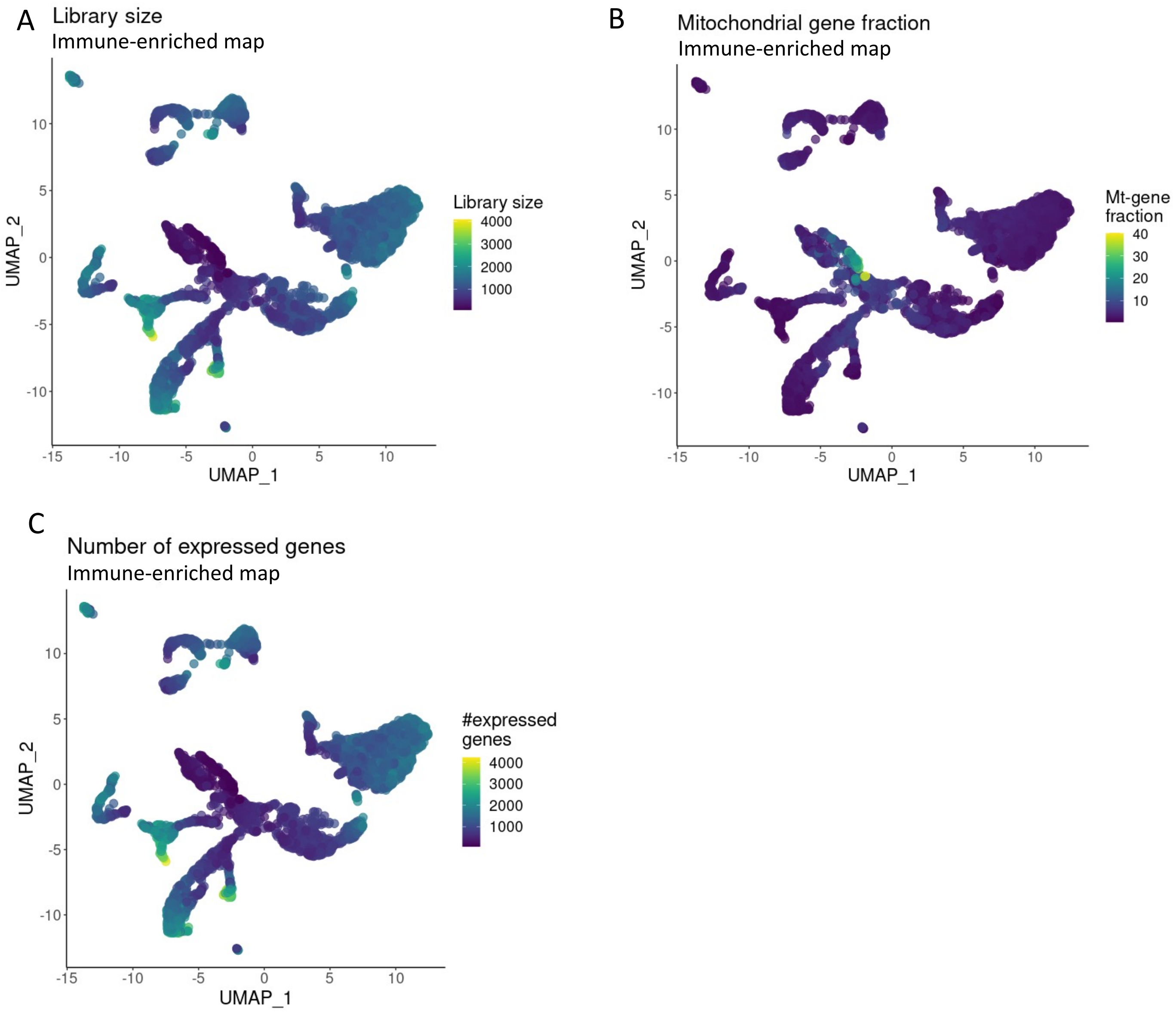


**Supplementary Figure 21: Immune-enriched rat liver map distribution of quality control covariates.** UMAP projection of immune-enriched map where cells are colored based on A) Library size, B) mitochondrial transcript proportion and C) the total number of expressed genes in each cell. Yellow indicates higher values and dark blue indicates lower values of the QC-covariates.

### Cd3⁺ T cell markers (Cluster 10) – Immune-enriched map

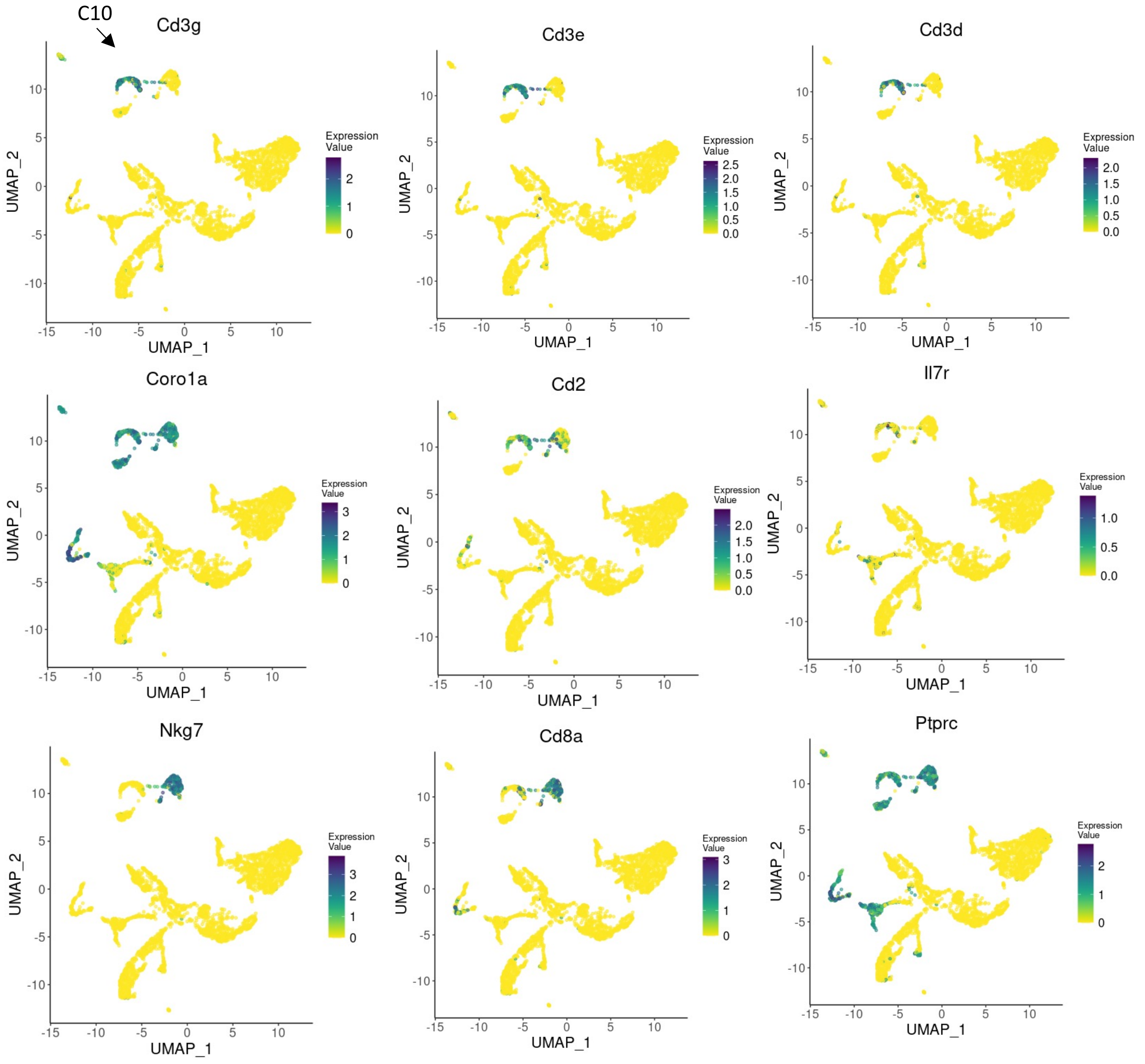


**Supplementary Figure 22: UMAP plots showing the relative distribution of commonly expressed Cd3+ T cell genes (cluster 10) in the healthy rat immune-enriched map.** Legend for the relative expression of each marker from lowest expression (yellow dots) to highest expression (dark blue dots) is placed on the right. C: cluster

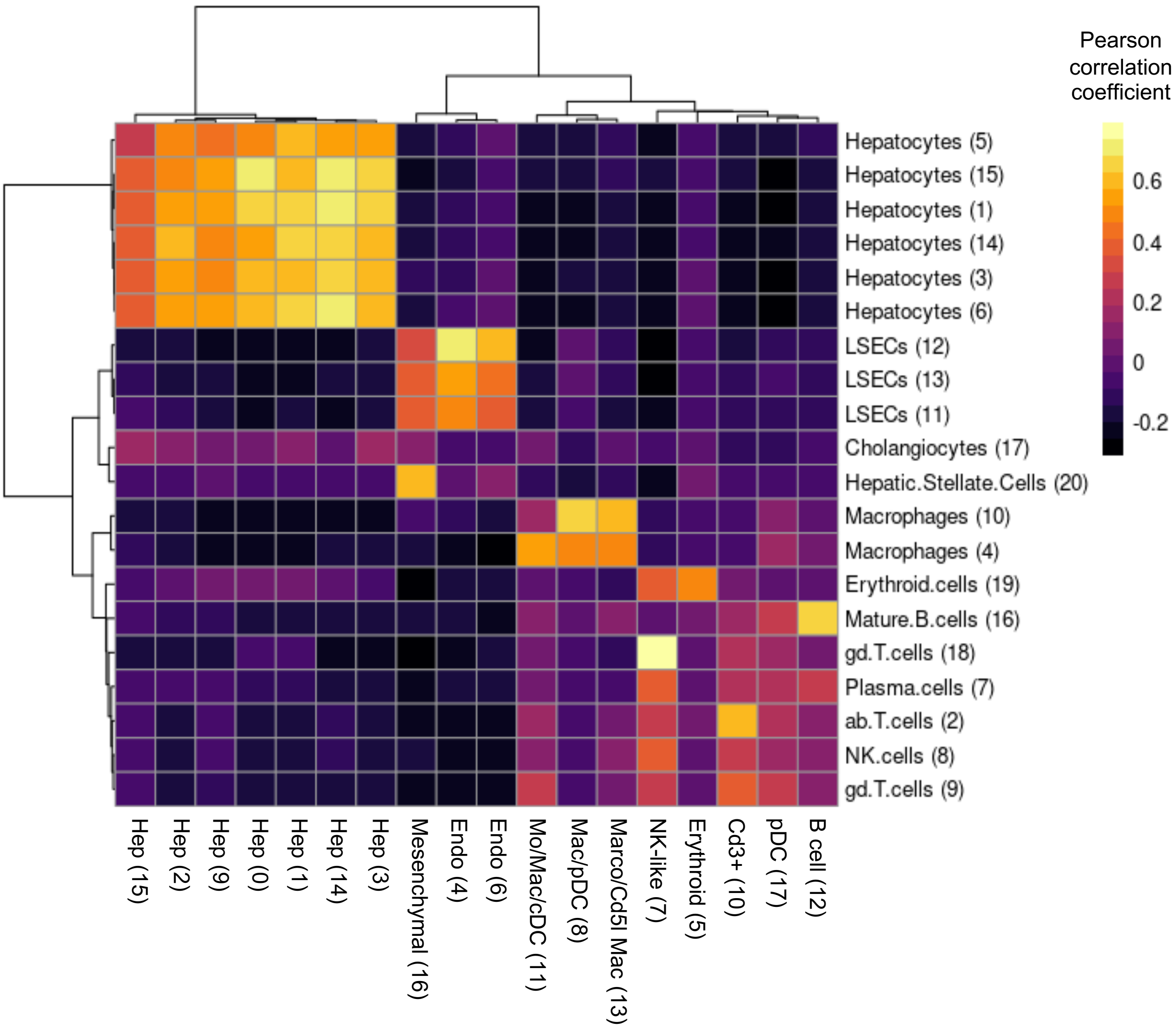


**Supplementary Figure 23: Comparison of rat immune-enriched liver map and human liver atlas.** Comparison of rat immune-enriched liver map and human liver map [Sonya A. MacParland et al.]. Rows and columns of the correlation heatmap represent the human and rat clusters, respectively. The color of the heatmap cells indicates Pearson correlation values between the cluster average expressions. The one-to-one orthologs in the top 2000 highly variable genes of the two maps were used for correlation calculation (see: methods). The comparison indicates a high consistency between the gene expression pattern of hepatic cell types between rats and humans.

B cell markers (Cluster 12) – Immune enriched map

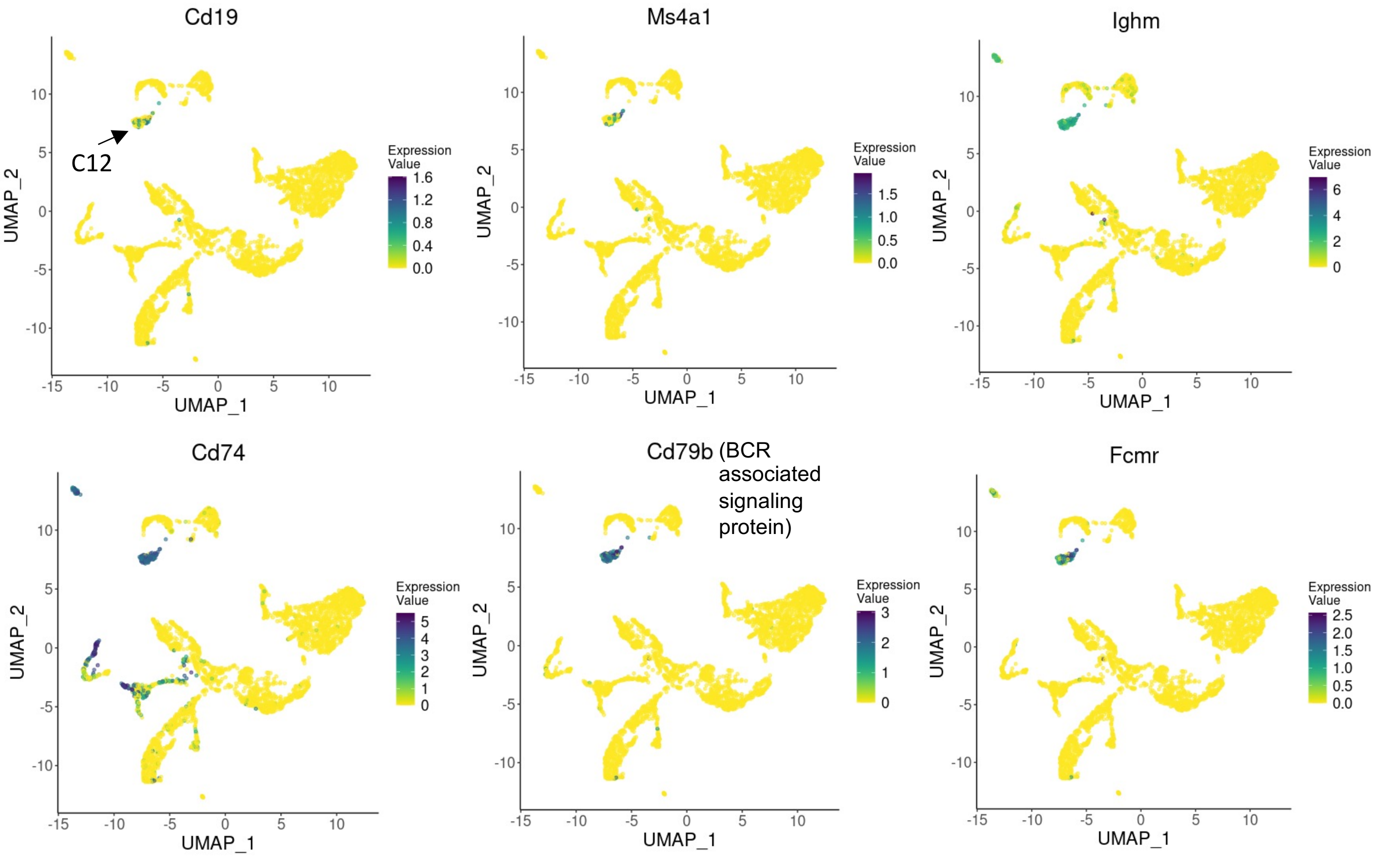


**Supplementary Figure 24: UMAP plots showing the relative distribution of commonly expressed B cell genes (cluster 12) in the healthy rat immune-enriched map.** Legend for relative expression of each marker from lowest expression (yellow dots) to highest expression (Purple dots) is placed on the right. C: cluster pDC-enriched markers (Cluster 17) – Immune enriched map

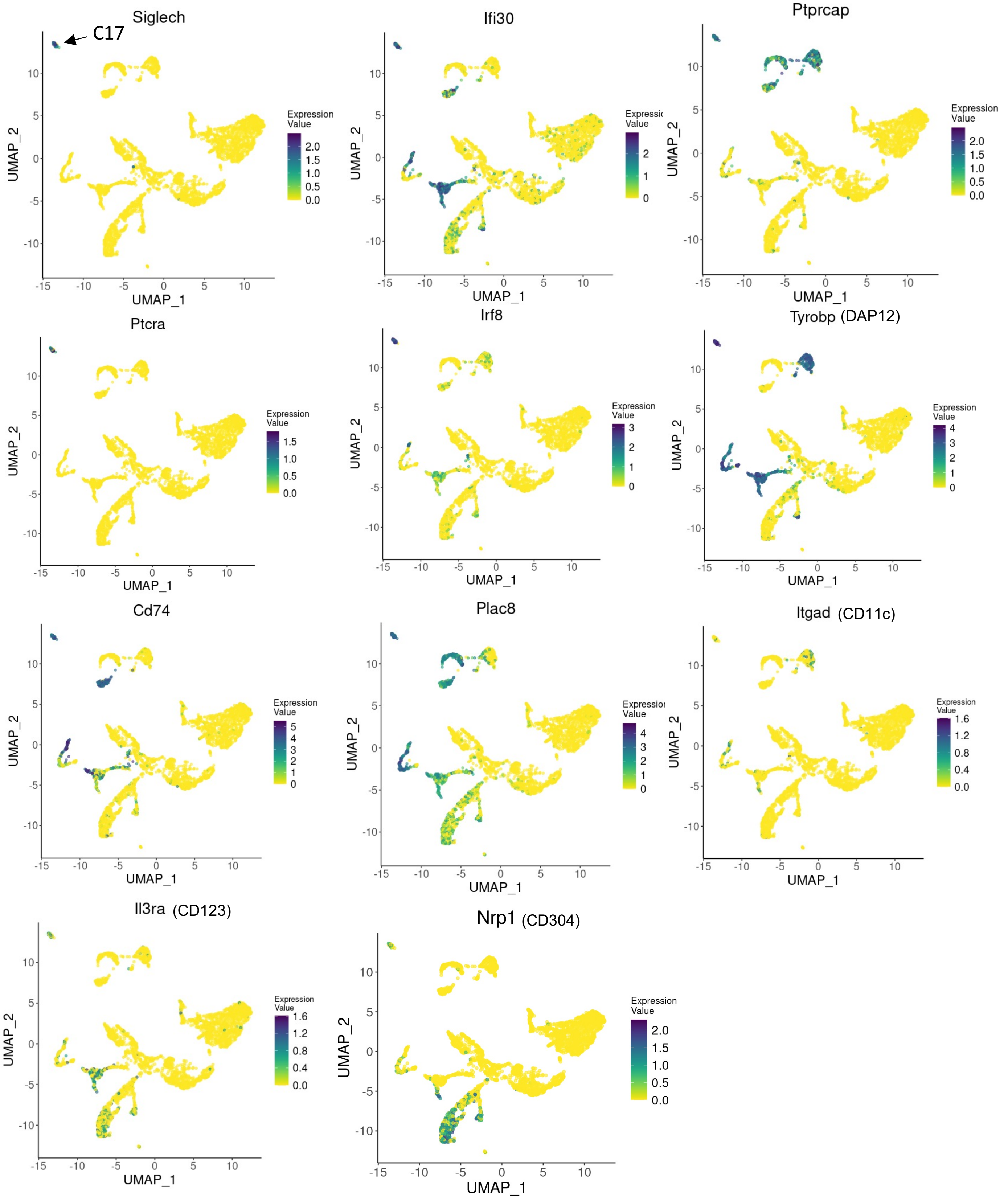


**Supplementary Figure 25: UMAP plots showing the relative distribution of commonly expressed pDC genes (cluster 17) in the healthy rat total liver homogenate map.** Legend for the relative expression of each marker from lowest expression (yellow dots) to highest expression (dark blue dots) is placed on the right. C: cluster

NK-like cell markers (Cluster 7) – Immune enriched map

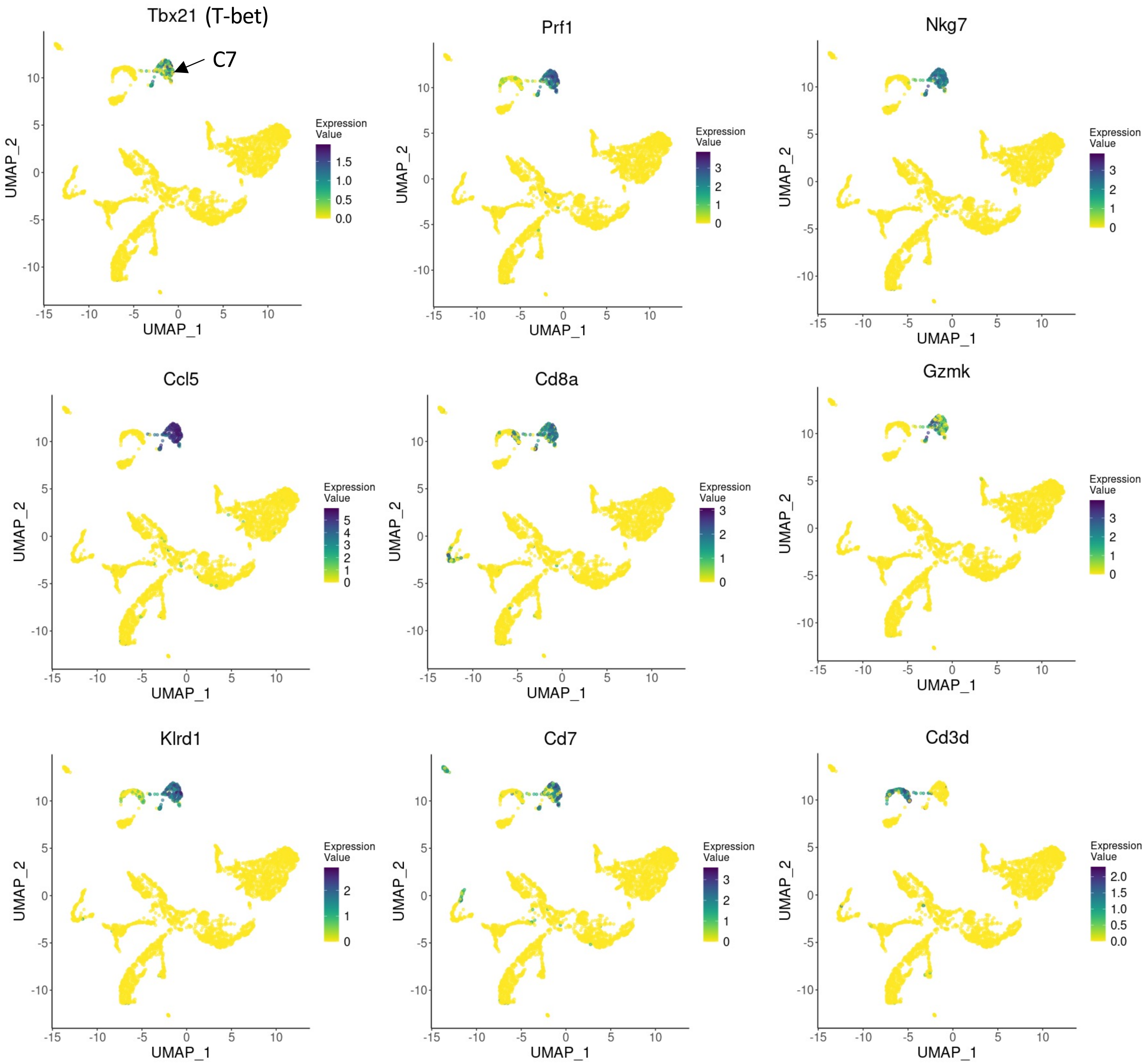


**Supplementary Figure 26: UMAP plots showing the relative distribution of commonly expressed NK-like cell genes (cluster 7) in the healthy rat total liver homogenate map.** Legend for the relative expression of each marker from lowest expression (yellow dots) to highest expression (Purple dots) is placed on the right. C: cluster cDC cell markers (part of cluster 11) – Immune enriched map

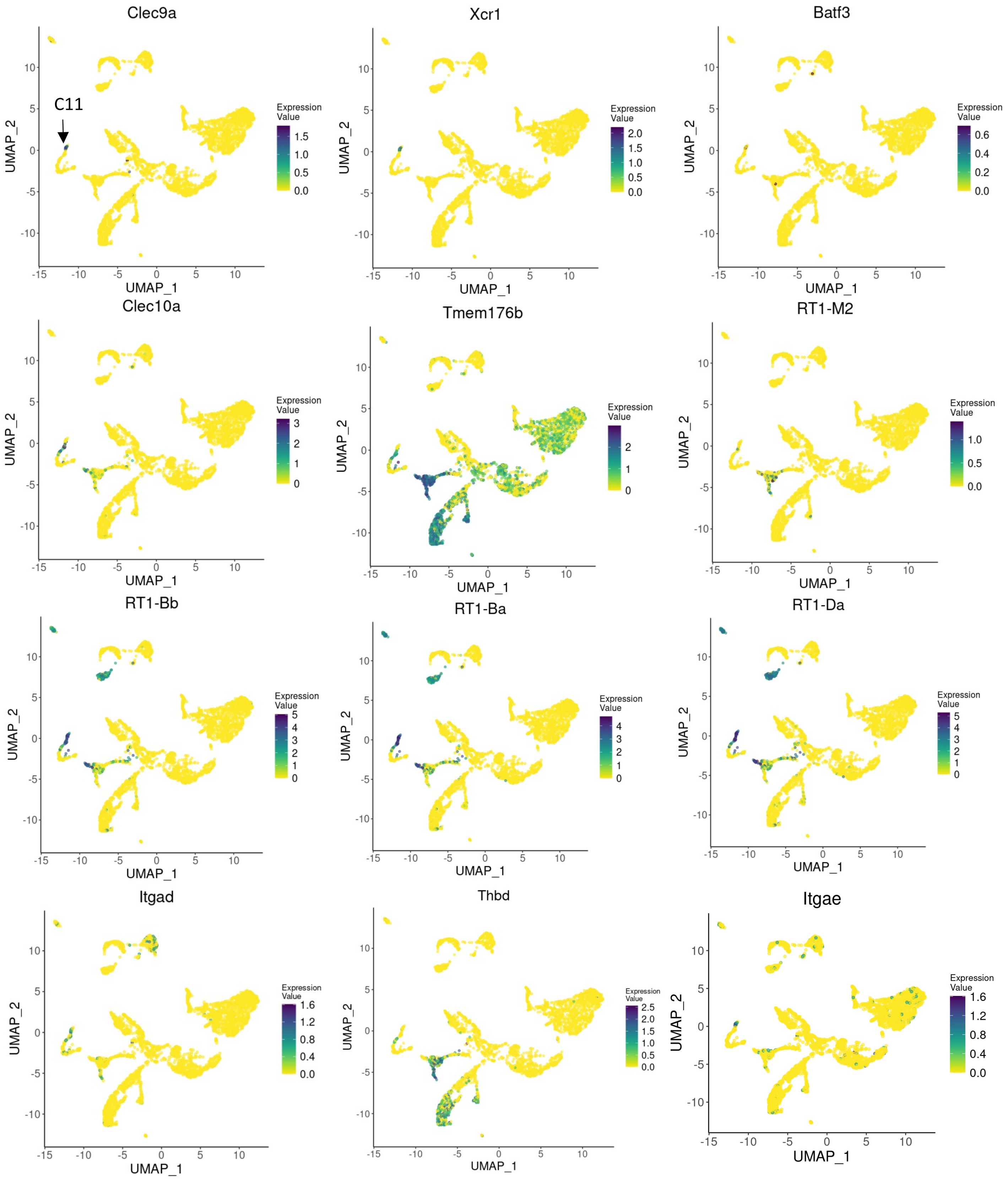


**Supplementary Figure 27: UMAP plots showing the relative distribution of commonly expressed cDC genes (cluster 11) in the healthy rat total liver homogenate map.** Legend for the relative expression of each marker from lowest expression (yellow dots) to highest expression (dark blue dots) is placed on the right. C: cluster

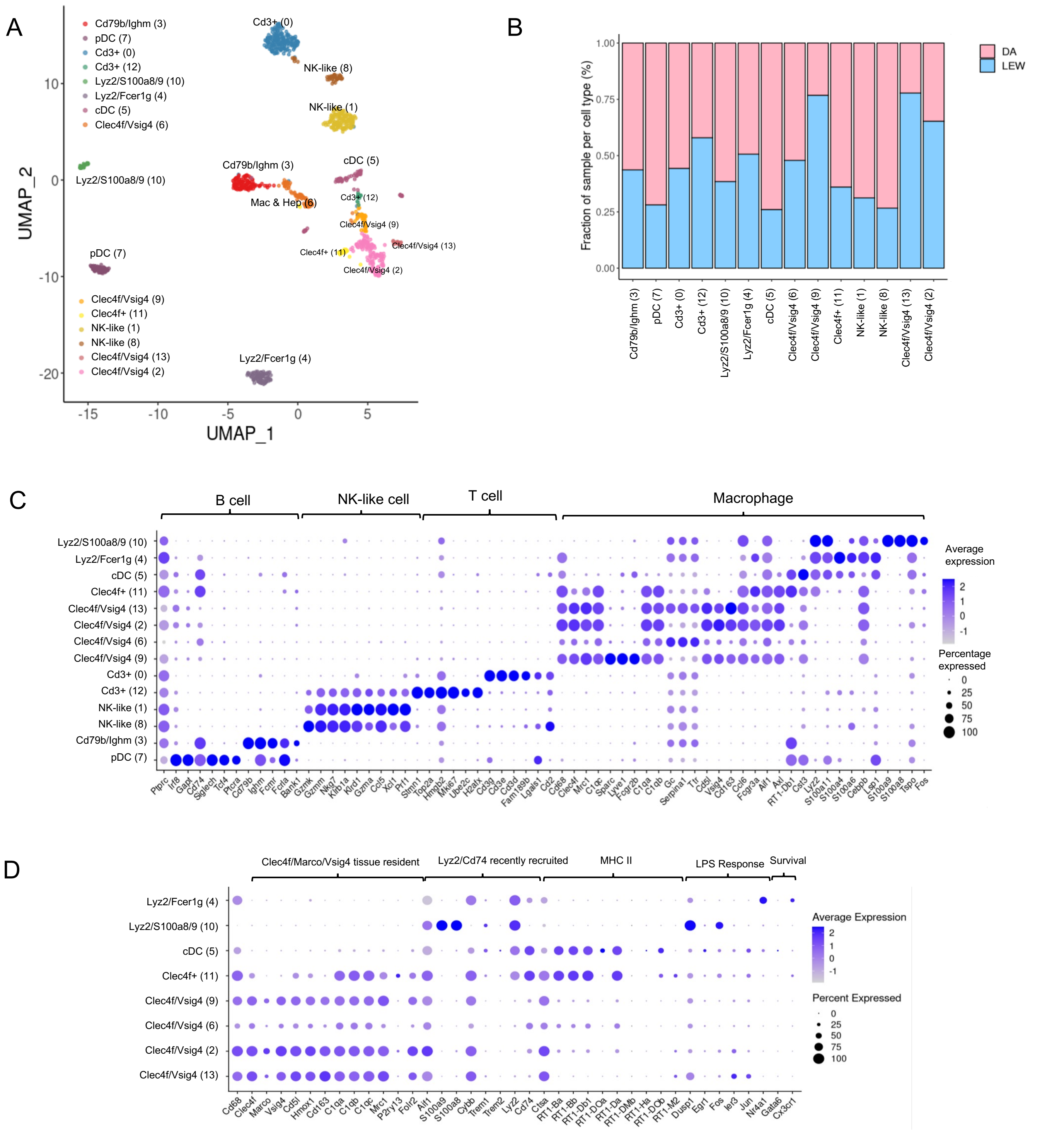
A
Cd79b/Ighm (3)
pDC (7)
Cd3+ (0)
Cd3+ (12)
Lyz2/S100a8/9 (10)
Lyz2/Fcer1g (4)
cDC (5)
Clec4f/Vsig4 (6)
Clec4f/Vsig4 (9)
Clec4f+ (11)
NK-like (1)
NK-like (8)
Clec4f/Vsig4 (13)
Clec4f/Vsig4 (2)
Mac & Hep (6)
UMAP_1
UMAP_2
B
Fraction of sample per cell type (%)
DA
LEW
C
B cell
NK-like cell
T cell
Macrophage
Average expression
Percentage expressed
Ptprc Irf8 Gapt Cd74 Siglech Tcf4 Ptcra Cd79b Ighm Fcmr Fcrla Bank1 Gzmk Gzmm Nkg7 Klrb1a Klrd1 Gzma Ccl5 Xcl1 Prf1 Stmn1 Top2a Hmgb2 Mki67 Ube2c H2afx Cd3g Cd3e Cd3d Fam189b Lgals1 Cd2 Cd68 Clec4f Mrc1 C1qc Sparc Lyve1 Fcgr2b C1qa C1qb Gc Serpina1 Ttr Cd5l Vsig4 Cd163 Ccl6 Fcgr3a Aif1 Axl RT1-Db1 Cst3 Lyz2 S100a11 S100a4 S100a6 Cebpb Lsp1 S100a9 S100a8 Tspo Fos
D
Clec4f/Marco/Vsig4 tissue resident
Lyz2/Cd74 recently recruited
MHC II
LPS Response
Survival
Average Expression
Percent Expressed
Cd68 Clec4f Marco Vsig4 Cd5l Hmox1 Cd163 C1qa C1qb C1qc Mrc1 P2ry13 Folr2 Aif1 S100a9 S100a8 Cybb Trem1 Trem2 Lyz2 Cd74 Ctsa RT1-Ba RT1-Bb RT1-Db1 RT1-DOa RT1-Da RT1-DMb RT1-Ha RT1-DOb RT1-M2 Dusp1 Egr1 Fos Ier3 Jun Nr4a1 Gata6 Cx3cr1

**Supplementary Figure 28: Subclustering of *Ptprc*+ cell types indicates 14 immune cell populations.** A) UMAP projection of the *Ptprc*+ subpopulation of the immune-enriched samples. The *Ptprc*+ clusters of the immune-enriched map were subclustered to provide a more detailed representation of immune subtypes. Colors indicate different subclusters. B) Bar plot indicating the relative contribution of input samples to each subcluster. All samples have been represented in each of the subclusters. C) Dot-plot indicating the relative expression of marker genes in each immune subcluster. Evaluation of the top markers subclusters 6 and 9 indicate that a few LSECs and hepatocytes might have been mixed within the immune clusters. The x-axis represents marker genes, and the y-axis represents the annotated subclusters within the map. The size of the circle indicates the percentage of cells in each population which express the marker of interest and the color indicates the average expression value (dark purple: high expression, grey: low expression). D) Dot plot demonstrating the expression pattern of different macrophage markers. Macrophage markers have been grouped into 6 categories (non-inflammatory, inflammatory, MHC II, LPS response, and survival) based on their potential role in macrophage function. The interpretation of the size and color of the circles is the same as in previous dot plots.

T cell markers (subcluster 0, 12) – Immune subclustering

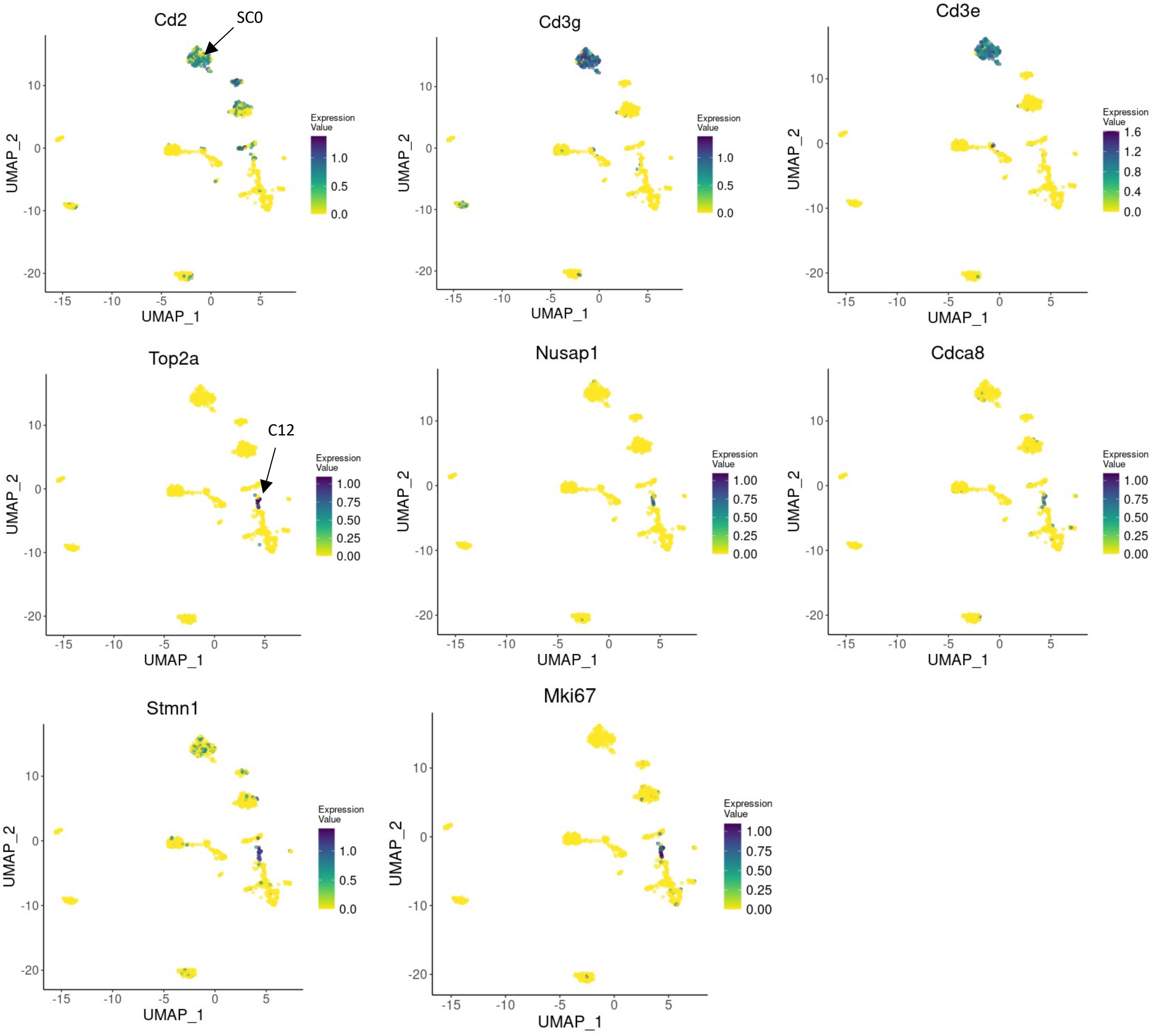


**Supplementary Figure X: UMAP plots showing the relative distribution of commonly expressed T cell markers (subclusters 0, 12) in the healthy rat immune population subclustering map.** Legend for the relative expression of each marker from lowest expression (yellow dots) to highest expression (dark blue dots) is placed on the right. SC: subcluster

NK-like cell markers (subcluster 1, 8) – Immune subclustering

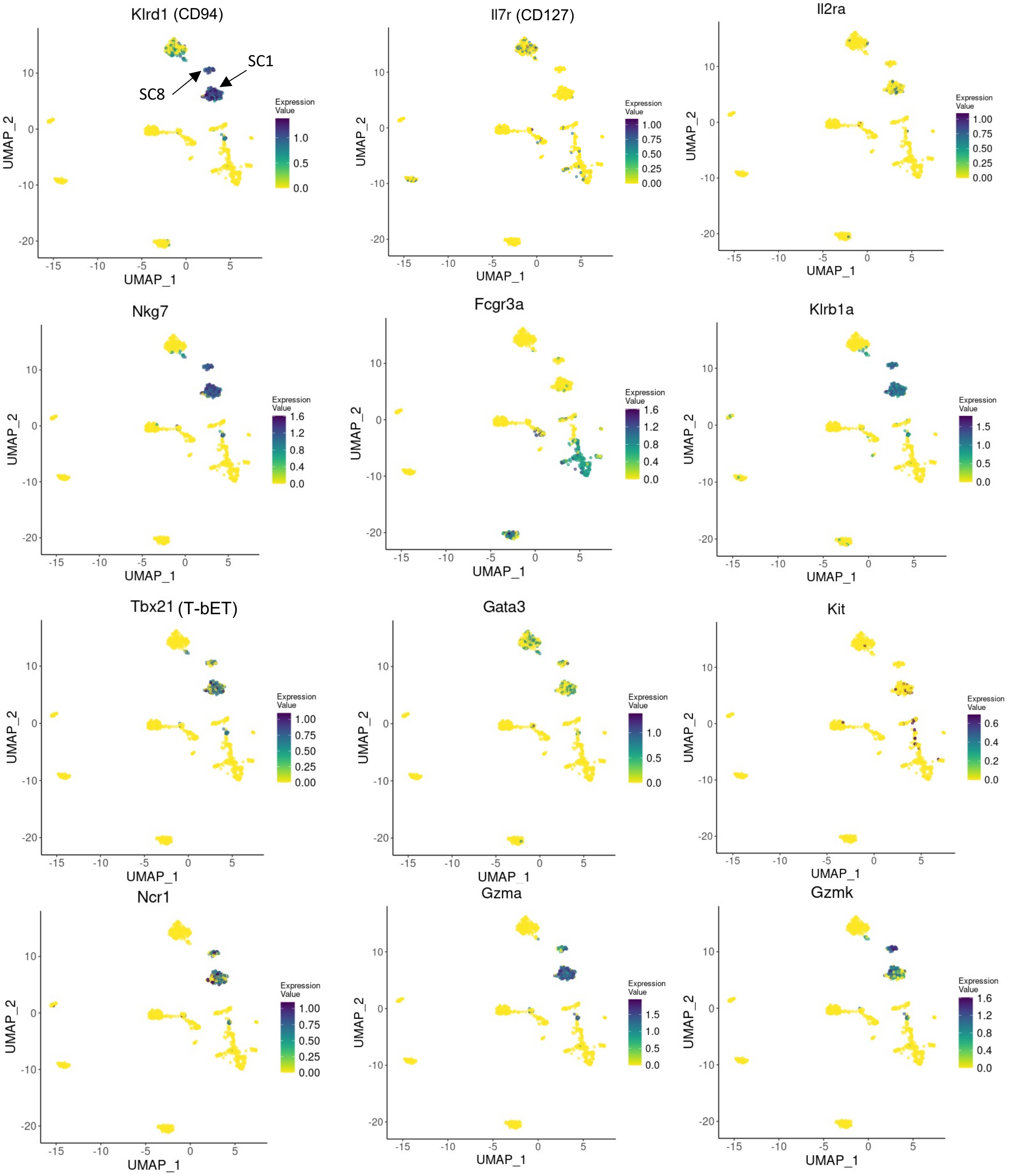


**Supplementary Figure X: UMAP plots showing the relative distribution of commonly expressed NK-like cell markers (subclusters 1, 8) in the healthy rat immune population subclustering map.** Legend for the relative expression of each marker from lowest expression (yellow dots) to highest expression (Purple dots) is placed on the right. SC: subcluster

B cell markers (subcluster 3) – Immune subclustering

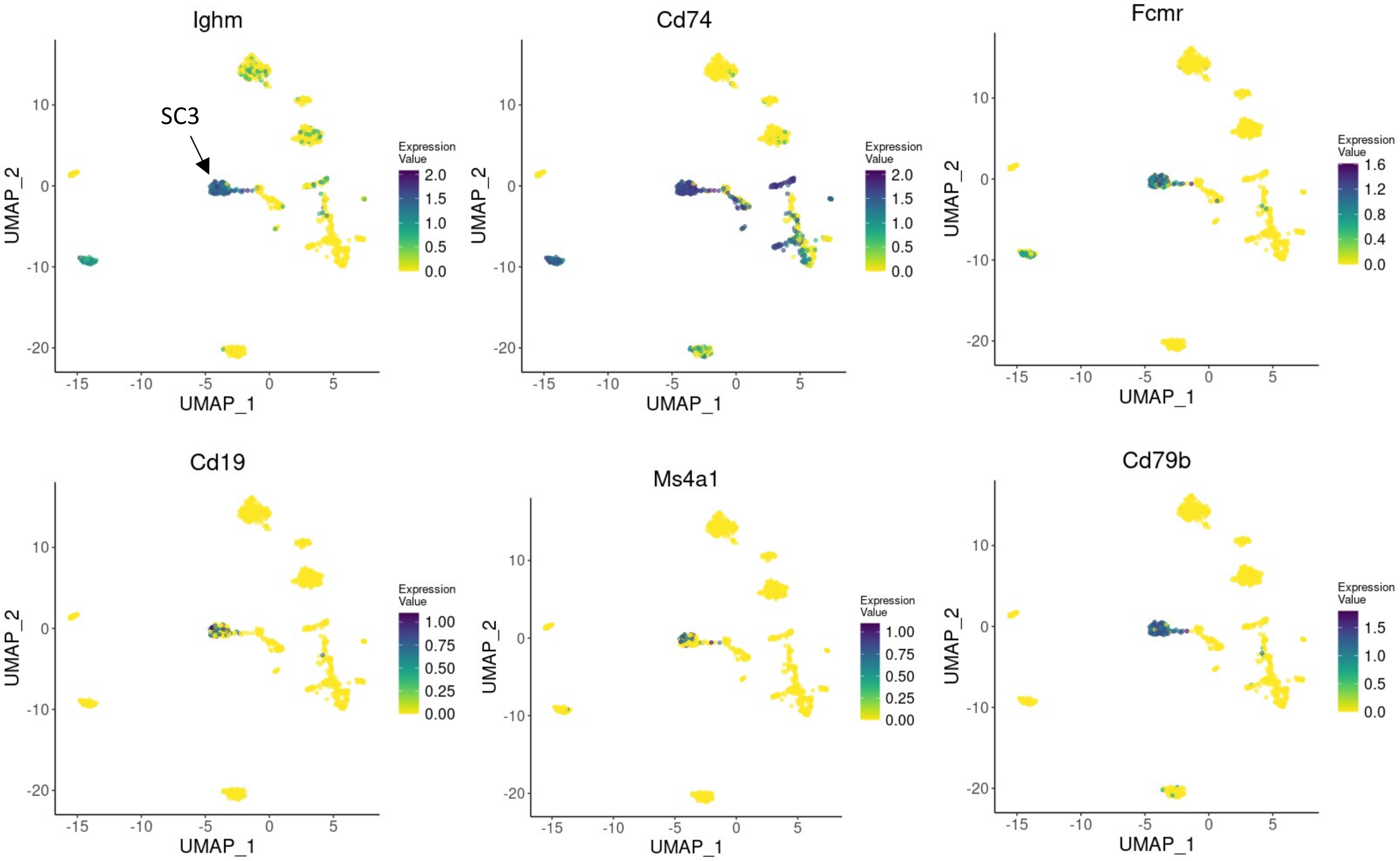


**Supplementary Figure X: UMAP plots showing the relative distribution of commonly expressed B cell markers (subcluster 3) in the healthy rat immune population subclustering map.** Legend for the relative expression of each marker from lowest expression (yellow dots) to highest expression (dark blue dots) is placed on the right. SC: subcluster

Myeloid cells markers (subcluster 2, 4, 6, 9, 11, and 13) – Immune subclustering

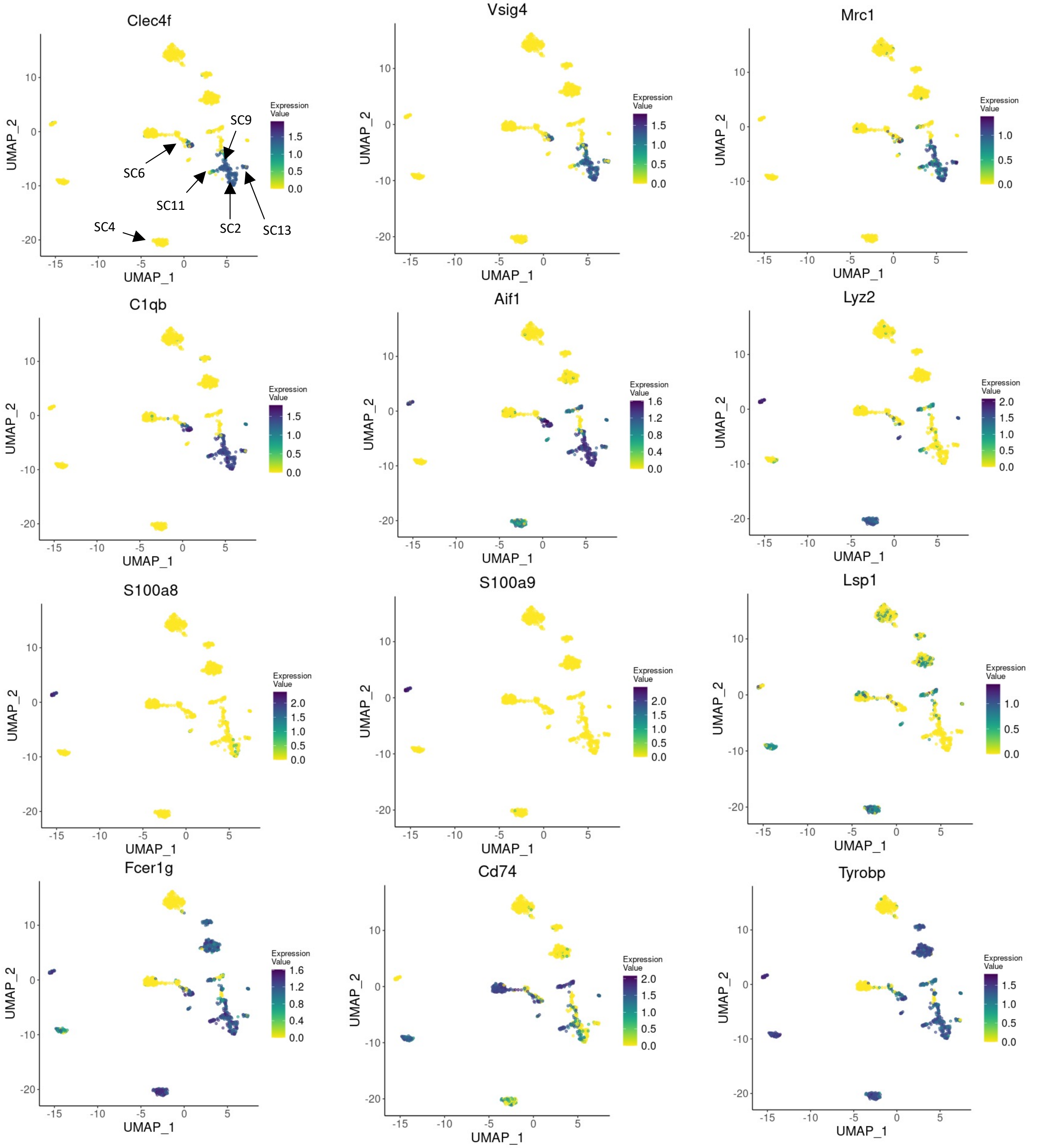


**Supplementary Figure X: UMAP plots showing the relative distribution of commonly expressed Myeloid cell markers (subcluster 2, 4, 6, 9, 11, and 13) in the healthy rat immune population subclustering map.** Legend for the relative expression of each marker from lowest expression (yellow dots) to highest expression (purple dots) is placed on the right. SC: subcluster

### pDC and cDC markers (subcluster 5,7) – Immune subclustering

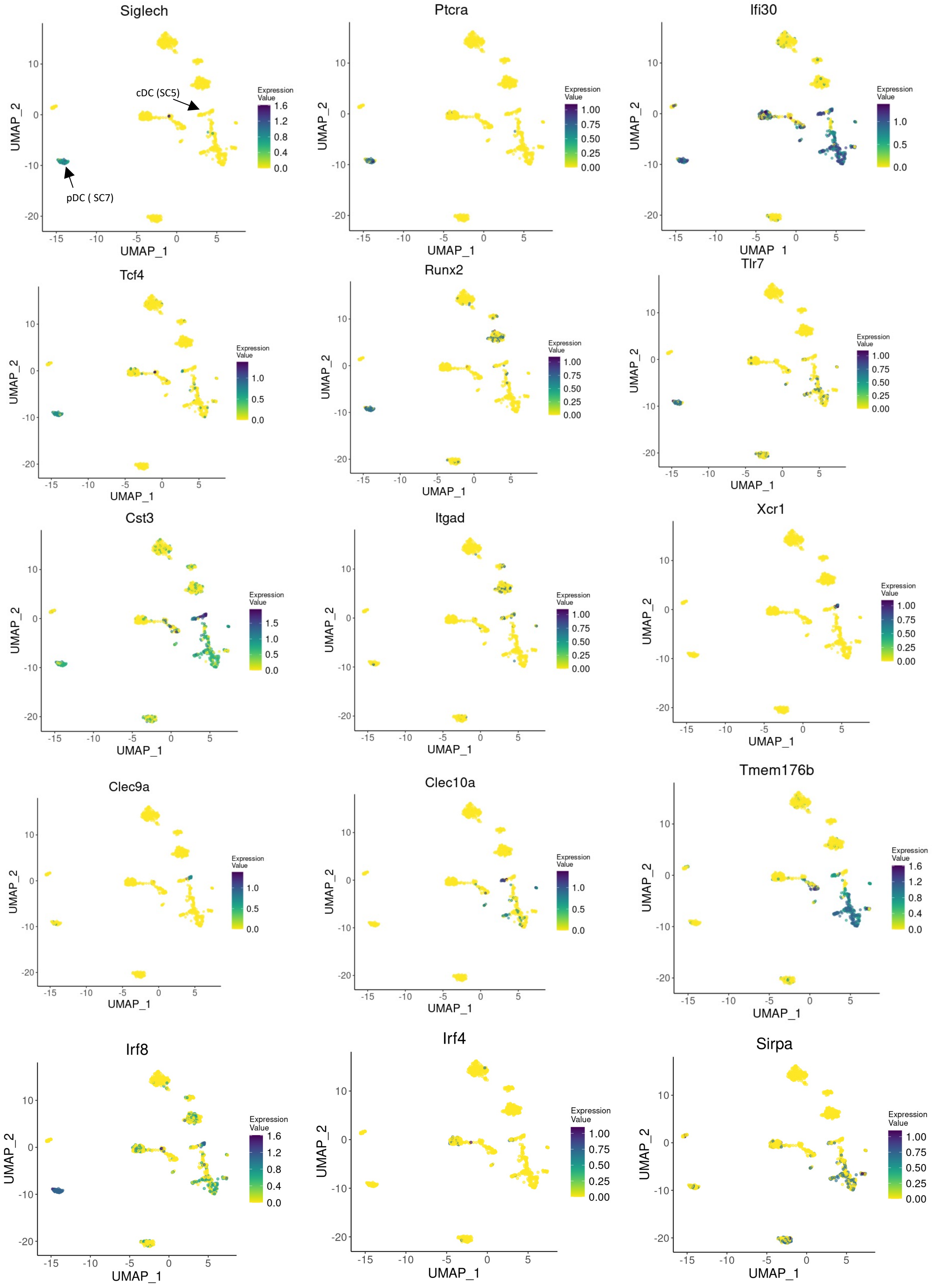

**Supplementary Figure X: UMAP plots showing the relative distribution of commonly expressed pDC and cDC markers (subcluster 5, 7) in the healthy rat immune population subclustering map.** Legend for the relative expression of each marker from lowest expression (yellow dots) to highest expression (dark blue dots) is placed on the right. SC: subcluster

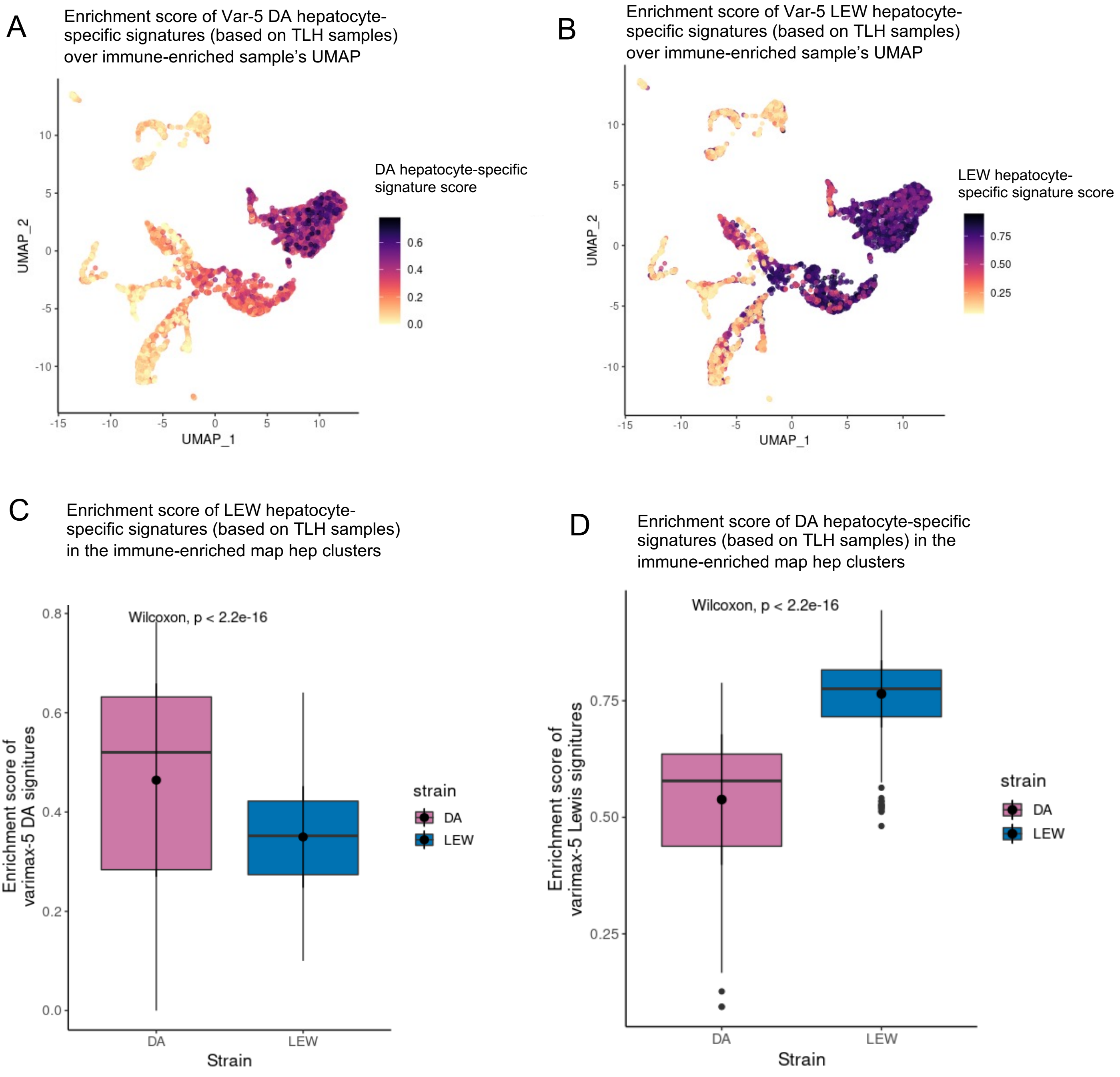

**Supplementary Figure 29) The hepatocyte-specific strain variations in the total liver map are verified in the immune-enriched map.** To verify the hepatocyte-specific strain variations (captured by varimax-5) identified in the total liver homogenate map, the top 10 positive (DA) and negative (LEW) scoring genes of this factor were selected and their enrichment in each cell within the immune-enriched map was calculated using Ucell. A) Overlying enrichment score of the top 10 DA-specific genes (varimax-5 positive loading gene) upon the immune-enriched UMAP. Cells with high enrichment of this geneset are colored dark purple and cells with zero enrichment are indicated as yellow. The distribution of the enrichment scores over the immune-enriched map clusters confirms that varimax-5 is hepatocyte-specific. B) Overlying enrichment score of the top 10 LEW-specific genes (varimax-5 negative loading gene) upon immune-enriched UMAP. The distribution of the enrichment scores over immune-enriched map clusters confirms that varimax-5 is hepatocyte-specific. C) Boxplot indicating the distribution of varimax-5's top positive gene enrichment scores within the hepatocyte population of each strain. In line with our predictions based on the total liver homogenate map, varimax-5 top positive genes are more enriched in the immune-enriched map DA hepatocyte compared to LEW (Wilcoxon-test p value <  2.2e-16). D) Boxplot indicating the distribution of varimax-5's top negative gene enrichment scores within the hepatocyte population of each strain. In line with our predictions based on the total liver homogenate map, varimax-15 top negative genes are more enriched in the immune-enriched map LEW hepatocyte compared to DA (Wilcoxon-test p value < 2.2e-16).

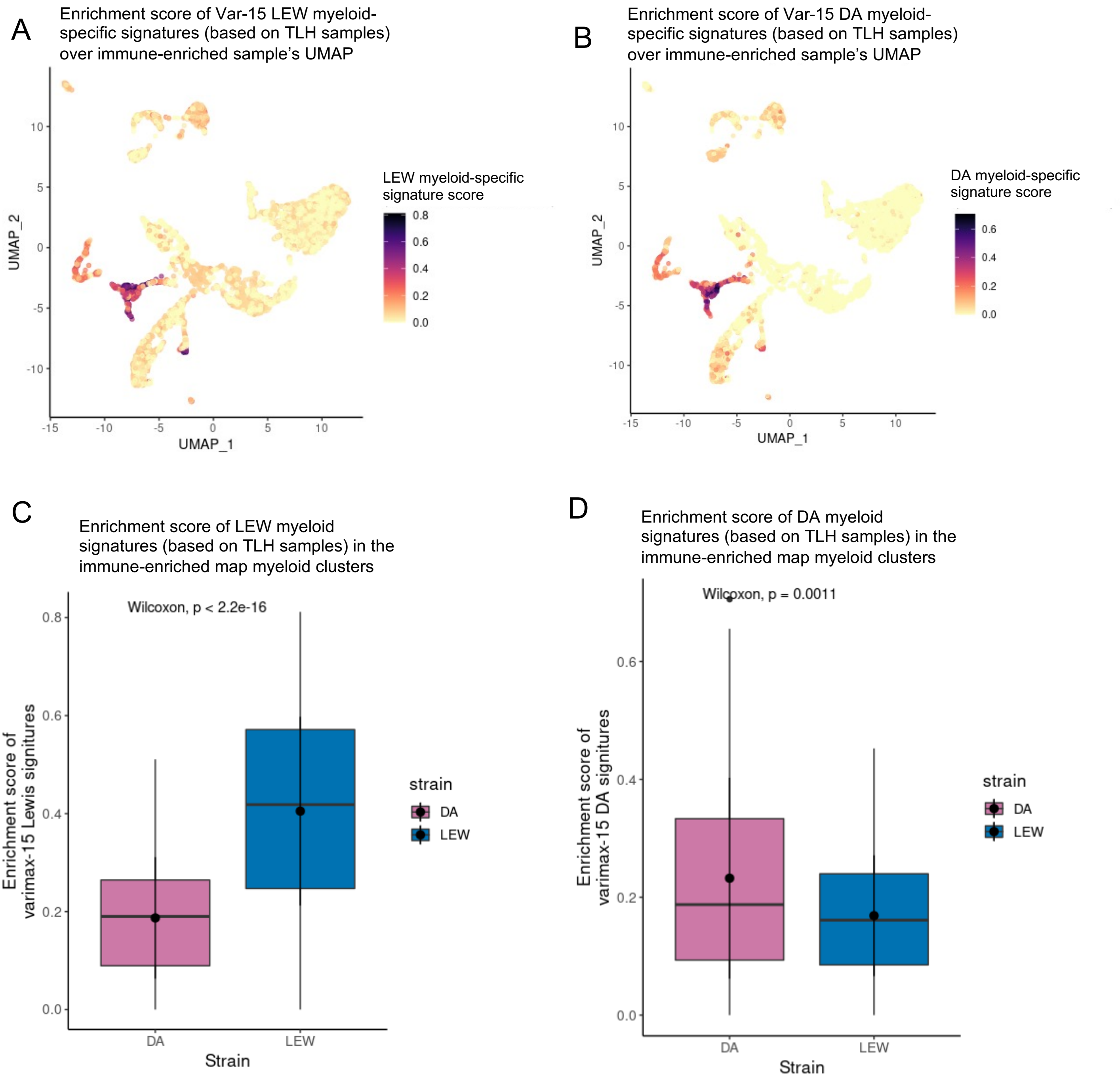
A
Enrichment score of Var-15 LEW myeloid-specific signatures (based on TLH samples) over immune-enriched sample's UMAP
LEW myeloid-specific signature score
0.8
0.6
0.4
0.2
0.0
UMAP_2
UMAP_1
10
5
0
-5
-10
-15
B
Enrichment score of Var-15 DA myeloid-specific signatures (based on TLH samples) over immune-enriched sample's UMAP
DA myeloid-specific signature score
0.6
0.4
0.2
0.0
UMAP_2
UMAP_1
C
Enrichment score of LEW myeloid signatures (based on TLH samples) in the immune-enriched map myeloid clusters
Wilcoxon, p < 2.2e-16
Enrichment score of varimax-15 Lewis signitures
0.8
0.6
0.4
0.2
0.0
DA
LEW
Strain
strain
DA
LEW
D
Enrichment score of DA myeloid signatures (based on TLH samples) in the immune-enriched map myeloid clusters
Wilcoxon, p = 0.0011
Enrichment score of varimax-15 DA signitures
0.6
0.4
0.2
0.0
DA
LEW
Strain
strain
DA
LEW

**Supplementary Figure 30) The macrophage-specific strain variations in the total liver map are verified in the immune-enriched map.** To verify the macrophage-specific strain variations (captured by varimax-15) identified in the total liver homogenate map, we selected the top 10 positive (LEW) and negative (DA) scoring genes of this factor and calculated their enrichment in each cell within the immune-enriched map using Ucell. A) Overlying enrichment score of the top 10 LEW-specific genes (varimax-15 positive loading gene) upon immune-enriched UMAP. Cells with high enrichment of this geneset are colored dark purple and cells with zero enrichment are indicated as yellow. The distribution of the enrichment scores over immune-enriched clusters confirms that varimax-15 is macrophage-specific. B) Overlying enrichment score of the top 10 DA-specific genes (varimax-15 negative loading gene) upon the immune-enriched UMAP. The distribution of the enrichment scores over immune-enriched map clusters confirms that varimax-15 is macrophage-specific. C) Boxplot indicating the distribution of varimax-15's top positive gene enrichment scores within the macrophage population of each strain. In line with our predictions based on the total liver homogenate map, varimax-15 top positive genes are more enriched in the immune-enriched map LEW macrophage compared to DA (Wilcoxon-test p value $< 2.2e-16$). D) Boxplot indicating the distribution of varimax-15's top negative gene enrichment scores within the macrophage population of each strain. In line with our predictions based on the total liver homogenate map, varimax-15 top negative genes are more enriched in the immune-enriched map DA macrophage compared to LEW (Wilcoxon-test p value $< 0.001$).

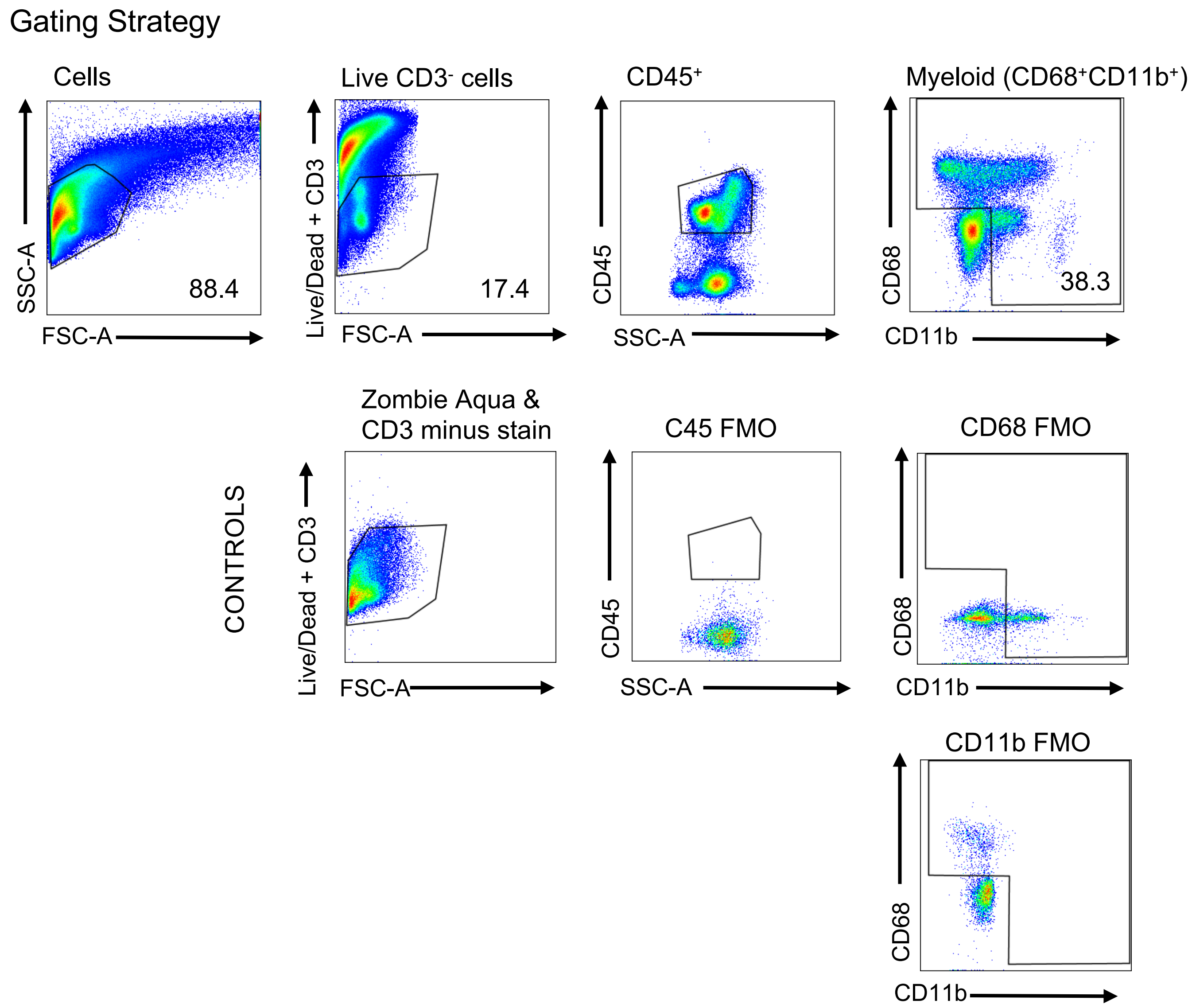


**Supplementary Figure 31) Staining controls for flow cytometry gating strategy**. The rationale for the gating strategy was based on partially stained cell suspensions. To obtain a live T cell-free population, cells stained positively for the Live/Dead Zombie Aqua dye and CD3 must be gated out. Positivity was determined via a partially stained control that consisted of the full antibody panel minus Live/Dead Zombie Aqua dye and CD3 antibody staining. For all other gating controls (CD45, CD68, CD11b), the fluorescence-minus-one (FMO) staining strategy was used.

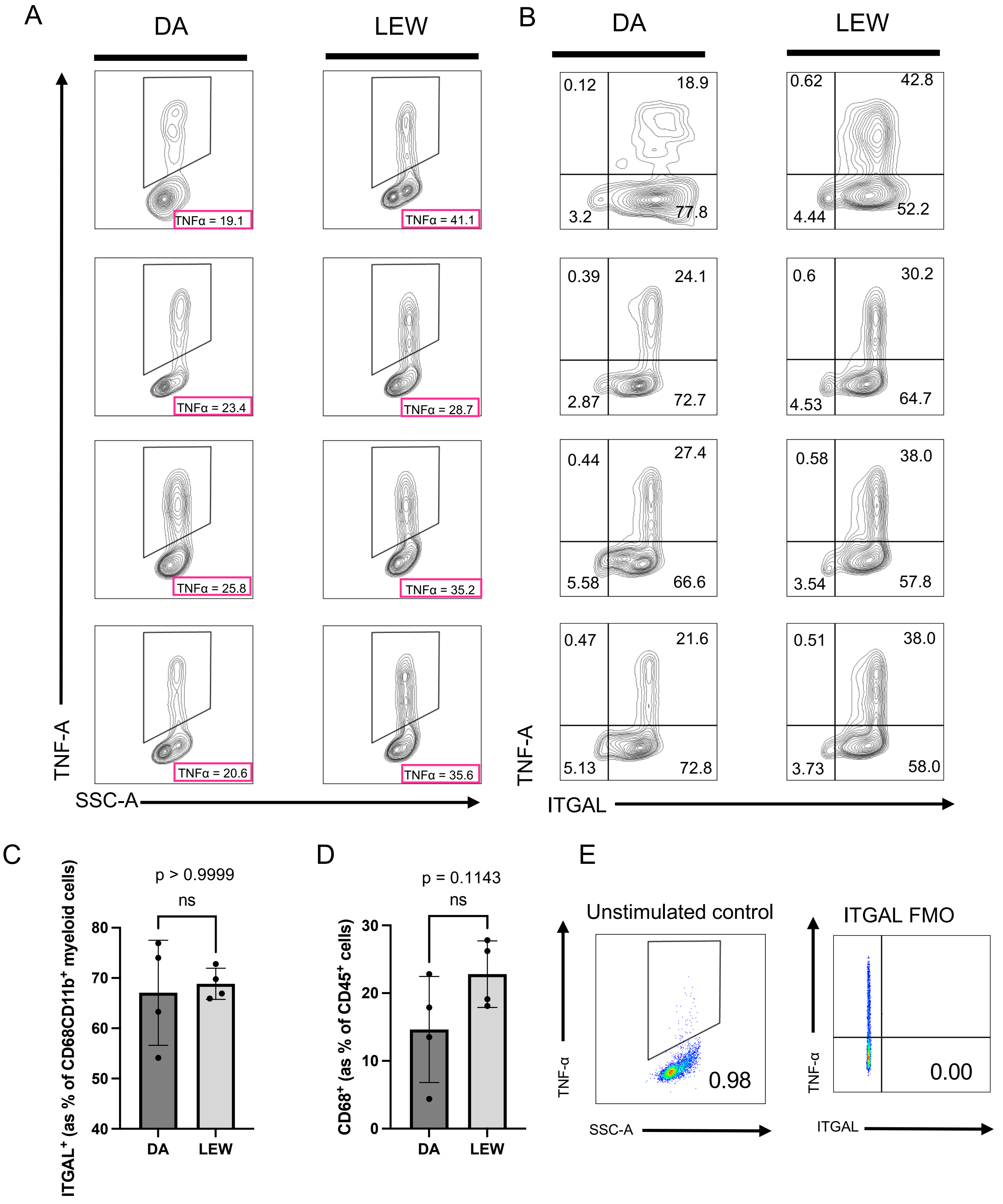

A
DA
LEW
TNFα = 19.1
TNFα = 41.1
TNFα = 23.4
TNFα = 28.7
TNFα = 25.8
TNFα = 35.2
TNFα = 20.6
TNFα = 35.6
TNF-A
SSC-A
B
DA
LEW
0.12 18.9 3.2 77.8
0.62 42.8 4.44 52.2
0.39 24.1 2.87 72.7
0.6 30.2 4.53 64.7
0.44 27.4 5.58 66.6
0.58 38.0 3.54 57.8
0.47 21.6 5.13 72.8
0.51 38.0 3.73 58.0
TNF-A
ITGAL
C
p > 0.9999
ns
ITGAL+ (as % of CD68CD11b+ myeloid cells)
80
70
60
50
40
DA
LEW
D
p = 0.1143
ns
CD68+ (as % of CD45+ cells)
30
20
10
0
DA
LEW
E
Unstimulated control
TNF-α
SSC-A
0.98
ITGAL FMO
TNF-α
ITGAL
0.00

**Supplementary Figure 32) Flow cytometry plots of all replicates of Lewis and DA intracellular cytokine stimulation assays.** ICS was performed on four animals per strain to investigate the inflammatory potentials of myeloid cells. As described in Figure 4, LPS- induced TNFα secretion was measured via intracellular cytokine staining after being treated with 1ng/mL LPS for 6 hours. Myeloid cells were gated via Live/Dead Zombie Aqua$^-$CD3$^-$CD45$^+$CD68$^+$CD11b$^+$ stains. Shown are A) the percentage of total TNFα$^+$, secreting CD68+CD11b+ myeloid cells of all LEW and DA pairs, B) TNFα secretion in ITGAL expressing CD68+CD11b+ myeloid cell subpopulations of all LEW and DA pairs, C) Summary graphs of ITGAL expressing myeloid populations of each strain. No significant differences between the strains were shown. ($p>0.9999$), D) Summary graph of CD68$^+$ myeloid cells as a percentage of CD45$^+$ cells collected at the end of intracellular cytokine assays. No significant differences between the strain were shown. ($p>0.1143$), E) controls for TNFα were based on unstimulated cells that were cultured without LPS in parallel to the LPS-treated cells whilst ITGAL staining controls were based on the FMO strategy.
